# The Genomic Ecosystem of Transposable Elements in Maize

**DOI:** 10.1101/559922

**Authors:** Michelle C. Stitzer, Sarah N. Anderson, Nathan M. Springer, Jeffrey Ross-Ibarra

**Affiliations:** Center for Population Biology and Department of Plant Sciences, University of California, 1 Shields Ave, Davis, CA 95616, USA; Department of Plant and Microbial Biology, University of Minnesota, Department of Plant and Microbial Biology, 140 Gortner Laboratory, 1479 Gortner Avenue, Saint Paul, MN 55108, USA; Genome Center, University of California, 1 Shields Ave, Davis, CA 95616, USA

**Keywords:** transposable elements, ecology, maize

## Abstract

Transposable elements (TEs) constitute the majority of flowering plant DNA, reflecting their tremendous success in subverting, avoiding, and surviving the defenses of their host genomes to ensure their selfish replication. More than 85% of the sequence of the maize genome can be ascribed to past transposition, providing a major contribution to the structure of the genome. Evidence from individual loci has informed our understanding of how transposition has shaped the genome, and a number of individual TE insertions have been causally linked to dramatic phenotypic changes. But genome-wide analyses in maize and other taxa have frequently represented TEs as a relatively homogeneous class of fragmentary relics of past transposition, obscuring their evolutionary history and interaction with their host genome. Using an updated annotation of structurally intact TEs in the maize reference genome, we investigate the family-level ecological and evolutionary dynamics of TEs in maize. Integrating a variety of data, from descriptors of individual TEs like coding capacity, expression, and methylation, as well as similar features of the sequence they inserted into, we model the relationship between these attributes of the genomic environment and the survival of TE copies and families. Our analyses reveal a diversity of ecological strategies of TE families, each representing the evolution of a distinct ecological niche allowing survival of the TE family. In contrast to the wholesale relegation of all TEs to a single category of junk DNA, these differences generate a rich ecology of the genome, suggesting families of TEs that coexist in time and space compete and cooperate with each other. We conclude that while the impact of transposition is highly family- and context-dependent, a family-level understanding of the ecology of TEs in the genome can refine our ability to predict the role of TEs in generating genetic and phenotypic diversity.

*‘Lumping our beautiful collection of transposons into a single category is a crime’*

-Michael R. Freeling, Mar. 10, 2017

## Introduction

Transposable elements (TEs) are pieces of DNA that can move from position to position in the genome. The majority of DNA in plant genomes is TE derived, and their replication and movement to new positions via transposition is the largest contributor to differences in genome size within and between taxa (Bennetzen and Kellogg, 1997). When they transpose, TEs also generate mutations as they insert into novel positions in the genome (Lisch, 2013; Oliver *et al*., 2013). These two linked processes — that of replication of the TE, and mutation suffered by the host genome — generate a conflict between individual lineages of TEs and their host genome. Individual TE lineages gain evolutionary advantage by increasing in copy number, while the host genome gains fitness if it can reduce deleterious mutations arising from transposition. As a result of this conflict, many genomes are littered with a bulk of TE-derived DNA that is often relatively transcriptionally and recombinationally inert (Fedoroff, 2012). But while this conflict between TEs and their host has long been noted to shape general patterns of TE evolution (Charlesworth and Charlesworth, 1983; Charlesworth and Langley, 1989; Kidwell and Lisch, 1997; Venner *et al*., 2009), the details of how this conflict unfolds are tenuous and rarely well understood (Linquist *et al*., 2015).

A major challenge in understanding the conflict between TEs and their host genome is simply the staggering diversity of TEs. For example, although they are united by their ability to move between positions in the host genome, the mechanisms by which TEs do so differ between the major TE classes. Class I retrotransposons, often the major contributor of TE DNA in plants (Bennetzen, 2000), can be further divided into three orders - long terminal repeat (LTR), long interspersed nuclear element (LINE), and short interspersed nuclear element (SINE). All class I TEs are transcribed to mRNA by host polymerases, some are translated to produce reverse transcriptase and other enzymes, and all use TE encoded enzymes for reverse transcription of a cDNA copy that can be integrated at a new position in the host genome. In contrast, the two major orders of class II DNA TEs transpose in different ways. TIR elements are physically excised from one position on the chromosome and moved by TE-encoded transposase proteins that recognize short, diagnostic, terminal inverted repeats (TIRs). Helitron elements transpose via a rolling circle mechanism that generates a new copy after a single strand nick by an element-encoded protein and subsequent strand invasion and repair (Thomas and Pritham, 2015). The process of transposition for most TEs (all LTR, TIR; some LINE, SINE) generates a target site duplication (TSD) in the host DNA at the integration site, and thus the identification of a TSD bordering a TE can confirm transposition. These well-described mechanisms of transposition generate predictable sequence organization that can be recognized computationally, but also generate differences in the genomic localization of these elements, via enzymatic site preference of TE encoded proteins (Labrador and Corces, 2002; Sultana *et al*., 2017).

The process of transposition generates new TE copies within a genome, forming relationships between TEs that allow their systematic grouping into families. Many taxonomic schemes for TEs exist (Finnegan, 1989; Jurka *et al*., 2005; Wicker *et al*., 2007; Kapitonov and Jurka, 2008; Piégu *et al*., 2015), but the most widely-applied approach for genomescale data (Wicker *et al*., 2007) relies on sequence homology between copies. Although not entirely representative of TE evolutionary history (Wicker *et al*., 2009; Wicker, 2012), such approaches nonetheless reflect to some degree the ability of TE encoded proteins to bind TE DNA and move other TE copies in *trans*, as recognition of specific nucleic acid sequences by TE encoded proteins is a necessary step in the transposition process. The resulting TE families thus represent groups of related TEs that share both evolutionary history and transposition machinery, and are the groupings most naturally analogous to species in higher eukaryotes.

TE families differ from one another in many ways, including their total copy number, where they insert in the genome, which tissues they are expressed in, and how they are restricted epigenetically by the host genome. In the maize genome, some families are small, found only in a few copies (e.g. *Bs*; Johns *et al*., 1985), some with tens of copies (e.g. *Ds1*; Sutton *et al*., 1984), while others contain tens of thousands of copies (e.g. *huck, cinful-zeon*; Hake and Walbot, 1980; Sanz-Alferez *et al*., 2003; Baucom *et al*., 2009; SanMiguel and Vitte, 2009; Diez *et al*., 2014). Some TE families are expressed in certain tissues, like *Misfit* in the shoot apical meristem (Vicient, 2010), while others are expressed more broadly across many (e.g. *cinful*; Vicient, 2010). Some families preferentially insert into genic regions (e.g. *Mu1*; Cresse *et al*., 1995), others in the centromere (e.g. *CRM1*; Zhong *et al*., 2002). And some families have DNA methylation across the entire body of the TE, while others lack DNA methylation, and yet others act to spread methylation outwards into flanking sequences (Eichten *et al*., 2012). In total, while it is clear that TE families differ, our understanding of their contribution to the maize genome is often studied in the context of a single family.

Although the major classes of TEs are found across taxa, their relative abundances differ (Elliott and Gregory, 2015) and there is no clear consensus as to the factors that explain the diversity of TEs within a genome (Ågren and Wright, 2011; Ågren *et al*., 2015; Sotero-Caio *et al*., 2017; Bast *et al*., 2018). One approach to understand the diversity of TEs is to consider the genome as a community and apply principles of community ecology to understand their distribution and abundance (Brookfield, 2005). Initially proposed in terms of a dichotomy between TEs that have specialized in heterochromatic or euchromatic niches (Kidwell and Lisch, 1997), thoughts about the ecology of the genome have been refined into a continuum of space, with different TE lineages existing in different genomic niches (Kidwell and Lisch, 2002; Brookfield, 2005; Venner *et al*., 2009). Empirical descriptions of TEs in a community ecology context, however, have been limited to a few families (Abrusán and Krambeck, 2006; Promislow *et al*., 1999).

Here, we take advantage of the diversity of TEs in the maize genome, the record of past transposition still detectable in the genome, and the rich developmental and tissue-specific resources of maize to investigate the family-level ecological and evolutionary dynamics of TEs in maize. We integrate many metrics that can be measured at the level of TE family to present a natural history of TEs in the B73 maize genome to characterize and describe the genomic features that differentiate superfamilies and families of TEs. We model survival of individual copies and families in the genome to facilitate an understanding of the complex and interactive strategies TEs use to associate with their host and each other, and identify suites of traits that act to define specific genomic niches and survival strategies. We conclude that understanding the diversity of TEs in the maize genome helps not only to describe TE function, but also that of the host genome.

## Methods

Scripts for generating summaries from data sources and links to summarized data are available at http://www.github.com/mcstitzer/maize_genomic_ecosystem. Interactive distributions per family can be found at https://mcstitzer.shinyapps.io/maize_te_families/.

### TE sequence properties

We base our analysis on an updated TE annotation of the maize inbred line B73 (Jiao *et al*., 2017), more fully capturing TIR elements. TEs that are nested inside of other TEs are divided for further analyses, by assigning each TE base pair in the genome to a single copy by iteratively removing copies in order of arrival. We remove from analysis any TE for which less than 50 bp remains after resolving nested copies. We add the positions of retrotransposon long terminal repeats (LTRs) to these annotations as produced by LTRharvest (Ellinghaus *et al*., 2008), and delimit the internal protein coding genes of LTR TEs using LTRdigest (Steinbiss *et al*., 2009) and GyDb 2.0 retrotransposon gene HMMs (Llorens *et al*., 2010). We additionally identify the longest open reading frame (ORF) in each TE model using transdecoder (Brian and Papanicolaou, 2018), and identify whether this longest ORF is homologous to known transposases, integrases, and replicases respectively for TIRs, nonLTR retrotransposons, and helitrons (JCVI GenProp1044 http://www.jcvi.org/cgi-bin/genome-properties/GenomePropDefinition.cgi?prop_acc=GenProp1044 and PFAM PF02689, PF14214, PF05970) using hmmscan (Eddy, 2018) with default parameters. We characterize copies as autonomous based on the content of their protein coding domains, requiring evidence of all 5 proteins (GAG, AP, RT, RNaseH, INT) for LTR retrotransposons, a reverse transcriptase match for LINEs, a transposase profile match for TIR transposons, and a Rep/Hel profile match for Helitrons. This measure is lenient in defining coding content, as it does not penalize stop codons and frameshifts throughout these coding regions.

After insertion, TE copies accumulate nucleotide substitutions that can be used to understand their age. To estimate age based on divergence of a TE copy from others in the genome, we generated phylogenies of TE copies by first aligning the entire TE sequence of each copy in each superfamily using Mafft (Katoh and Standley, 2013) (allowing sequences to be reverse complemented with the option `-adjustdirection`) and then building an unrooted tree using FastTree (Price *et al*., 2010). To make tree building computationally efficient in spite of the high number of TE copies and large element size, we use a maximum of 1000 bp for tree building for the largest 5 superfamilies (3’ terminal for Helitrons, 5’ terminal for LTR retrotransposons and TIR elements). The terminal branch length of each copy is used as a measure of its age, representing nucleotide substitutions since divergence from the closest related copy in the B73 reference genome. This measure of age makes a number of assumptions about the tempo and mode of transposition — for example, we assume nucleotide mutations in a TE arose at its current location, which may not be true for TIR elements that excise and move to a new location. Nonetheless, it is the only approach to calculate ages of individual TIR and Helitron elements (Bergman and Bensasson, 2007; Fiston-Lavier *et al*., 2012) without relying on a consensus element generated from a multiple sequence alignment that can be biased towards recently transposed copies that have not yet been removed by natural selection or genetic drift (Brookfield and Johnson, 2006; Fiston-Lavier *et al*., 2012).

Because the 5’ and 3’ LTR of LTR retrotransposons are identical upon insertion (SanMiguel *et al*., 1996), we also estimate their time since insertion using the number of substitutions that occur between the two LTRs. For each LTR retrotransposon copy, we align both LTRs with Mafft (Katoh and Standley, 2013) and calculate nucleotide divergence with a K2P correction using dna.dist in the ape package of R (Paradis and Schliep, 2018; R Core Team, 2018). For all age measures, we relate nucleotide divergence to absolute time using a mutation rate of 3.3 × 10^−8^ substitutions per site per year (Clark *et al*., 2005). These LTR-LTR estimates are generally in line with terminal branch length age estimates (Spearman’s correlation 0.65), with LTR-LTR ages often older than terminal branch length ages (Supp. Figure S6).

### TE environment and regulation

We characterize the genomic environment of the TE and features that overlap the TE. For each TE, we characterize the distance to the closest gene (gene annotation AGPv4, Zm00001d.2, Ensembl Plants v40) irrespective of strand using GenomicRanges (Lawrence *et al*., 2013). We additionally measure expression of these closest genes across a developmental atlas of the maize inbred line B73 (Walley *et al*., 2016) (accessed from MaizeGDB as walley_fpkm.txt using AGPv4 gene names). In order to estimate the overall dynamics and tissue-specificity of expression, we calculated both the median expression and *τ* (Kryuchkova-Mostacci and Robinson-Rechavi, 2016) for each of these genes. *τ* is calculated as the summed deviance of each tissue from the tissue of maximal expression, divided by total number of tissues minus 1. *τ* values thus range from 0 to 1, with low values representing constitutive expression and high values indicating tissue-specific expression.

In addition to host genes, TEs themselves can be transcribed. Using RNAseq reads from the Walley *et al*. (2016) expression atlas (NCBI SRP029238), we counted reads that align uniquely to a specific member of a TE family, as well as multiply mapped reads that align to a single family, as in Anderson *et al*. (2018). This allows estimation of the expression level of a TE family, despite the repetitive nature of TEs that limits unique mapping of reads. Reads that map to TEs located within genic sequences (generally within introns) were excluded because their expression is indistinguishable from transcription from the gene promoter. We take the mean value of reads per million across the two to three replicates per tissue, and divide by the total family size to get a per-copy metric of expression. As with genes, we calculate median expression across tissues and tissue specificity using *τ*.

To identify the recombinational environment in which each TE exists, we use a 0.2 cM genetic map of maize generated from the Nested Association Mapping (NAM) panel (Ogut *et al*., 2015). We convert AGPv2 coordinates to AGPv4 coordinates using the Ensembl variant converter (Monaco *et al*., 2014). To approximate the recombination rate in genomic regions, we fit a monotonic polynomial function to each chromosome (Murray *et al*., 2016). Using this function and TE start and end positions, we calculate a cM value for each TE, and convert to cM/Mb values by dividing by the length of the TE in megabases.

The chromatin environment a TE exists in can impact transposition (Liu *et al*., 2009). We converted data on MNase hypersensitive sites in roots and shoots (Rodgers-Melnick *et al*., 2016) from the AGPv3 reference genome to AGPv4 coordinates using the Ensembl variant converter (Monaco *et al*., 2014). We counted how many hypersensitive sites exist in each TE, as well as the proportion of base pairs of the TE that are hypersensitive. We also calculate these metrics for the 1 kb region flanking the TE on both sides.

Regulation of TEs by the host genome is often mediated via epigenetic modifications. We map bisulfite sequencing reads from shoot apical meristem, anther, ear shoot, seedling leaf, and flag leaf (Li *et al*., 2015; Eichten *et al*., 2013) using bsmap 2.7.4 with parameters (`-v 5 -r 0 -q 20`) (Xi and Li, 2009), and summarize in 100 bp windows as in Li *et al*. (2015), to characterize the local proportion of methylated cytosines in all three contexts (CG, CHG, CHH; where H is any base but G). We summarize the average levels of each measure over each TE copy and each of 20 100 bp windows of flanking sequence on either side, imputing missing data with the family mean.

To identify differences between TE copies in their base composition, we calculate GC content plus the number of di- and tri-nucleotide sites containing cytosines in a methylatable context (CG, CHG, CHH). We count these contexts in each TE using the bedtools`nuc` command (Quinlan and Hall, 2010) and divide by TE length to determine the proportion of the sequence that is methylatable for each context. We also calculate these measures of methylatability for the 1 kb flanking the TE on each side.

We also measure the number of segregating sites per TE base pair and the 1 kb flanking in the *Zea mays* Hapmap3.2.1 dataset (Bukowski *et al*., 2018) as well as the subgenome (Jiao *et al*., 2017) each TE is found within.

As we cannot calculate accurate summaries of genomic features for families with a small number of TE copies, we include only those families with more than ten copies when presenting results in the text that identify specific outlier families, such as the family with highest GC content. When presenting summaries at the superfamily and order level or results modeling TE age, we include information from all TE copies, including those from smaller families.

### Analysis and interpretation

We implement random forest regression models (in the R package ‘randomForest’ (Liaw and Wiener, 2002)) to understand the importance of different genomic features to TE survival in the genome, as measured as the age of individual extant copies. We train models on 31,000 TEs (≈ 10% of copies), and summarize 1000 iterations of trees. The remaining ≈ 280,000 TEs are retained as a test set to validate the model. Any missing data is assigned a value of -1, and the categorical variable of super-family is considered as a factor. Because of limitations to the conversion of numbers to binary, we limit categorical variable of family to the 31 largest families, and code all others as ‘smaller.’ We summarize the overall importance of each feature in predicting age by permuting its values across individual TE copies and observing the change in mean squared error of the model prediction of the actual value, scaled by its standard deviation. We summarize features into categories reflecting features specific to TE taxonomy, TE base composition, TE methylation and chromatin accessibility, TE expression, TE-encoded proteins, nearest gene expression, regional base composition, regional methylation and chromatin accessibility, and regional recombination and selection. A full description of the individual measurements that go into each category are found in Supp. Table S1.

In order to interpret family-specific relationships for top predictors of age, we perform further analyses. To further interpret these top variables, we calculate the Pearson’s correlation coefficient of each with age, using samples from each family. To visualize the nonlinear relationships and interactions produced by such models, we calculate Individual Conditional Expectations (ICE plots (Goldstein *et al*., 2015), R package ‘pdp’ (Greenwell, 2017)), which summarize the contributions of permuted values of a variable of interest to the response, while conditioning on observed values at all other variables. We provide permuted values summarizing 95% of the observed data, to provide predictions in a region of parameter space the model is trained on. We summarize these responses as deviation of the predicted value generated with permuted data from the true value, and plot as individual lines and superfamily averages.

## Results

### General features of TE orders and superfamilies

We identified members of each of the 13 superfamilies of transposable elements (TEs) previously identified in plants (Wicker *et al*., 2007) in our structural annotation of the maize B73 reference genome. This annotation resolves nested insertions of TEs within other elements, resulting in a total of 143,067 LTR retrotransposons (RLC, RLG, and RLX superfamilies), 1,640 LINE and SINE (nonLTR) retrotransposons (RIL, RIT, and RST superfamilies), 171,570 TIR transposons (DTA, DTC, DTH, DTM, DTT, and DTX superfamilies), and 22,234 Helitrons (DHH superfamily) (Table1, Figure 1A). We determined the number of families, average length, average age, distance to the nearest gene, and the number of base pairs each superfamily contributes to the genome (Figure 1; Interactive distributions per family: https://mcstitzer.shinyapps.io/maize_te_families/). For each family and superfamily, we determined the proportion of elements that are nested within another TE and the proportion of elements that are split into multiple pieces by other TE insertions.

**Fig. 1.**
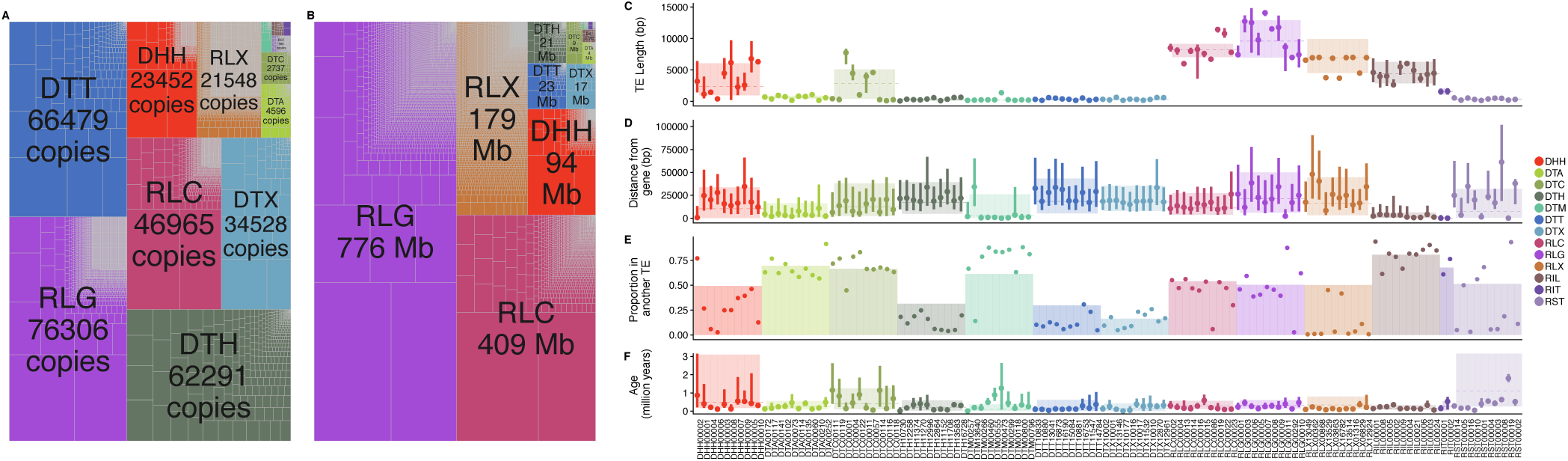
Characteristics of each superfamily of TE. (A-B) The relative copy number (A) and size in Mbp (B) of families and superfamilies shown by the size of the rectangle. Superfamilies are denoted by color, and each family is bounded by gray lines within the superfamily. (C-F) Family characteristics of each of the most numerous 10 families (≥ 10 copies) of each superfamily. (C) TE length, (D) Distance to the closest gene, (E) proportion of TE copies found within another TE, and (F) TE age. In (C, D, & F) families are shown with medians as points and lines representing ranges of upper to lower quartiles. Superfamilies are shown as colored rectangles, where the dotted line reflects the median and box boundaries reflect lower and upper quartiles. In (E), families are shown as points and superfamily proportions as a barplot.

Even at the broad taxonomic level of order, there are considerable differences among TEs. Because of their size, (median length 8.4 kb; Figure 1C, Supp. Figure S1B) LTR retrotransposons contribute more total base pairs to the genome (1,363 Mb; Figure 1B) and are commonly disrupted by another TE copy (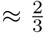 disrupted; Supp. Figure S1C). LTR retrotransposons are also typically far from genes (median distance 16.4 kb, only 3.5% within a gene transcript; Figure 1D, Supp. Figure S1A) and 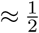 of copies insert into a preexisting TE copy (Figure 1E). Additionally, they inserted into the genome a median of 315,000 years ago (Figure 1F). In contrast, despite having more copies (Table 1), TIR elements contribute fewer base pairs to the genome (74.1 Mb) and are rarely disrupted by the insertion of another TE copy (*<* 5% disrupted) (Supp. Figure S1C), presumably due to their much smaller size (median length 306 bp; Figure 1C, Supp. Figure S1B). TIR elements as a group are also slightly further from genes (median distance 17.2 kb, 1.7% within a gene transcript; Figure 1D, Supp. Figure S1A), and commonly insert into preexisting TE copies (≈ 70% of copies; Figure 1E). They represent the most recent insertions, with a median age of 185,000 years ago (Figure 1F). And although Helitron elements are fewer in number than TIR elements, they contribute more base pairs to the genome (93.8 Mb) and are more commonly disrupted by the insertion of another TE (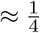 of copies; Supp. Figure S1C) due to their increased length (median length 2.4 kb). Helitrons are also closer to genes than TIR elements (median distance 10.4 kb, with 22.9% overlapping a gene transcript; Figure 1D, Supp. Figure S1A), and less frequently insert into a preexisting copy (50% of copies are found within another TE). Helitrons are represented by relatively old copies, with a median age of 500,000 years (Figure 1F). NonLTR retrotransposons (LINEs and SINEs) contribute only 2.9 Mb, are of relatively short (median length 548 bp), and only 5% of copies are disrupted by the insertion of another TE (Supp. Figure S1C). LINEs and SINEs are however often close to genes (median distance 2.3 kb, 18.6% in a gene transcript; Figure 1D, Supp. Figure S1A), and only 37% insert into another TE copy (Figure 1E). These nonLTR elements arrived in the genome a median of 350,000 years ago (Figure 1F).

**Table 1.** Superfamilies in the maize genome

| Class | Order | Superfamily | Common Name | Number Copies | Number Families |
| --- | --- | --- | --- | --- | --- |
| DNA transposon | Helitron | DHH | Helitron | 22,339 | 1,722 |
| DNA transposon | TIR | DTA | hAT | 5,096 | 275 |
| DNA transposon | TIR | DTC | CACTA | 2,768 | 73 |
| DNA transposon | TIR | DTH | Pif/Harbinger | 63,216 | 458 |
| DNA transposon | TIR | DTM | Mutator | 928 | 67 |
| DNA transposon | TIR | DTT | Tc1/Mariner | 67,533 | 269 |
| DNA transposon | TIR | DTX | Unknown TIR | 34,778 | 76 |
| Retrotransposon | LTR | RLC | Ty1/Copia | 46,553 | 2,788 |
| Retrotransposon | LTR | RLG | Ty3/Gypsy | 75,761 | 7,719 |
| Retrotransposon | LTR | RLX | Unknown LTR | 20,789 | 13,290 |
| Retrotransposon | nonLTR | RIT | RTE | 296 | 2 |
| Retrotransposon | nonLTR | RIL | L1 | 477 | 29 |
| Retrotransposon | nonLTR | RST | SINE | 892 | 533 |

But within these orders, variation also exists among superfamilies (Figure 1). For example, TE superfamilies are found nonuniformly along chromosomes (Figure 2, Supp. Figure S2): while some superfamilies like RLG (Ty3/Gypsy) and DTC (CACTA) are enriched in centromeric and pericentromeric regions, others, like RLC (Ty1/Copia) and DTA (hAT) are found more commonly on chromosome arms. As maize genes are enriched on chromosome arms, this distribution is reflected in the distance each superfamily is found from genes (Figure 1D). Similarly, while most TIR superfamilies are found far from genes (median 17.2 kb), DTM (Mutator) elements are only a median distance of 2.4 kb away from genes (Figure 1D). And although TIR elements are often short (median 311 bp), DTC elements have a median length of 2886 base pairs (Figure 1C).

**Fig. 2.**
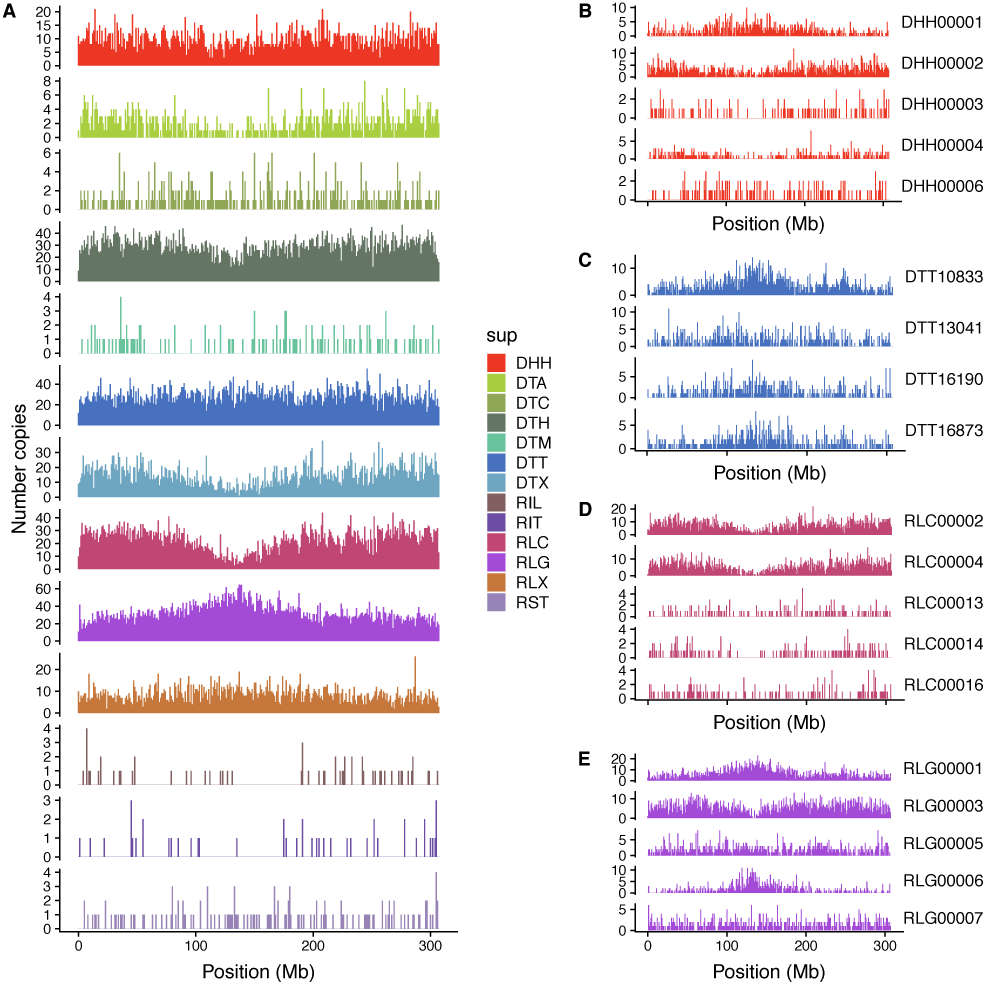
Chromosomal distribution of superfamilies and example families. Counts of number of insertions in 1 Mb bins across chromosome 1 for (A) TE superfamilies and (B-D) the 5 families with highest copy number in each of four superfamilies, DHH (B), DTT (C), RLC (D), and RLG (E).

### Features of TE families

These descriptive statistics measured at the order and superfamily level are an aggregate across many TE families. There are thousands of families of LTR retrotransposon and Helitron elements, and hundreds of families of DNA TIR elements (Table 1). And although the majority of all TE families have less than ten copies (Fig. 1A), the largest LTR retrotransposon and Helitron families in the genome contain thousands of copies. Consistent with previous analyses built on subsets of BACs (Baucom *et al*., 2009; Schnable *et al*., 2009), a majority (75%) of maize LTR retrotransposon families are present only as a single copy in the B73 genome. The average LTR family contains 6.1 copies, with this distribution ranging from 1 to 16,289 copies. In contrast, the family size distribution of TIR transposons is more uniform, with the average family containing 142 elements (range 1 to 9953) and only 10% of families represented by a single copy. Helitron families are smaller, with 14 copies on average (66% represented by a single copy), and nonLTR retrotransposon families have on average 3 copies (77% consisting of a single copy).

Families are also found nonuniformly along chromosomes (Figure 2B-E, Supp. Figure S3). Sometimes, the distribution of copies in the largest families in a superfamily match the pattern seen when summarized across all members of a superfamily, such as for RLC families which all share an enrichment on chromosome arms (Figure 2D). But there are also families that differ from the aggregate superfamily distribution. For example, the second largest RLG family (RLG00003) is enriched on chromosome arms, and the third largest RLG family (RLG00005) is more uniformly distributed along the chromosome (Figure 2E).

Further, the ages of different TE families vary greatly as well (Figure 3, Supp. Figure S4). Some families have not had a new insertion in the last 100 kya, while others have expanded rapidly in that time frame (Fig. 3B-E). Some families display cyclical dynamics, readily generating new insertions that are retained, with pulses of stasis in between (e.g. DTA00073, Figure 3C). Others show sustained activity in the past (e.g. DHH00004, Figure 3B). In total, 70% of TIR families, 20% of LTR families (estimated with LTR-LTR divergence), 15% of nonLTR families and only 7% of Helitron families have been active in the last 100 kya.

**Fig. 3.**
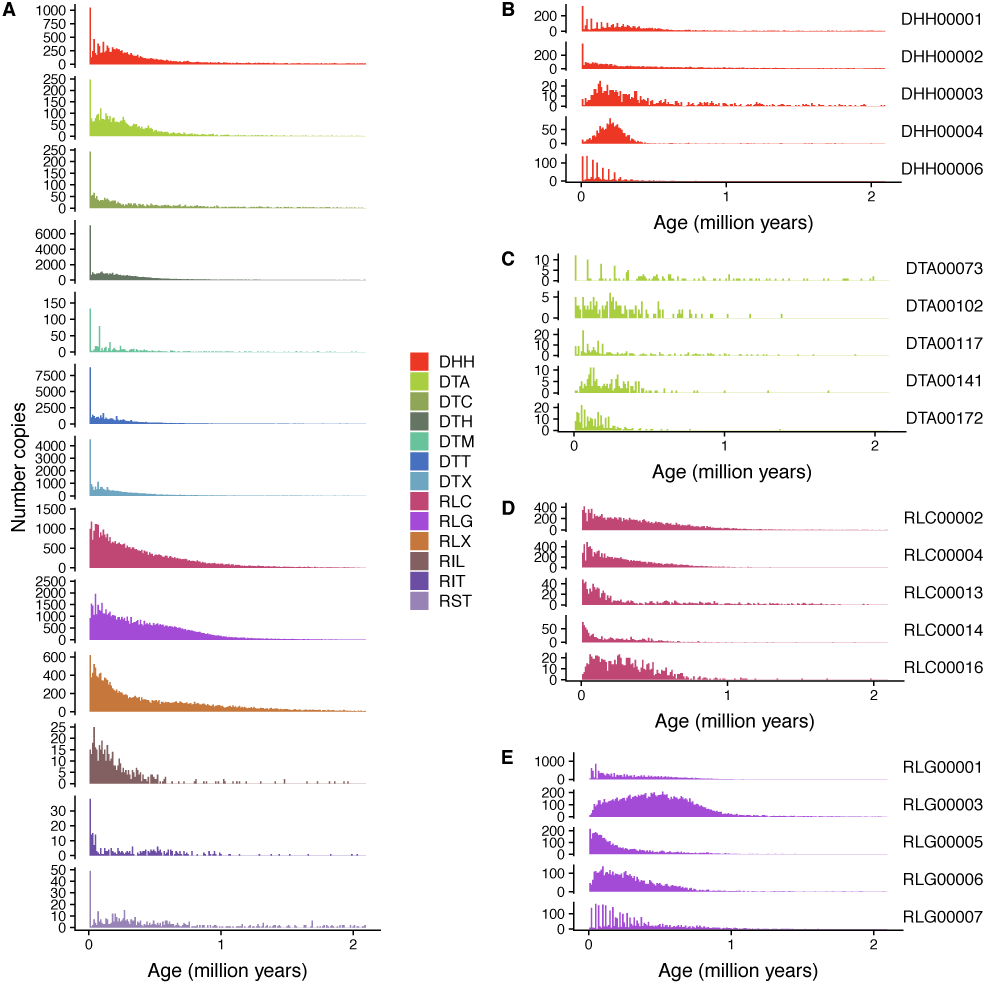
Age distribution of (A) superfamilies and (B-E) five largest families of (B) DHH, (C) DTA, (D) RLC, and (E) RLG. Relative abundance within a family are shown in (B-E). Counts of number of insertions in 10,000 year bins are shown. As they are rare, TE copies older than 2.1 million years are not shown. Ages are calculated with terminal branch lengths for all TEs except LTR retrotransposons, which are calculated with LTR-LTR divergence. See Supp. Figure S5 for LTR retrotransposon plots with terminal branch length ages.

### Features of the transposition process

As families arise via transposition to new positions, we address different features that restrict and allow movement of TE copies.

#### TE proteins

Numerous sequence features of the TE itself are required for the complex transposition process to occur, which is best understood at the level of TE family. One requirement is the presence of TE encoded proteins that catalyze movement. Functional characterization of TE protein coding capacity is complicated by difficulty in identifying the effect of stop codons or nonsynonymous changes on transposition — instead we measure homology to TE proteins, which may not fully reflect whether a TE copy can produce a transpositionally-competent protein product. Although TE-encoded proteins are often of similar length within a TE superfamily due to domain conservation and shared ancestry, the longest ORF in a TE varies by family (Figure 4A). Sometimes this is due to the presence of nonautonomous or non-coding copies. While nonautonomous copies rely on protein production in *trans* by other family members, autonomous TE copies encode their own transposition machinery in *cis*. 52% of LTR families, 0.6% of TIR families, 0.3% of helitron families, and 0.2% of nonLTR families have at least one member that retains some remnant of coding capacity for all the TE proteins necessary for transposition, with substantial variation within families (Figure 4B, Supp. Fig. S11A-G). Several LTR retrotransposon families have a small proportion of autonomous copies (Figure 4C), and yet other families partition coding potential for required proteins between different TE copies (e.g. RLG00001, where only 0.3% of copies code exclusively for GAG and 12.1% of copies code for only POL, although both proteins are required for retro-transposition; Figure 4C). Also, families range from having almost exclusively autonomous copies (14 families of DTC, RLC, and RLG have at least 75% of copies in the family carrying coding capacity, Supp. Table S2), to having exclusively nonautonomous copies (842 families, spanning all 13 super-families) (Supp. Table S3).

**Fig. 4.**
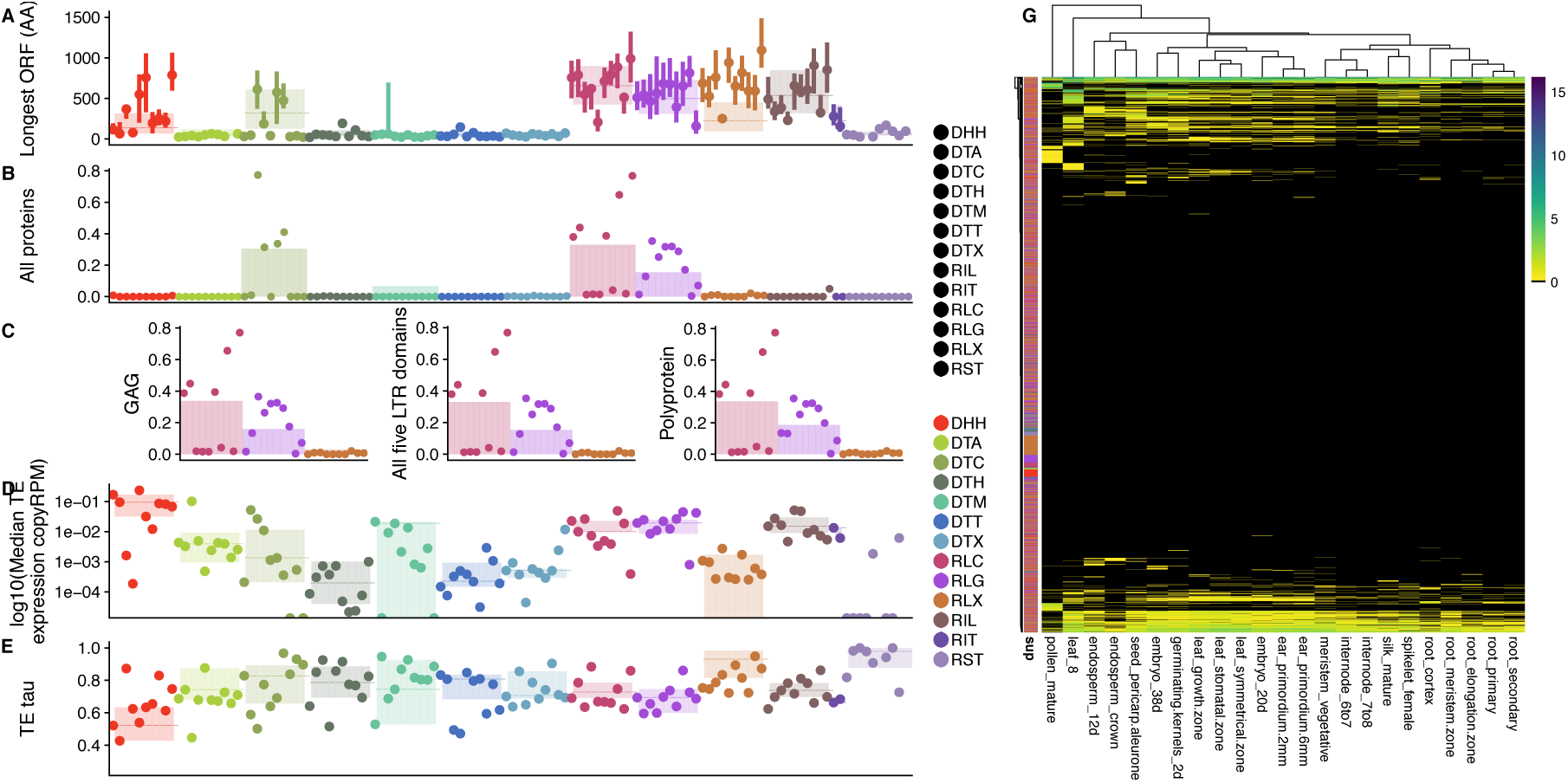
TEs code for proteins that are expressed, and expression varies by family across tissues. In A-F, families are in the same order as presented in Figure 1. (A) Length of longest open reading frame within the TE, measured in amino acids. (B) Presence of all proteins required for transposition. (C) Presence of GAG, all five domains (GAG and Pol), and Pol (which encodes four domains) in LTR retrotransposons. (D) log10 median TE expression across tissues, per-TE copy. (E) Tissue specificity of TE expression *τ·*, with low values representing constitutive expression, and high values representing tissue specificity. (F) Per copy TE expression across tissues (RPM), clustered by expression level. Families with greater than 10 copies in rows, tissues in columns.

Coding capacity for TE proteins likely dictates the ability to generate new insertions, and as such is associated with TE age. This is not always mediated by the age of a specific TE, but whether a family member exists that codes for protein. Averaged across all orders, TEs that code for proteins are younger than their family members that do not code for proteins (median age of 198 kya vs. 285 kya), and families that lack a coding member in B73 show an intermediate median age (243 kya). But this pattern holds across only a few superfamilies (DTC, DTX, RLC, and RLG), and instead, for most superfamilies, coding members are older than noncoding copies from coding families (Supp. Figure S7).

#### TE expression

Beyond simply coding for TE proteins, another requirement for TE transposition and transgenerational inheritance is expression of the TE itself, such that the TE-encoded protein can be generated. Mapping of RNA-seq reads to repetitive TE families is a challenge, as it can be impossible to identify the exact copy that is expressed when a read maps equally well to multiple TE copies (Slotkin, 2018). We choose to summarize multiply mapping reads and TE expression at the level of per-copy RPM of the family, which likely averages relevant variation in expression known to exist within maize TE families (Anderson *et al*., 2018). Large families are generally transcriptionally repressed, while small families show higher median per-copy expression levels. While superfamily medians and median expression per copy of the ten largest families per superfamily show below 0.1 RPM per copy (Figure 4D), per copy rates of expression can be higher for small families. For example, the 19 copies of RLC00184 (also known as *stonor*) show high median expression of 4.33 RPM per copy. Tissue specificity can reflect different strategies for TE survival, like that a TE must jump in germline tissue to ensure its transgenerational inheritance at a new locus. Tissue specificity is highest when values of *τ* are equal to 1, and 0 when constitutively expressed at identical levels across all tissues. Helitrons and most LTR retrotransposon superfamilies (RLC and RLG) show lower *τ* than TIR and nonLTR retrotransposon superfamilies (Figure 4E). Tissue specificity can be extreme, with some families showing expression in only one tissue (Figure 4E)). For example, DTH00434 shows maximal per copy expression in mature pollen (4.3 RPM), with highly tissue specific expression (*τ* =0.998).

#### TE regulation

TE expression is likely limited by regulation of the TE by the host genome, which we measure via DNA methylation and MNase hypersensitivity in the TE and regions surrounding it. TEs on average are heavily regulated by their host genome: average cytosine methylation across structurally intact TEs is high (averaged across five tissues, 82% of cytosines in a CG context in a TE are methylated, 67% in a CHG context, and 4% in a CHH context), although this varies across superfamilies (Supp. Table S4) and families (Figure 5A,C,E).Only a small fraction of base pairs within TEs is in chromatin accessible to MNase, only 0.2% in shoot tissue, and 0.08% in root tissue (Figure S9E,G), both lower than genome-wide proportions (0.5% in shoot, 0.2% in root).

**Fig. 5.**
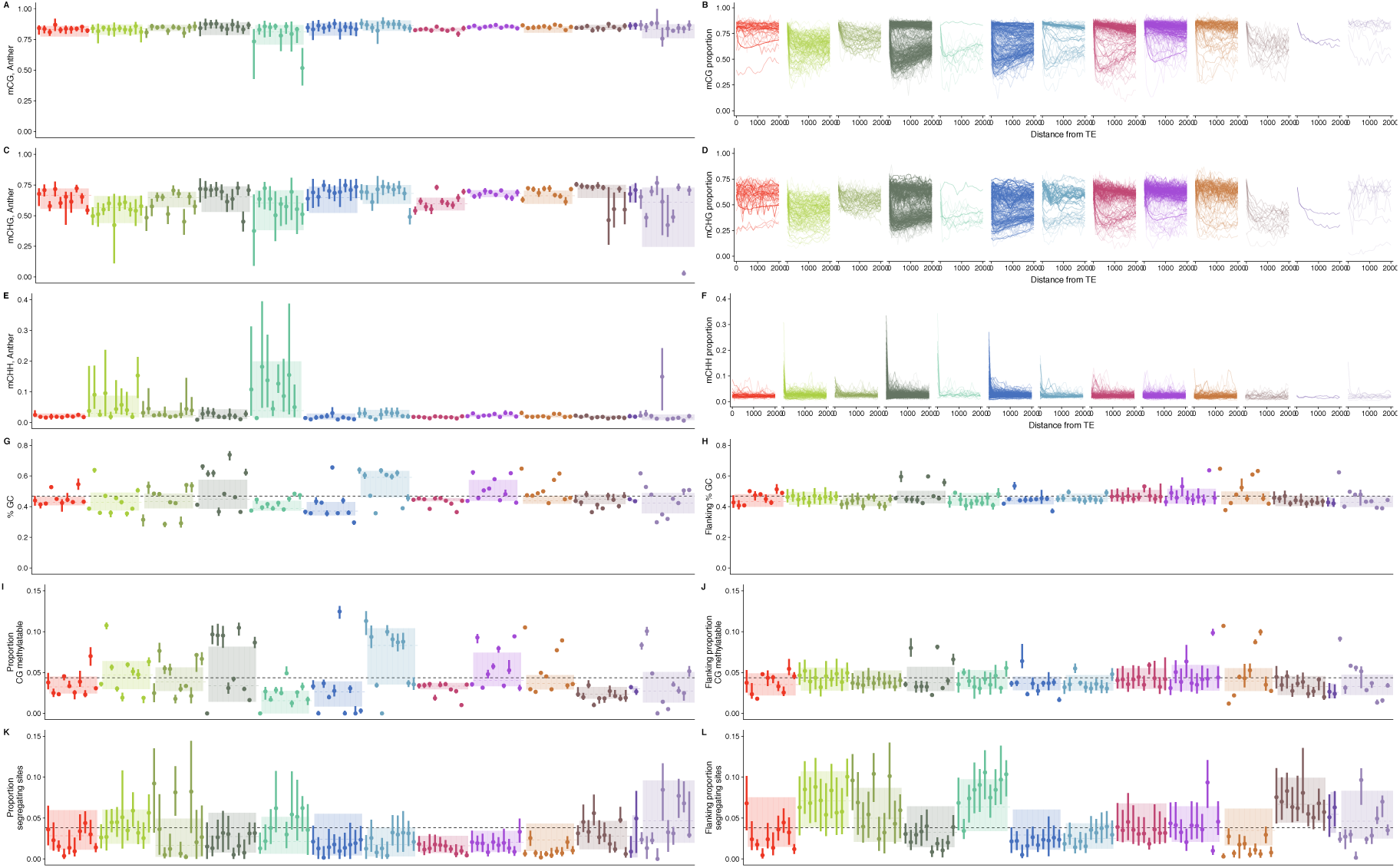
TEs and their flanking sequences are regulated by their host genome. Families are presented in the same order as in Figure 1. CG methylation in TE (A) and 2 kb flank (B), CHG methylation in TE (C) and 2 kb flank (D), and CHH methylation in TE (E) and 2kb flank (F). All methylation data from anther tissue, other tissues shown in Supp. Figure S8. GC content in TE (G) and 1 kb flank (H), percent CG methylatable base pairs in TE (I) and 1 kb flank (J), proportion segregating sites in TE (K) and 1 kb flank (L). In A,C,E, G, H, I, J, K, and L, superfamily median shown as a dashed line with the interquartile range in the shaded box. Each of the ten largest families with ten copies are shown with points denoting medians, and lines denoting the interquartile ranges. Genome-wide values for each measure are shown as gray dashed lines. In B,D, and F, median methylation for regions up to 2 kb up and downstream of the TE are plotted for each family, with family size denoted by line transparency (darker lines are larger families).

But despite this overall pattern of regulation, the host genome restricts some families of TEs differently. For example, the median CG methylation of the family DTM00796 is only 52% in seedling leaf tissue (Figure 5A), despite most other families showing higher methylation. There is even more extreme variation in CHG methylation across TE families (Figure 5C), and while many TE families show low CHH methylation across the body of the TE, some families of DNA transposons, largely in the superfamilies DTA and DTM, show relatively high CHH methylation (Figure 5E). Although the numbers presented here are for anther tissue, these patterns are robust across tissues (Supp. Figure S8).

Methylation levels in the region surrounding a TE insertion can remain similar to that of the TE, or decay to background genomic levels. Of the 1,243 TE families with ten or more copies, 734 TE families have elevated CG methylation within the TE compared to 500 bp away, 957 show elevated CHG methylation, and 1086 families show elevated CHH methylation, when median methylation levels are averaged across all tissues. This pattern can be visualized as the decay of methylation moving away from the TE for CG and CHG methylation (Figure 5B,D,F).The magnitude of reduction in local methylation moving away from the TE differs in extent and pattern, including families where methylation is reduced immediately adjacent to the TE, and others with minimal reductions even 2 kb away from the TE (Figure 5B,D). In contrast, most families show rapid reductions in CHH methylation within 100 bp away from the edge of the TE (Figure 5F).

#### TE base composition

Observed DNA methylation levels may be impacted by the base composition of the TE, as cytosines must be present to be methylated. TE families differ in GC content (Figure 5G); with extremes ranging from 21% (DTT13542) to 84% (DTH14236) median GC content. This appears to be a consequence of bases carried by the TE itself and not of regional mutation pressure, as variation in GC content in the TE is greater than that of the flanking sequence (Figure 5H). For example, GC content in the 1kb flanking DTH14236 is over 30% lower than that in the TE (52% GC in the flanking region). But beyond the proportion of cytosines in the sequence, the context in which these cytosines are found can impact whether and how they are methylated. For example, 51 families have a median of 0 cytosines that can be methylated in either the CG or CHG context (Supp. Table S5). And even with similar GC content, families differ in the contexts in which they have those cytosines, as families can have moderate GC proportions, but high proportions of these in a CG context (e.g. DTM00473; Figure 5G,I). Notably, these enriched proportions of methylatable cytosines we observe within the TE do not exist for the region flanking the TE (Figure 5H,J).

Although difficulty in mapping short reads to a highly repetitive genome precludes a comprehensive analysis of population frequencies of TEs across maize individuals, we use as a proxy for copy number the proportion of segregating sites within TEs in maize HapMap3 individuals (Bukowski *et al*., 2018), a panel that includes 1,218 maize and teosinte individuals. While as a whole TEs have fewer segregating sites per base pair (median 0.022) than the genome-wide proportion (0.0395) (Figure 5K), some TE families show high numbers of segregating sites (e.g. DTH10060, 0.177 segre gating sites per bp), suggesting differences in copy number and mutational pressures in divergent germplasm may have generated these differences. In contrast to the sequence carried by the TE, variation in the region the TE is inserted into is considerably closer to genome-wide averages than that of the TE itself (median 0.034 segregating sites per bp; Figure 5L).

#### Features structuring TE survival after insertion

The recombinational environment that a TE exists in can impact the efficacy of natural selection on the TE, as higher recombination can unlink deleterous variation from adaptive mutations (Hill and Robertson, 1966), leading to a positive relationship between recombination and diversity. While LTR retrotransposons are more commonly found in low recombination regions (median 0.30 cM/Mb), Helitrons and TIR elements are more commonly found in higher recombination regions (both show a median 0.43 cM/Mb), and nonLTR retrotransposons are found in the highest recombination regions (median 0.57 cM/Mb). But this varies by family, and for example the two largest families of DTT differ in median recombination regions from 0.14 cM/Mb to 0.53 cM/Mb (Supp. Figure S10A).

Additionally, selection can act on TEs if they have an impact on the expression of genes they land near. Although it is impossible to determine whether a TE insertion causes changes in nearby gene expression using only the B73 genome, we observe differences in the expression levels of the genes closest to superfamilies and families of TEs. Across tissues, genes near TIR and nonLTR elements have higher median expression (1.37 RPKM for TIR and 1.83 RPKM for nonLTR) than genes near LTR (1.04 RPKM) and Helitrons (0 RPKM) (Supp. Figure S10C). Notably, this pattern intensifies for genes within 1 kb of the TE, where median median gene expression is over 4 RPKM for genes near TIR and nonLTR elements, but 0 RPKM for these genes close to LTR and Helitron elements (Supp. Figure S10D). Some families are often found near highly expressed genes (e.g. DTA00133, median expression 22.38 RPKM), while 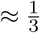 of families are closest to genes that are not expressed. However, when genes near TEs are expressed, their expression is much more constitutive than that of TE families (Supp. Figure S10E, Figure 4E), with mean *τ* values of 0.75 for genes near TEs and 0.93 for TE families themselves. Tissue specificity varies by family and superfamily as well, and there is a weak correlation between tissue specificity of expression of TE families and expression of the genes they are closest to (Pearson’s correlation 0.067, p=4e-12).

The maize genome arose from an autopolyploidy event (Swigoňová *et al*., 2004), and has been sorted into two extant subgenomes (Schnable *et al*., 2011). Subgenome A has retained more genes and base pairs than subgenome B, accounting for 64.8% of sequence (Jiao *et al*., 2017), and 64% of all TEs (Supp. Figure S10B). Consistent with weaker purifying selection and less conservation in subgenome B (Schnable *et al*., 2011; Pophaly and Tellier, 2015), the median age of TEs in subgenome B is slightly lower (0.24 Mya) than those in subgenome A (0.26 Mya). The lack of subgenome differentiation in TE distributions is likely due to the effect of ongoing transposition erasing any signature of TE differences between parents of the allopolyploidy event, as genome-wide the family with the oldest median age (DTH16531) is only 8.5 million years old.

### Modeling survival of TEs

To account for the myriad differences of these 341,426 TE copies in 27,444 families, we approach our understanding of the survival of TEs in the genome by modeling age as a response to these TE-level features and the genomic regions in which TEs exist today. Age reflects survival of TEs, measuring the amount of time since transposition that they have persisted at a genomic position, not being lost by selection or drift. Hence, we measure the predictive ability of features of the TE itself and the genomic region it inserted into on TE survival as measured by age.

Although relative age differences between TE insertions are limited only by our ability to count mutations, absolute age estimates can be shifted by mutation rate estimates. We use a maize-specific mutation rate (Clark *et al*., 2005), which explains our younger age of maize LTR retrotransposons than the 3-6 million years originally defined by SanMiguel *et al*. (1998). Additionally, as nucleotide mutation rates in TEs may be higher than other parts of the genome (≈2 fold higher in TEs in *Arabidopsis thaliana*, (Weng *et al*., 2018)), we consider our age estimates to represent an upper bound of TE age. Nonetheless, age represents a comparable metric of survival in the genome, especially when summarized across multiple copies and families. Furthermore, as the choice of mutation rate shifts only absolute age in years, age is only linearly rescaled, not changing qualitative results.

Random forest regressions using age as a response variable and features that are measured at the level of the individual TE explain moderate amounts of variance (23.6%), and show low mean squared error (0.014). Across all TEs, information on the superfamily a TE belongs to contributes the most to prediction accuracy for age; after permuting their values, the square root of mean squared error (RMSE) increases by 123 kya (Figure 6B). Other features that increase RMSE by over 100 kya include the number of segregating sites per bp within the TE, the size of the family it comes from, and the extent of the TE in base pairs along the genome (including bases coming from copies nested within it). In aggregate, features of the region flanking each TE explains approximately as much variation in age as features of the TE itself, but there are more flanking features than those measured on the TE. On average, each feature of the TE contributes over 3 times more predictive power than that of a flanking feature (square root mean squared error of 31 kya for a TE feature, 7 kya for a flanking feature) (Figure 6A, Supp. Table S1).

**Fig. 6.**
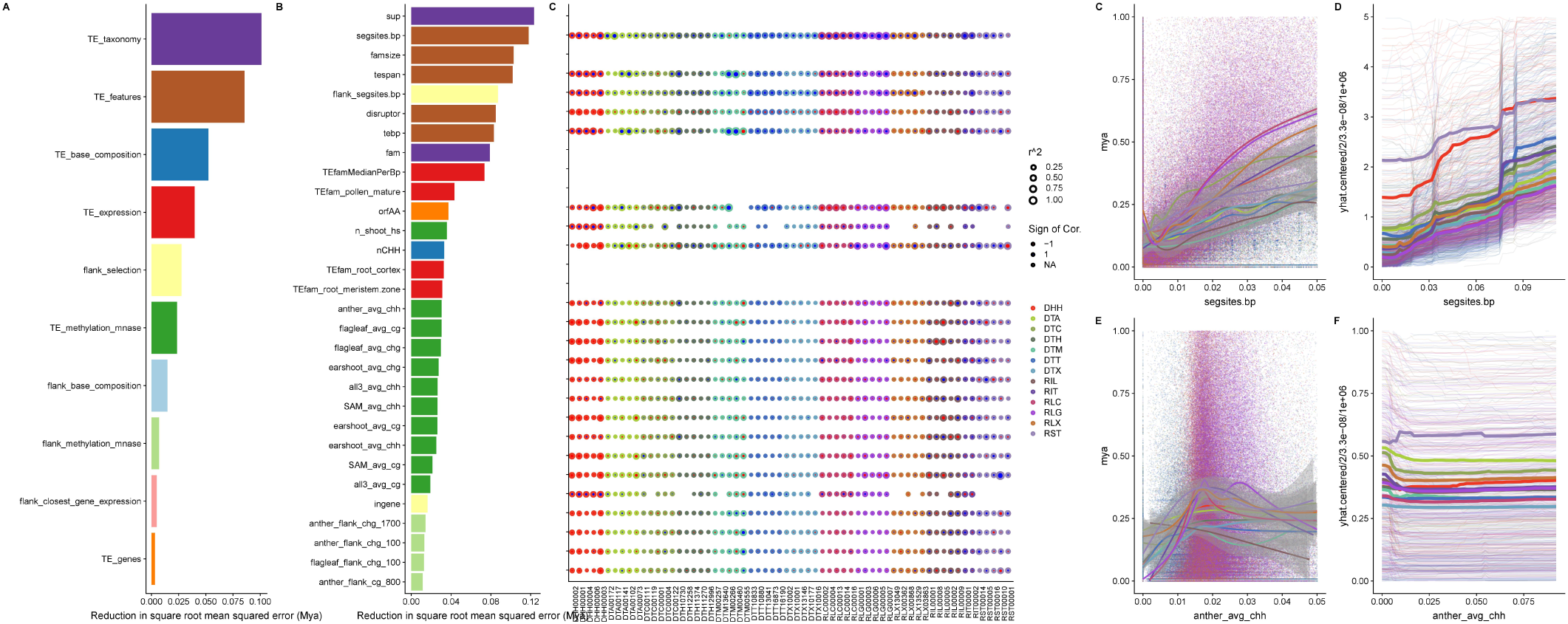
Features ranked by importance. (A) Reduction in mean squared error gained by including a feature in a model, summarized into categories. (B) Reduction in mean squared error for top 30 individual features. (C) Correlations of each of the top 30 features with age for the five largest families in each superfamily. Size of point is scaled by correlation coefficient, and color by whether the relationship is positive or negative. Rows without values are features that are fixed within a family, thus have no variance. (D) Raw correlations between age and segregating sites per base pair (E) Model predictions for the relationship between age and segregating sites per base pair (F) Raw correlations between age and anther CHH methylation of the TE (G) Model predictions for the relationship between age and anther CHH methylation of the TE.

These generalities reflect underlying nonlinearities in the relationships between individual features and age, which are often family-specific. Indeed, correlations of these top features with age differ not only in magnitude, but even in sign within individual families (Figure 6C). We use the fitted random forest models to predict age for TE copies as we vary a feature of interest. This allows insight into the local behavior of the model, to determine the relationship between a feature of interest and age. To investigate the relationship between the number of segregating sites in the TE and its flanking sequence, we predict age values for TEs with perturbed segregating site values. We find a positive relationship, consistent with patterns observed in the raw data (Figure 6D), where higher values of segregating sites within the TE across maize lines predict older TEs (Figure 6E). Despite this overall pattern, individual families vary in the slope and even sign of the relationship (Figure 6D). Other features, like CHH methylation of the TE in anther tissue, show relationships that vary by superfamily, where RIT and DHH appear older with increasing CHH methylation of the TE in the anther, while other superfamilies show decreasing age. The interactions with different variables likely give rise to these differences between raw values and adjusted values. In total, these important genomic and TE features contribute to prediction of age, but interactions with other features dominate their interpretation.

## Discussion

### General Patterns

As 85% of the maize genome is repetitive sequence (Schnable *et al*., 2009; Baucom *et al*., 2009), and 63% structurally recognizable TE sequence (Jiao *et al*., 2017), TEs contribute more to the maize genome than sequence that is uniquely ‘maize.’ Like most plant genomes (Bennetzen, 2000), retrotransposons contribute more base pairs to the maize genome than do DNA transposons (Table 1, Figure 1B). This is a consequence of the high number of copies (Figure 1A) and the large size of individual retrotransposons (Figure 1C), likely due to a ‘copy and paste’ replication mode that leaves existing copies intact when generating new copies. And as like other plant genomes (Bennetzen *et al*., 2005; Han *et al*., 2013), most superfamilies of DNA transposon in the maize genome are found closer to genes than are retrotransposons (Figure 1D). This is likely due to targeted insertion into euchromatic sequences (Jiang and Wessler, 2001; Liu *et al*., 2009), and differences in removal through natural selection after insertion (Wright *et al*., 2001; Tenaillon *et al*., 2010).

### TE superfamilies

But the bulk of TE sequence is often described at a finer scale, that of individual superfamilies of TEs. Each TE superfamily defined in the maize genome has representatives across the tree of life (Eickbush and Jamburuthugoda, 2008; Yuan and Wessler, 2011; Kapitonov and Jurka, 2007), suggesting an ancient origin of these genomic parasites. And some superfamilies have retained dramatic and consistent differences in their spatial patterning across the genome across hundreds of millions of years of radiations of plants and animals. For example, the superfamily RLG is enriched near centromeres in all plants (Du *et al*., 2010; Neumann *et al*., 2011; Slotkin *et al*., 2012) including maize (Figure 2A), highlighting a genomic niche that allows long-term survival near the centromere. Similar patterns exist at deep time scales for DNA transposon superfamilies, which preferentially insert near genes in both monocots and dicots (Bureau and Wessler, 1994) and in maize are enriched on chromosome arms where genes are concentrated (Figure 2A).

These patterns are a result of the evolution of different ecological strategies of TEs in the genome. Kidwell and Lisch (1997) described two extremes to the ‘ecology of the genome’ — one, a TE that preferentially inserts far from genes, into low recombination heterochromatic regions, and a second, risky TE that inserts near low copy sequences, more likely to disrupt gene function. These extremes are certainly at play in the maize genome. LTR retrotransposons dominate the heterochromatic space, with over half of all copies greater than 16 kb from a gene (Figure 1C), and most copies are heavily methylated (Figure 5A,C,E). The alternate strategy also exists in the maize genome, with risky insertions near genes and transcribed regions seen for several TIR superfamilies, like DTM, where over half of copies are found within 1 kb of a gene (and over one quarter of DTM within 100 bp of a gene) (Figure 1C).

### TE Families

While superfamily level observations are useful for gaining an overview of the distribution and survival of TEs in a genome, more detailed study on a time scale relevant to the evolution of the genus *Zea* comes from studying TE families. Maize TE families are shared with closely related host species, but the number of shared families rapidly decreases with phylogenetic distance. Many families are shared with congeners *Zea diploperennis* (Zhang *et al*., 2000; Meyers *et al*., 2001; Estep *et al*., 2013) and *Zea luxurians* (Tenaillon *et al*., 2011), but few families investigated are found in maize’s sister genus *Tripsacum* (1 mya divergence; (Ross-Ibarra *et al*., 2009)) (Zhang *et al*., 2000; Meyers *et al*., 2001; Gerlach *et al*., 1987; Purugganan and Wessler, 1994; Estep *et al*., 2013), and the only families shared between maize and *Sorghum* (12 mya; (Swigoňová *et al*., 2004)) are shared only as a result of horizontal transfer events between the species (Roulin *et al*., 2009). This suggests that in order to understand TE evolution at a timescale relevant to maize as a species, it is essential to understand families of TEs, rather than the aggregate properties of superfamilies or orders.

Indeed, family-level analysis can also reveal patterns obscured when averaged together at the level of superfamily. For example, despite the fact that the RLG superfamily is enriched in centromeric and pericentromeric domains (Figure 2A), the second largest family RLG00003 (homologous to huck) is predominantly found on chromosome arms (Figure 2D). While many RLG elements contain a chromodomain targeting domain in their polyprotein (Malik and Eickbush, 1999) allowing targeted insertion to centromeres, RLG00003 does not (Supp. Figure S11G). This lack of a chromodomain may explain a proximal cause of this differing ecological niche of RLG00003, although other factors are certainly at play, as other families with centromeric enrichment also lack chromodomains (Supp. Figure S11G). DNA transposons are also best described at the family level. While Mutator (DTM) elements are found a median distance of 2.5 kb from genes (Figure 1D) and have long been observed to target insertions near genes in maize (Cresse *et al*., 1995; Liu *et al*., 2009; Jiang *et al*., 2011), the second largest family, DTM13640, is found a median distance of 34 kb away from genes (Figure 1D). The mechanism for gene targeting seems to be mediated through recognition of open chromatin (Singer *et al*., 2001; Liu *et al*., 2009), but precise details of the targeting are unknown and further investigation into the families that insert near and far from genes may pinpoint how their molecular mechanisms of targeting may differ.

Furthermore, differences in the timing of transpositional activity vary extensively between families. Most TE families in maize have had most new insertions in the last 1 million years (Figure 3). But some TE families have bursts of activity, punctuated by a lack of surviving new insertions, while others appear to be headed towards extinction. All of these timings are much more recent than allopolyploidy in maize (≈12 mya) (Swigoňová *et al*., 2004) and show no subgenome bias in their distribution, suggesting that these represent lineages evolving within maize.

Maize was domesticated from teosinte (*Zea mays* subsp. *parviglumis*) 9,000 years ago (Matsuoka *et al*., 2002; Piperno *et al*., 2009). It is tempting to address the contribution of TEs to this major transition, especially given numerous cases of insertional mutations in domestication genes (Studer *et al*., 2011; Yang *et al*., 2013; Huang *et al*., 2018). Although we caution that mutation rates and estimation can complicate ascertainment (see below), 46,949 TEs across all 13 superfamilies have an estimated age of less than 9,000 years, and 24,630 TEs have an estimated age of 0. This suggests that transposition has been ongoing since the divergence of maize from its wild ancestor, but we caution that we lack appropriate confidence intervals for these estimates, especially as non-zero age requires observing at least one mutation.

### The Family-level Ecology of the Genome

It can be difficult to predict exactly why a particular TE family differs from other families. Community ecologists aim to understand the environmental factors that give rise to the observed diversity of organisms living in one place, including not just features of the environment, but also interactions between species. The species of the genomic ecosystem are families of TEs, and because the genomic environment a TE experiences is constrained to the cell, TEs are forced to interact in both time (Figure 3) and space (Figure 2). We predict each family of TE is adapted to its genomic ecological niche, where the genomic features we measure represent the environmental conditions and resources limiting a species’ ecological niche (Hutchinson, 1957).

In the genomic ecosystem, we can observe interactions between species much like we would see in a traditional ecosystem. We see a number of patterns, including cyclical dynamics of TE activity through time for several families, sustained activity through time, and a reduction in new copies towards the present (Figure 3, Supp. Figure S4). This means that the genomic environment a newly arriving TE experiences is dependent on the activity and abundance of all other TE families in the genome. At one extreme, members of the same family can even partition protein coding domains between different pools of TE copies, leading to a symbiotic mutualism where both copies are required to be transcribed and translated for either to transpose. Previous knowledge of these systems was limited to the maize retrotransposon families Cinful, which codes for polyprotein domains, and Zeon, which codes for GAG (Sanz-Alferez *et al*., 2003) (represented here by a single family, RLG00001). This strategy has been successful in maize, and RLG00001 (Jiao *et al*., 2017; Estep *et al*., 2013) for example makes up 135Mb of sequence. *Sorghum*, in contrast, has a genome 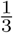 the size of maize (Paterson *et al*., 2009) and lacks homologs to RLG00001. Such symbiotic relationships within a TE family have been thought of as remarkably rare (Le Rouzic *et al*., 2007; Wagner, 2006); however we identify 25 LTR retrotransposon families where GAG and POL protein domains are found in separate TE copies but less than 1% of copies contain both, suggesting that this pattern is much more prevalent than previously described. These types of elements are best classified as sub-types of a single family, because the *cis* components of the LTR are recognized by protein domains of both GAG and POL proteins, leading to homogenization of sequence signals. As noted by Le Rouzic *et al*. (2007), symbiotic TE families face a major barrier in being horizontally transferred, as both copies must be transmitted through an already rare process. Their prevalence in the maize genome thus supports instead a long term coevolution of the maize genome and the TEs that live within it, specializing and diversifying with different ecological strategies. In total, the genomic ecosystem provides a complex community ecology of interacting families of TEs.

But unlike most ecological communities, which are censused when a researcher surveys them, the genomic ecosystem carries a record of past transposition. We can investigate this past ‘fossil’ record using the age of individual TE copies. This allows a robust analysis of the features that define TE survival across time. The TEs we see today are a readout of the joint processes of new transposition — which may not be uniform through time — and removal through selection, deletion, and drift (Tenaillon *et al*., 2010). Survival of a TE can be measured by its age or time since insertion, as our observation of a TE is conditioned on the fact it has not been removed by either neutral processes or selection. Our random forest model predicting age of TEs thus relates the action of transposition to the processes that occur afterwards on an evolutionary time scale. The model shows that nucleotide variation, measured as the proportion of segregating sites in the HapMap3 panel of 1,218 maize lines, provides the best predictive power for age (Figure 6B), and that age generally increases with nucleotide variation (Figure 6C,D). This positive relationship is expected, as the chance of a nucleotide mutation, required for measuring age, increases with time. We predict future efforts to address whether these insertions are shared across maize individuals will see a higher proportion of segregating sites in shared TEs. We caution, however, that SNP calls within and adjacent to TEs are likely incorrect, as nonhomologous TE insertions could be mapped to this locus in B73 and coverage cutoffs likely limit SNP calls in high copy number families.

Other predictors of age are expected. For example, we expect a new insertion to be younger if we show that the TE disrupts another TE. And we expect a higher proportion of segregating sites in the region flanking a TE insertion will be positively associated with TE age, as it reflects a count of the mutations that have accumulated on the haplotype carrying the TE. We note that imprecise repair of a double stranded break after excision of a TIR element (Wicker *et al*., 2016) could obscure this signal to some extent, increasing the number of flanking SNPs while decreasing the average frequency of the TE. Consistent with this mechanism, the superfamily DTT, which excises precisely without introducing nucleotide mutations (Gilbert *et al*., 2015) shows fewer flanking segregating sites per base pair (0.0421) than TIR elements from other superfamilies (0.0424).

Another TE feature with high predictive power for age and survival in the genome is the length of the TE. In other taxa, selection is stronger on long TEs, mediated by a higher potential for nonhomologous (ectopic) recombination (Petrov *et al*., 2003; Shen *et al*., 2013; Hollister and Gaut, 2007). We find that the relationship between TE length and age in maize is more nuanced, with some long TEs surviving over millions of years.Perhaps a genome as repetitive and TE rich as maize could not have evolved without mechanisms to prevent improper pairing of nonhomologous sequences with high nucleotide similarity.

Similarly, CHH methylation of TEs has been shown to be associated with recently activated TEs in *Arabidopsis thaliana* (Cavrak *et al*., 2014) and TEs near genes in maize (Gent *et al*., 2013; Li *et al*., 2015). Consistent with repressive activity of actively transposing TEs, we find complicated, nonlinear relationships of CHH methylation with age (Figure 6E,F). This likely reflects a natural senescence of a TE copy, where young copies are not yet silenced by the genome and lack CHH methylation, intermediate age copies effectively silenced with higher CHH methylation levels, and the oldest TEs with low CHH methylation as defunct copies incapabale of transposition that are no longer silenced.

Another explanatory variable with predictive power is TE expression in pollen, as expected since transposition in germline tissue is necessary for observing new TE copies. We observe an on average negative relationship between age and TE expression in pollen, although for some families there is a positive relationship. Since we use RNA-seq data from pooled mature pollen, we are unable to resolve expression in the vegetative and sperm nucleus. Some maize TE families show enriched expression in sperm cells (Vicient, 2010), while others are expressed in the vegetative nucleus, presumably as a mechanism to reinforce silencing in the sperm cell (Slotkin *et al*., 2009). This difference may explain why some families show opposite relationships with age, since sperm-expressed families are more likely to generate new insertions transmitted to the next generation than vegetative nucleus-expressed families for which transposition is silenced in the sperm cell.

In spite of previous predictions, distance to a gene and recombination are not found in the top 30 explanatory variables of age. Old TEs are underrepresented near genes in humans and *Arabidopsis thaliana* (Medstrand *et al*., 2002; Hollister and Gaut, 2009), consistent with selection against such insertions of long TEs. And recombination has been implicated in both the removal of TEs (Hill and Robertson, 1966) and modifying their impact on fitness via ectopic recombination (Charlesworth and Langley, 1989). Due to the colinearity of the genome and these features, distance to gene and recombination rate may be better represented by a combination of other features. For example, regions with high recombination rate generally show low CG methylation in maize (Rodgers-Melnick *et al*., 2015), but a subset of genes in such regions show CG methylation across the gene body. Since CG methylation plays a role in TE survival (Figure 6B), inclusion of this feature in our models will thus reduce the importance of recombination rate. And while CHH methylation is most prominent in regions of the genome close to genes (Gent *et al*., 2013), RNA-directed DNA methylation reinforces the boundary between heterochromatin and euchromatin, which is often over the TE closest to the gene (Li *et al*., 2015). Hence, the distance of a TE to the closest gene may provide redundant information about the TE, beyond what is provided by measurements of CHH methylation.

Finally, in spite of the fact our model includes more than 400 features of the genomic environment, TE taxonomy contributes substantially to prediction of TE age (Figure 6A). We have seen that relevance and direction of effect of individual features can differ among families (Figure 6C), essentially generating niche in the genomic ecosystem that a family survives within. The importance of taxonomy in our model may also suggest that there are still unmeasured latent variables that are best captured with superfamily and family labels. In fact, there is no genomic feature we measure which shows even the same direction of correlation across all families. This further emphasizes that it is essential to interpret individual families of TEs in the maize genome, such that their unique nature is not obscured by averages. The level of analysis of a TE in maize should be that of its family, as each family is surviving in a slightly different way, exploiting a unique genomic niche.

## Conclusion

Genes in the maize genome are ‘buried in non-genic DNA,’ consisting predominately of TEs (Bennetzen, 2009) and the interaction between TEs and the genes of the host genome can structure and inform genome function. The diversity of TEs in an elaborate genome like maize generates a complex ecosystem, with many interdependencies and nuances, limiting the ability to predict the functional consequences of a particular TE based only on superfamily or order. Instead, TE families represent a biologically relevant level on which to understand TE evolution, and the features most important for determining survival of individual copies represent dimensions of the ecological niche they inhabit. These observations suggest that the co-evolution between TE and host is ongoing, and inference of the impacts of transposons requires a multifaceted approach. The nuanced understanding generated from exhaustive analysis of genomic features and survival of individual families of TEs serves as a starting point to begin to understand not only TE evolution, but also the evolution of the host genomes they have coevolved with.

## ACKNOWLEDGEMENTS

M.C.S. and J.R.-I. are supported by the National Science Foundation Plant Genome award 1238014. M.C.S. acknowledges support from the National Science Foundation Graduate Research Fellowship under Grant No. 1650042; J.R.-I. acknowledges support from the USDA Hatch project CA-D-PLS-2066-H. S.N.A. and N.M.S. are supported by a grant from USDA-NIFA (2016-67013-24747).

