## Supplemental Figures and Tables for "The Genomic Ecosystem of Transposable Elements in Maize"

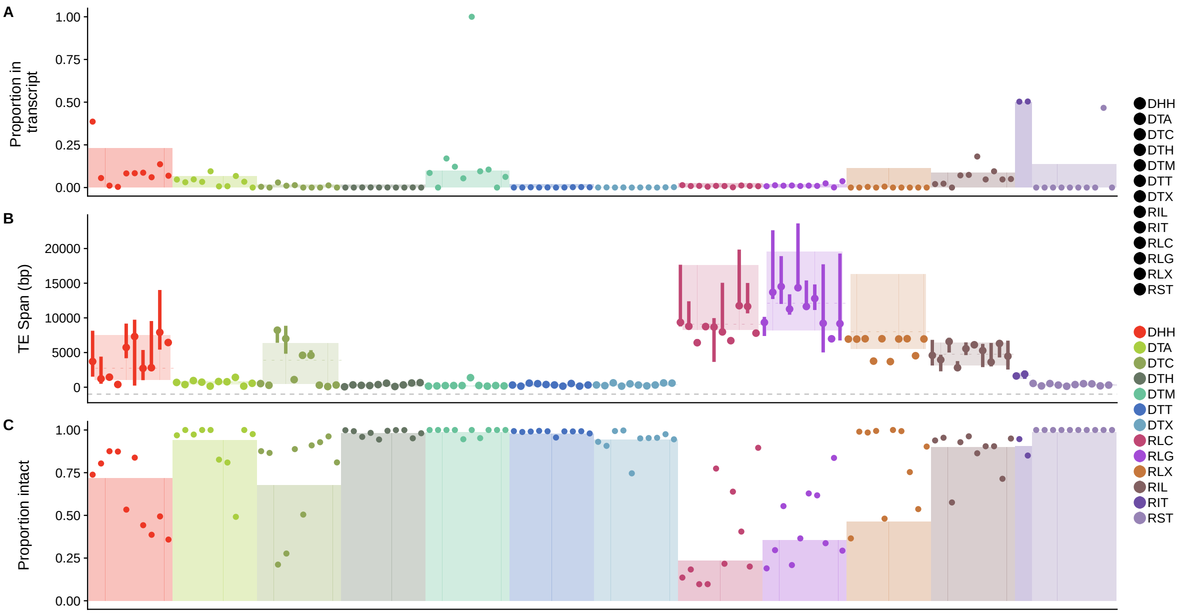


**Fig. S1. Family characteristics of each of the largest 10 families of each superfamily with at least 10 copies.** (A) Proportion of TEs within the transcript of a gene, including introns and UTRs. (B) TE span along the genome, summing both the base pairs of the TE and the base pairs of the TEs nested within it. (C) Proportion of TEs that are intact, that is, uninterrupted by the insertion of another TE. In (A and C), families are shown as points and superfamily proportions as a barplot, and in (B) families are shown with medians as points and lines representing ranges of upper to lower quartiles, with superfamilies shown as colored rectangles.

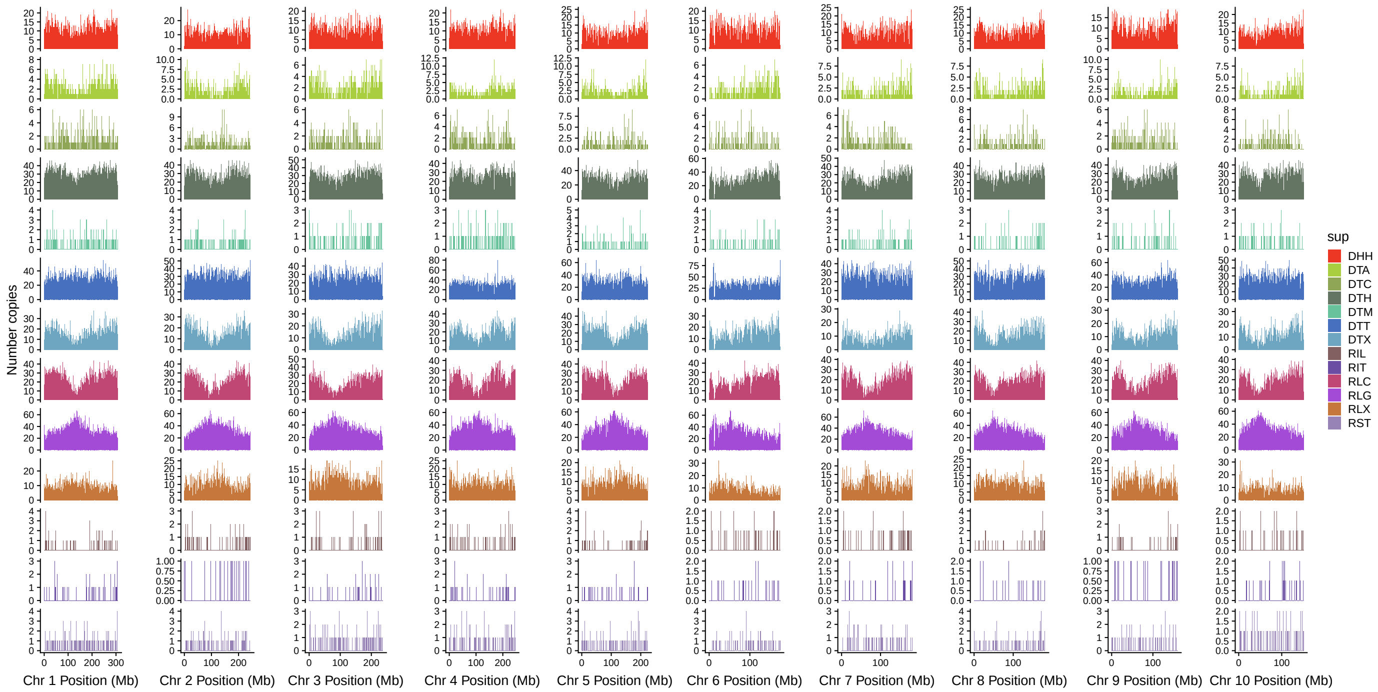


**Fig. S2. Chromosomal distribution of superfamilies across all 10 maize chromosomes** Count of TE copies of each superfamily in 1 megabase bins across each chromosome.

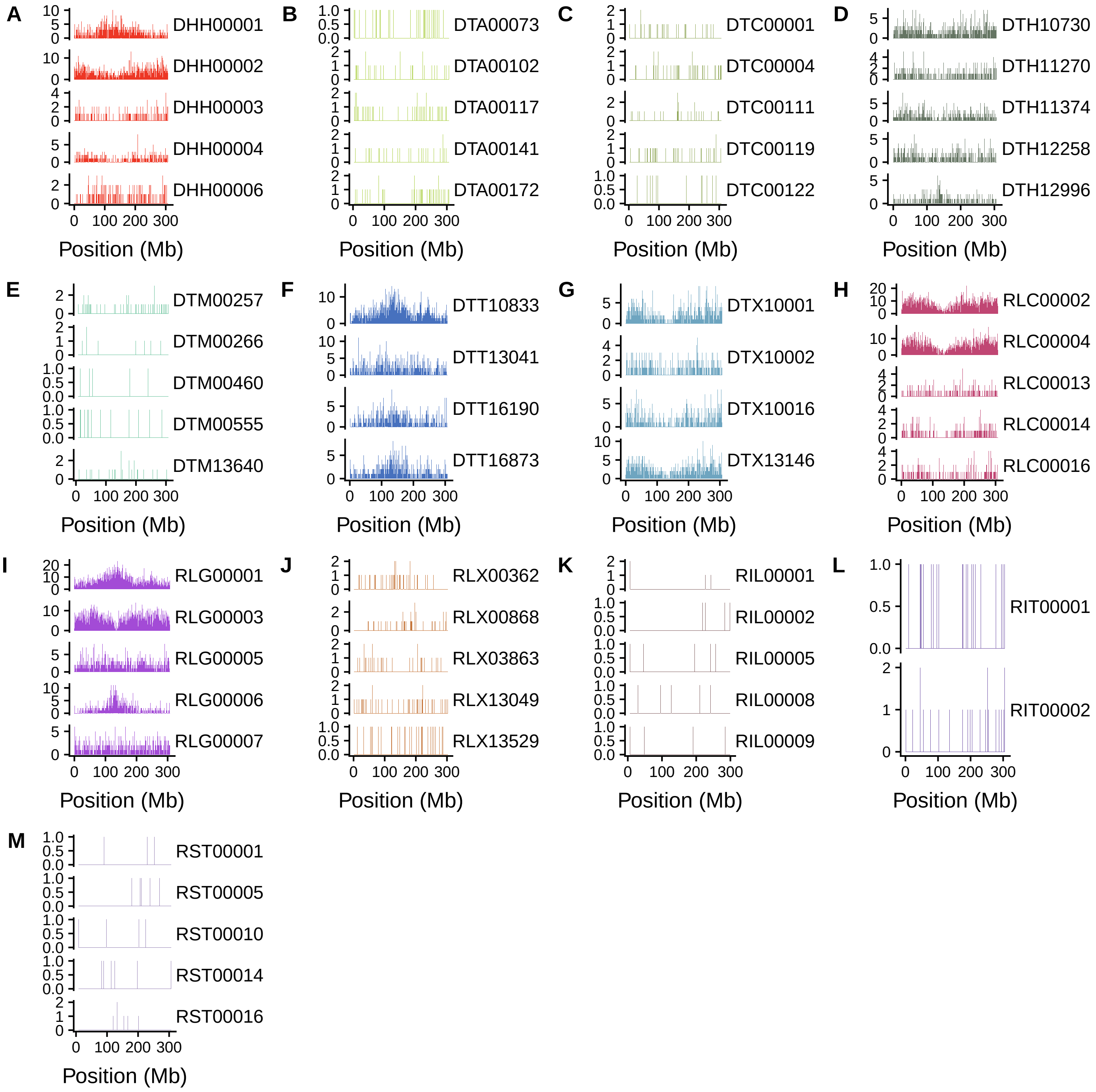


**Fig. S3. Distribution on chromosome 1 of five largest families with at least ten copies in each superfamily** Count of TE copies in 1 megabase bins along chromosome 1. (A) DHH, (B) DTA, (C) DTC, (D) DTH, (E) DTM, (F) DTT, (G) DTX, (H) RLC, (I) RLG, (J) RLX, (K) RIL, (L) RIT, (M) RST. Note that some families have no copies on chromosome 1, including DTT10880 and DTX10177. Additionally, the RIT superfamily only has two families.

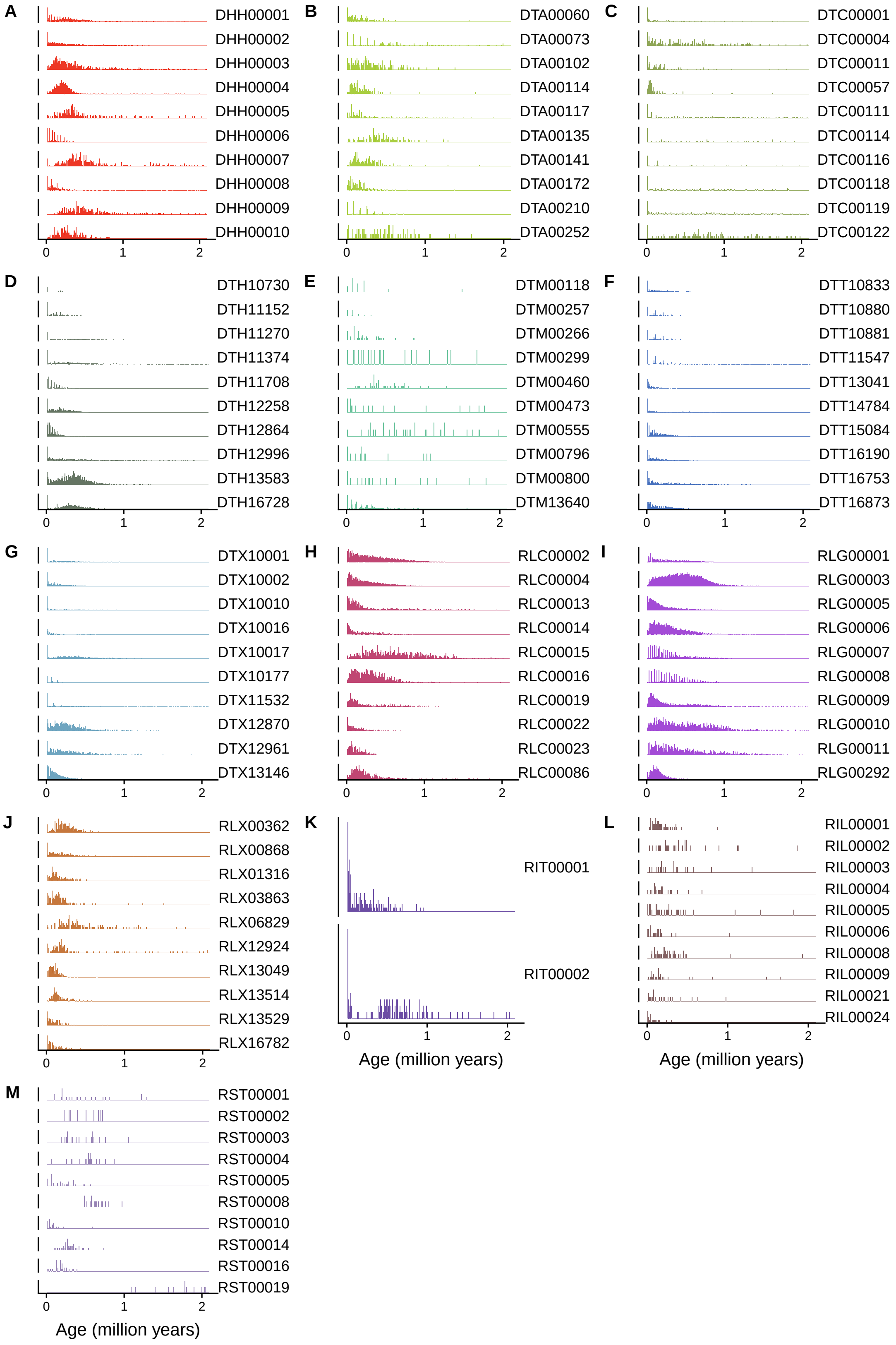


**Fig. S4. Ages in 10,000 year bins across each of the largest 10 families of each superfamily with at least 10 copies.** (A) DHH, (B) DTA, (C) DTC, (D) DTH, (E) DTM, (F) DTT, (G) DTX, (H) RLC, (I) RLG, (J) RLX, (K) RIL, (L) RIT, (M) RST. The RIT superfamily only contains two families.

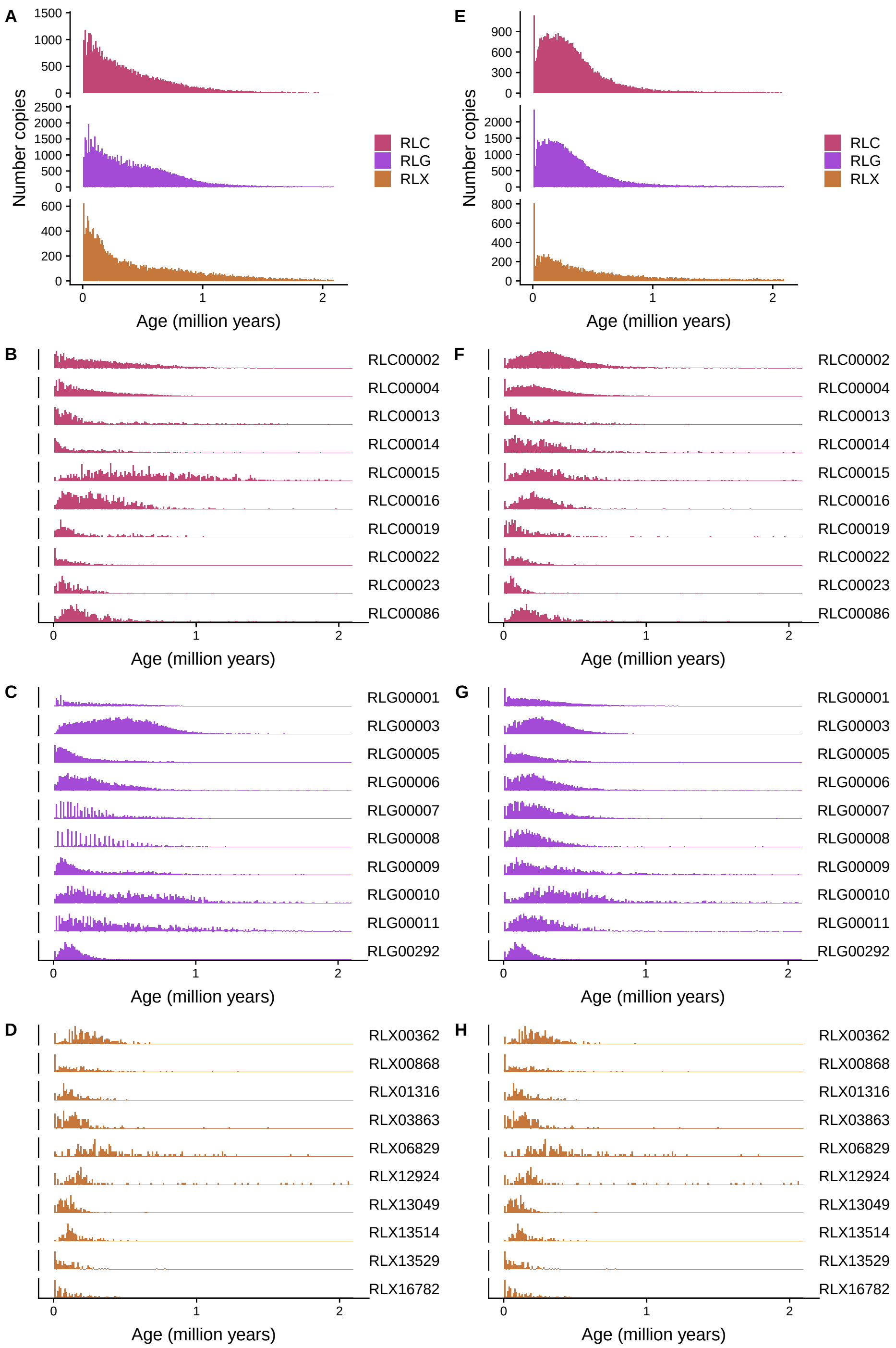


**Fig. S5. LTR-LTR ages and terminal branch length ages for LTR retrotransposons.** Ages in 10,000 year bins across each of the largest 10 families of each superfamily with at least 10 copies. Left plots (A-D) show LTR-LTR ages, right plots (E-H) show terminal branch length (TBL) ages. (A) all copies, LTR-LTR, (B) all copies, TBL, (C) RLC families, LTR-LTR, (D) RLC families, TBL, (E) RLG families, LTR-LTR, (F) RLG families, TBL, (G) RLX families, LTR-LTR, (H) RLX families, TBL.

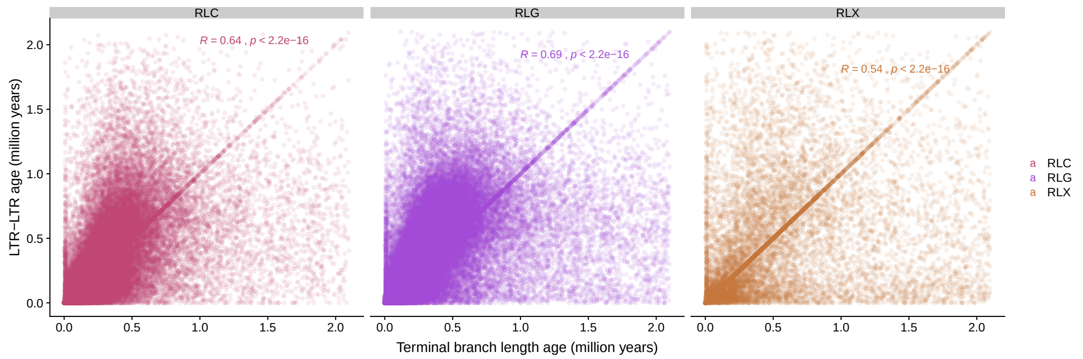


**Fig. S6. LTR-LTR ages vs. terminal branch length ages for LTR retrotransposon superfamilies.** Spearman's correlation coefficient shown on plot for each superfamily.

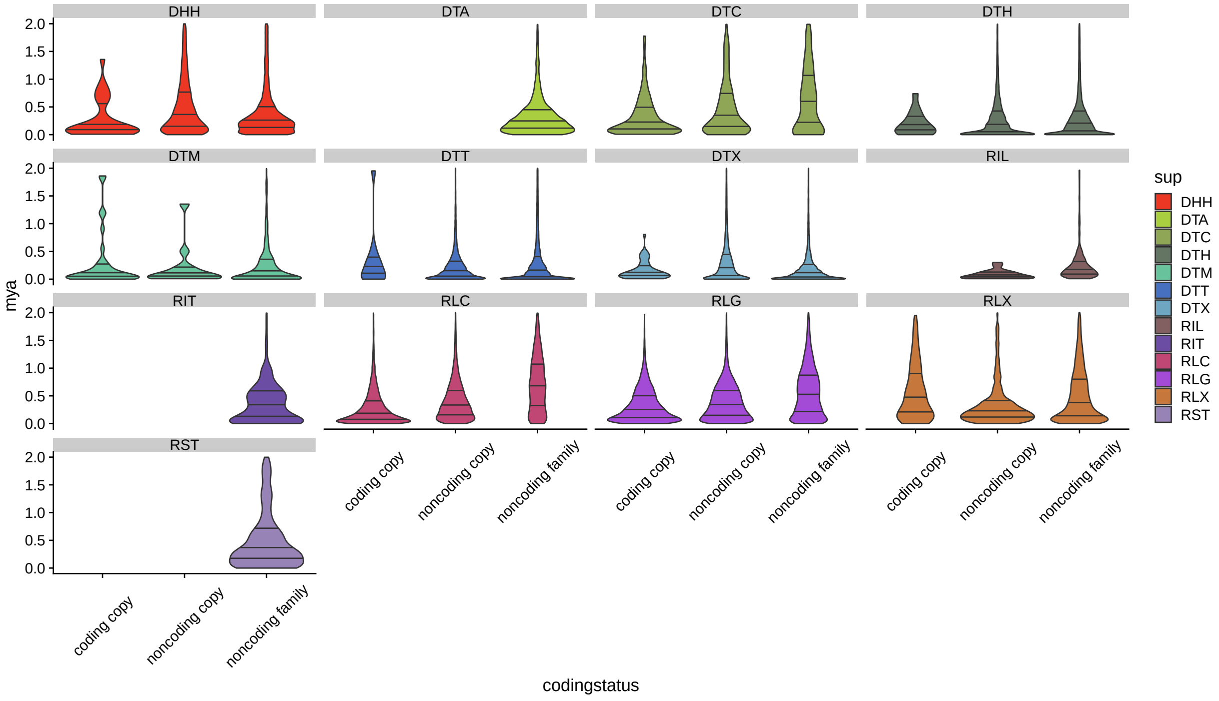


**Fig. S7. Age of TE copies split by coding potential of self and family** Violin plots with three lines, at median and 25th and 75th percentile. Only ages younger than 2 million years are shown. "Coding copy" refers to those copies that code for protein, "noncoding copy" refers to those copies that don't code for protein, but a family member does, and "noncoding family" refers to copies from families without a coding member in B73.

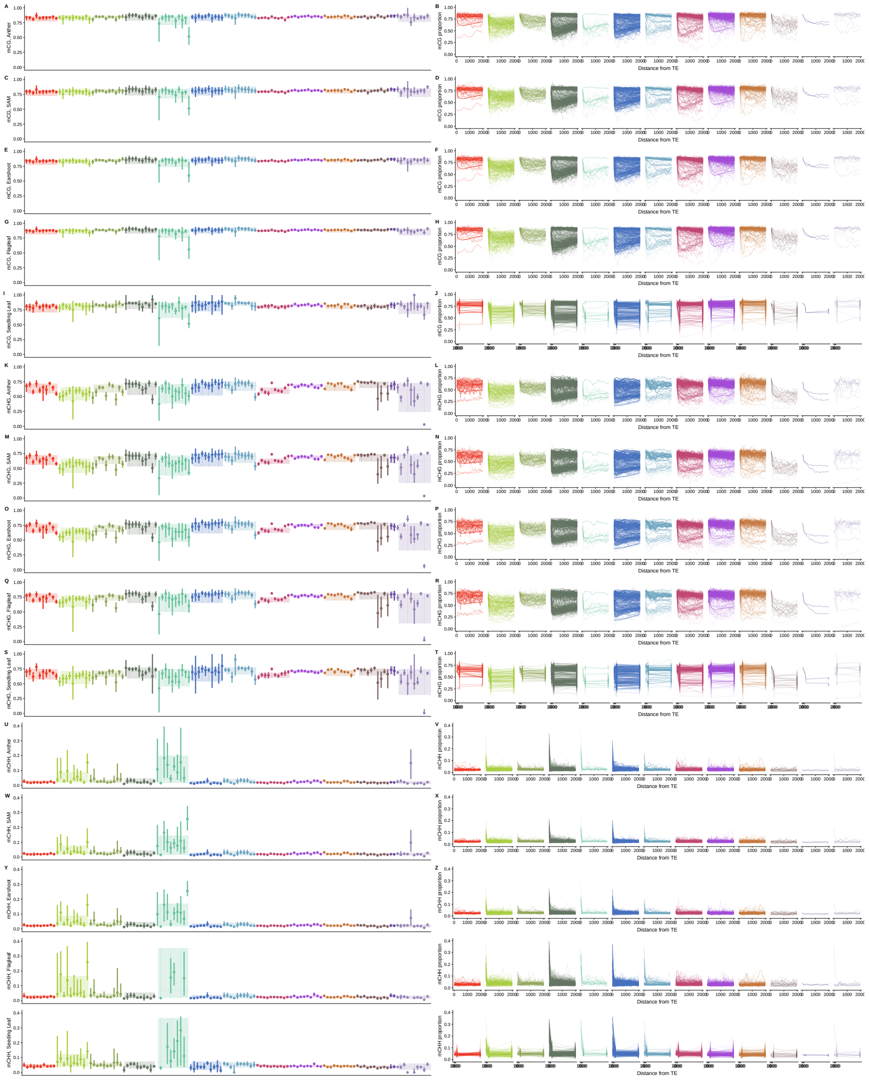

**Fig. S8. Methylation in TE and flanking sequence, across tissues**. A-J: mCG; K-T: mCHG; U-end mCHH. Tissues on y-axis, from top to bottom: Anther, SAM, Earshoot, Flagleaf, Seedling leaf.

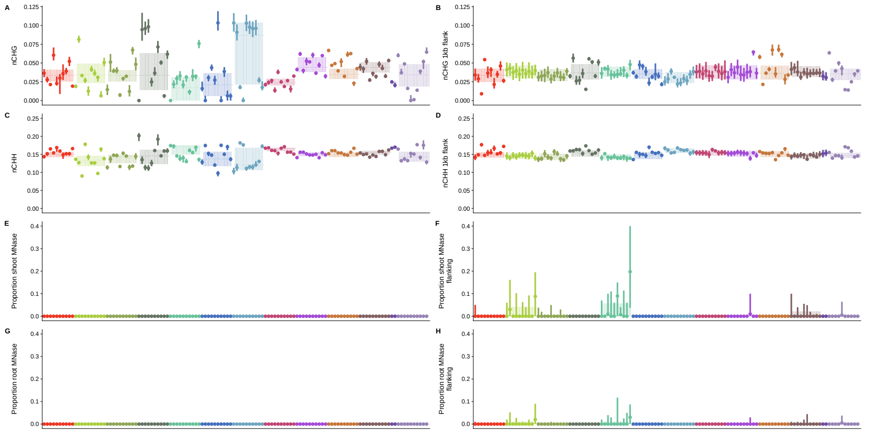

**Fig. S9. Features of the TE and flanking sequences**. Proportion of sites methylatable in CHG context in the TE (A) and 1kb flanking sequence (B), methylatable in the CHH context in the TE (C) and 1kb flanking sequence (D). Proportion of sites in MNase hypersensitive regions in shoot in TE (E) and 1kb flank (F), and root in TE (G) and 1kb flank (H).

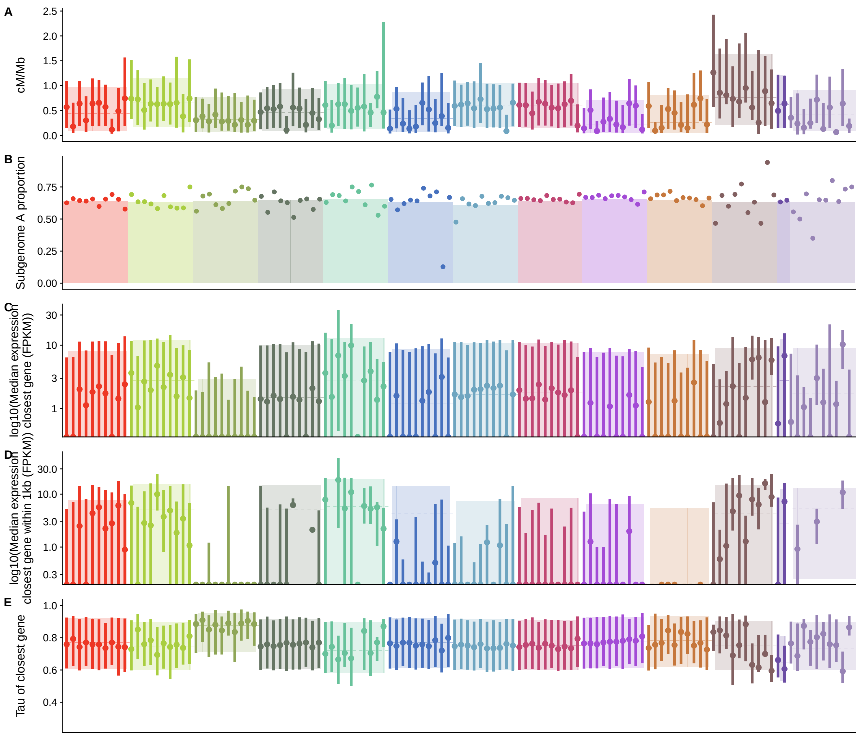


**Fig. S10. Recombination, subgenome, and expression of closest gene**. (A) Recombination rate across the TE, (B) proportion of TEs in subgenome A, (C) median expression of the closest gene to each TE, (D) median expression of genes within 1kb of the TE (E) Tau of closest gene to each TE.

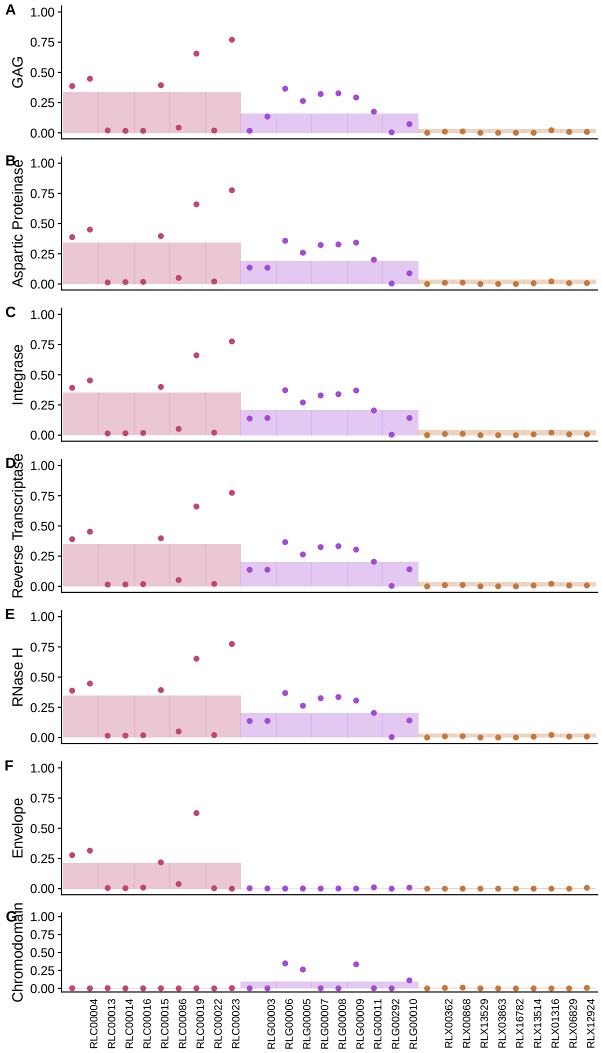


**Fig. S11. Protein coding gene presence of individual LTR GAG and POL domains** Shown are (A) the propotion of TEs with evidence of agglutination factor (GAG) domain present, (B) aspartic proteinase (AP) domains present, (C) Integrase (INT) domains present, (D) reverse transcriptase (RT) domains present, (E) RNaseH domains present, (F) envelope (ENV) domains present, and (G) chromodomain (CHR) domains present. Families are shown as points and superfamily proportions as barplot.

**Table S1.** Categories that each feature is classified into.

| feature | category |
|---|---|
| fam | TE_taxonomy |
| sup | TE_taxonomy |
| closest | flank_selection |
| tebp | TE_features |
| tespan | TE_features |
| pieces | TE_features |
| disruptor | TE_features |
| ingene | flank_selection |
| helprot | TE_encoded_proteins |
| rveprot | TE_encoded_proteins |
| tpaseprot | TE_encoded_proteins |
| GAG | TE_encoded_proteins |
| AP | TE_encoded_proteins |
| INT | TE_encoded_proteins |
| RT | TE_encoded_proteins |
| RNaseH | TE_encoded_proteins |
| ENV | TE_encoded_proteins |
| CHR | TE_encoded_proteins |
| pol | TE_encoded_proteins |
| auton | TE_encoded_proteins |
| GAGfam | TE_encoded_proteins |
| APfam | TE_encoded_proteins |
| INTfam | TE_encoded_proteins |
| RTfam | TE_encoded_proteins |
| RNaseHfam | TE_encoded_proteins |
| ENVfam | TE_encoded_proteins |
| CHRfam | TE_encoded_proteins |
| polfam | TE_encoded_proteins |
| autonfam | TE_encoded_proteins |
| helprotfam | TE_encoded_proteins |
| rveprotfam | TE_encoded_proteins |
| tpaseprotfam | TE_encoded_proteins |
| orfAA | TE_encoded_proteins |
| percGC | TE_base_composition |
| nCG | TE_base_composition |
| nCHG | TE_base_composition |
| nCHH | TE_base_composition |
| percGC_1kbflank | flank_base_composition |
| nCG_1kbflank | flank_base_composition |
| nCHG_1kbflank | flank_base_composition |
| nCHH_1kbflank | flank_base_composition |
| n_root_hs | TE_methylation_mnase |
| root_bp | TE_methylation_mnase |
| root_prop | TE_methylation_mnase |
| n_shoot_hs | TE_methylation_mnase |
| shoot_bp | TE_methylation_mnase |
| shoot_prop | TE_methylation_mnase |
| flank_n_root_hs | flank_methylation_mnase |
| flank_root_bp | flank_methylation_mnase |
| flank_root_prop | flank_methylation_mnase |
| flank_n_shoot_hs | flank_methylation_mnase |
| flank_shoot_bp | flank_methylation_mnase |
| flank_shoot_prop | flank_methylation_mnase |
| segsites.bp | TE_features |

| | |
|---|---|
| flank_segsites.bp | flank_selection |
| anther_avg_cg | TE_methylation_mnase |
| anther_avg_chg | TE_methylation_mnase |
| anther_avg_chh | TE_methylation_mnase |
| anther_flank_cg_100 | flank_methylation_mnase |
| anther_flank_cg_200 | flank_methylation_mnase |
| anther_flank_cg_300 | flank_methylation_mnase |
| anther_flank_cg_400 | flank_methylation_mnase |
| anther_flank_cg_500 | flank_methylation_mnase |
| anther_flank_cg_600 | flank_methylation_mnase |
| anther_flank_cg_700 | flank_methylation_mnase |
| anther_flank_cg_800 | flank_methylation_mnase |
| anther_flank_cg_900 | flank_methylation_mnase |
| anther_flank_cg_1000 | flank_methylation_mnase |
| anther_flank_cg_1100 | flank_methylation_mnase |
| anther_flank_cg_1200 | flank_methylation_mnase |
| anther_flank_cg_1300 | flank_methylation_mnase |
| anther_flank_cg_1400 | flank_methylation_mnase |
| anther_flank_cg_1500 | flank_methylation_mnase |
| anther_flank_cg_1600 | flank_methylation_mnase |
| anther_flank_cg_1700 | flank_methylation_mnase |
| anther_flank_cg_1800 | flank_methylation_mnase |
| anther_flank_cg_1900 | flank_methylation_mnase |
| anther_flank_cg_2000 | flank_methylation_mnase |
| anther_flank_chg_100 | flank_methylation_mnase |
| anther_flank_chg_200 | flank_methylation_mnase |
| anther_flank_chg_300 | flank_methylation_mnase |
| anther_flank_chg_400 | flank_methylation_mnase |
| anther_flank_chg_500 | flank_methylation_mnase |
| anther_flank_chg_600 | flank_methylation_mnase |
| anther_flank_chg_700 | flank_methylation_mnase |
| anther_flank_chg_800 | flank_methylation_mnase |
| anther_flank_chg_900 | flank_methylation_mnase |
| anther_flank_chg_1000 | flank_methylation_mnase |
| anther_flank_chg_1100 | flank_methylation_mnase |
| anther_flank_chg_1200 | flank_methylation_mnase |
| anther_flank_chg_1300 | flank_methylation_mnase |
| anther_flank_chg_1400 | flank_methylation_mnase |
| anther_flank_chg_1500 | flank_methylation_mnase |
| anther_flank_chg_1600 | flank_methylation_mnase |
| anther_flank_chg_1700 | flank_methylation_mnase |
| anther_flank_chg_1800 | flank_methylation_mnase |
| anther_flank_chg_1900 | flank_methylation_mnase |
| anther_flank_chg_2000 | flank_methylation_mnase |
| anther_flank_chh_100 | flank_methylation_mnase |
| anther_flank_chh_200 | flank_methylation_mnase |
| anther_flank_chh_300 | flank_methylation_mnase |
| anther_flank_chh_400 | flank_methylation_mnase |
| anther_flank_chh_500 | flank_methylation_mnase |
| anther_flank_chh_600 | flank_methylation_mnase |
| anther_flank_chh_700 | flank_methylation_mnase |
| anther_flank_chh_800 | flank_methylation_mnase |
| anther_flank_chh_900 | flank_methylation_mnase |
| anther_flank_chh_1000 | flank_methylation_mnase |
| anther_flank_chh_1100 | flank_methylation_mnase |
| anther_flank_chh_1200 | flank_methylation_mnase |
| anther_flank_chh_1300 | flank_methylation_mnase |

| | |
|---|---|
| anther_flank_chh_1400 | flank_methylation_mnase |
| anther_flank_chh_1500 | flank_methylation_mnase |
| anther_flank_chh_1600 | flank_methylation_mnase |
| anther_flank_chh_1700 | flank_methylation_mnase |
| anther_flank_chh_1800 | flank_methylation_mnase |
| anther_flank_chh_1900 | flank_methylation_mnase |
| anther_flank_chh_2000 | flank_methylation_mnase |
| earshoot_avg_cg | TE_methylation_mnase |
| earshoot_avg_chg | TE_methylation_mnase |
| earshoot_avg_chh | TE_methylation_mnase |
| earshoot_flank_cg_100 | flank_methylation_mnase |
| earshoot_flank_cg_200 | flank_methylation_mnase |
| earshoot_flank_cg_300 | flank_methylation_mnase |
| earshoot_flank_cg_400 | flank_methylation_mnase |
| earshoot_flank_cg_500 | flank_methylation_mnase |
| earshoot_flank_cg_600 | flank_methylation_mnase |
| earshoot_flank_cg_700 | flank_methylation_mnase |
| earshoot_flank_cg_800 | flank_methylation_mnase |
| earshoot_flank_cg_900 | flank_methylation_mnase |
| earshoot_flank_cg_1000 | flank_methylation_mnase |
| earshoot_flank_cg_1100 | flank_methylation_mnase |
| earshoot_flank_cg_1200 | flank_methylation_mnase |
| earshoot_flank_cg_1300 | flank_methylation_mnase |
| earshoot_flank_cg_1400 | flank_methylation_mnase |
| earshoot_flank_cg_1500 | flank_methylation_mnase |
| earshoot_flank_cg_1600 | flank_methylation_mnase |
| earshoot_flank_cg_1700 | flank_methylation_mnase |
| earshoot_flank_cg_1800 | flank_methylation_mnase |
| earshoot_flank_cg_1900 | flank_methylation_mnase |
| earshoot_flank_cg_2000 | flank_methylation_mnase |
| earshoot_flank_chg_100 | flank_methylation_mnase |
| earshoot_flank_chg_200 | flank_methylation_mnase |
| earshoot_flank_chg_300 | flank_methylation_mnase |
| earshoot_flank_chg_400 | flank_methylation_mnase |
| earshoot_flank_chg_500 | flank_methylation_mnase |
| earshoot_flank_chg_600 | flank_methylation_mnase |
| earshoot_flank_chg_700 | flank_methylation_mnase |
| earshoot_flank_chg_800 | flank_methylation_mnase |
| earshoot_flank_chg_900 | flank_methylation_mnase |
| earshoot_flank_chg_1000 | flank_methylation_mnase |
| earshoot_flank_chg_1100 | flank_methylation_mnase |
| earshoot_flank_chg_1200 | flank_methylation_mnase |
| earshoot_flank_chg_1300 | flank_methylation_mnase |
| earshoot_flank_chg_1400 | flank_methylation_mnase |
| earshoot_flank_chg_1500 | flank_methylation_mnase |
| earshoot_flank_chg_1600 | flank_methylation_mnase |
| earshoot_flank_chg_1700 | flank_methylation_mnase |
| earshoot_flank_chg_1800 | flank_methylation_mnase |
| earshoot_flank_chg_1900 | flank_methylation_mnase |
| earshoot_flank_chg_2000 | flank_methylation_mnase |
| earshoot_flank_chh_100 | flank_methylation_mnase |
| earshoot_flank_chh_200 | flank_methylation_mnase |
| earshoot_flank_chh_300 | flank_methylation_mnase |
| earshoot_flank_chh_400 | flank_methylation_mnase |
| earshoot_flank_chh_500 | flank_methylation_mnase |
| earshoot_flank_chh_600 | flank_methylation_mnase |
| earshoot_flank_chh_700 | flank_methylation_mnase |

| | |
|---|---|
| earshoot_flank_chh_800 | flank_methylation_mnase |
| earshoot_flank_chh_900 | flank_methylation_mnase |
| earshoot_flank_chh_1000 | flank_methylation_mnase |
| earshoot_flank_chh_1100 | flank_methylation_mnase |
| earshoot_flank_chh_1200 | flank_methylation_mnase |
| earshoot_flank_chh_1300 | flank_methylation_mnase |
| earshoot_flank_chh_1400 | flank_methylation_mnase |
| earshoot_flank_chh_1500 | flank_methylation_mnase |
| earshoot_flank_chh_1600 | flank_methylation_mnase |
| earshoot_flank_chh_1700 | flank_methylation_mnase |
| earshoot_flank_chh_1800 | flank_methylation_mnase |
| earshoot_flank_chh_1900 | flank_methylation_mnase |
| earshoot_flank_chh_2000 | flank_methylation_mnase |
| flagleaf_avg_cg | TE_methylation_mnase |
| flagleaf_avg_chg | TE_methylation_mnase |
| flagleaf_avg_chh | TE_methylation_mnase |
| flagleaf_flank_cg_100 | flank_methylation_mnase |
| flagleaf_flank_cg_200 | flank_methylation_mnase |
| flagleaf_flank_cg_300 | flank_methylation_mnase |
| flagleaf_flank_cg_400 | flank_methylation_mnase |
| flagleaf_flank_cg_500 | flank_methylation_mnase |
| flagleaf_flank_cg_600 | flank_methylation_mnase |
| flagleaf_flank_cg_700 | flank_methylation_mnase |
| flagleaf_flank_cg_800 | flank_methylation_mnase |
| flagleaf_flank_cg_900 | flank_methylation_mnase |
| flagleaf_flank_cg_1000 | flank_methylation_mnase |
| flagleaf_flank_cg_1100 | flank_methylation_mnase |
| flagleaf_flank_cg_1200 | flank_methylation_mnase |
| flagleaf_flank_cg_1300 | flank_methylation_mnase |
| flagleaf_flank_cg_1400 | flank_methylation_mnase |
| flagleaf_flank_cg_1500 | flank_methylation_mnase |
| flagleaf_flank_cg_1600 | flank_methylation_mnase |
| flagleaf_flank_cg_1700 | flank_methylation_mnase |
| flagleaf_flank_cg_1800 | flank_methylation_mnase |
| flagleaf_flank_cg_1900 | flank_methylation_mnase |
| flagleaf_flank_cg_2000 | flank_methylation_mnase |
| flagleaf_flank_chg_100 | flank_methylation_mnase |
| flagleaf_flank_chg_200 | flank_methylation_mnase |
| flagleaf_flank_chg_300 | flank_methylation_mnase |
| flagleaf_flank_chg_400 | flank_methylation_mnase |
| flagleaf_flank_chg_500 | flank_methylation_mnase |
| flagleaf_flank_chg_600 | flank_methylation_mnase |
| flagleaf_flank_chg_700 | flank_methylation_mnase |
| flagleaf_flank_chg_800 | flank_methylation_mnase |
| flagleaf_flank_chg_900 | flank_methylation_mnase |
| flagleaf_flank_chg_1000 | flank_methylation_mnase |
| flagleaf_flank_chg_1100 | flank_methylation_mnase |
| flagleaf_flank_chg_1200 | flank_methylation_mnase |
| flagleaf_flank_chg_1300 | flank_methylation_mnase |
| flagleaf_flank_chg_1400 | flank_methylation_mnase |
| flagleaf_flank_chg_1500 | flank_methylation_mnase |
| flagleaf_flank_chg_1600 | flank_methylation_mnase |
| flagleaf_flank_chg_1700 | flank_methylation_mnase |
| flagleaf_flank_chg_1800 | flank_methylation_mnase |
| flagleaf_flank_chg_1900 | flank_methylation_mnase |
| flagleaf_flank_chg_2000 | flank_methylation_mnase |
| flagleaf_flank_chh_100 | flank_methylation_mnase |

| | |
|---|---|
| flagleaf_flank_chh_200 | flank_methylation_mnase |
| flagleaf_flank_chh_300 | flank_methylation_mnase |
| flagleaf_flank_chh_400 | flank_methylation_mnase |
| flagleaf_flank_chh_500 | flank_methylation_mnase |
| flagleaf_flank_chh_600 | flank_methylation_mnase |
| flagleaf_flank_chh_700 | flank_methylation_mnase |
| flagleaf_flank_chh_800 | flank_methylation_mnase |
| flagleaf_flank_chh_900 | flank_methylation_mnase |
| flagleaf_flank_chh_1000 | flank_methylation_mnase |
| flagleaf_flank_chh_1100 | flank_methylation_mnase |
| flagleaf_flank_chh_1200 | flank_methylation_mnase |
| flagleaf_flank_chh_1300 | flank_methylation_mnase |
| flagleaf_flank_chh_1400 | flank_methylation_mnase |
| flagleaf_flank_chh_1500 | flank_methylation_mnase |
| flagleaf_flank_chh_1600 | flank_methylation_mnase |
| flagleaf_flank_chh_1700 | flank_methylation_mnase |
| flagleaf_flank_chh_1800 | flank_methylation_mnase |
| flagleaf_flank_chh_1900 | flank_methylation_mnase |
| flagleaf_flank_chh_2000 | flank_methylation_mnase |
| all3_avg_cg | TE_methylation_mnase |
| all3_avg_chg | TE_methylation_mnase |
| all3_avg_chh | TE_methylation_mnase |
| all3_flank_cg_100 | flank_methylation_mnase |
| all3_flank_cg_200 | flank_methylation_mnase |
| all3_flank_cg_300 | flank_methylation_mnase |
| all3_flank_cg_400 | flank_methylation_mnase |
| all3_flank_cg_500 | flank_methylation_mnase |
| all3_flank_cg_600 | flank_methylation_mnase |
| all3_flank_cg_700 | flank_methylation_mnase |
| all3_flank_cg_800 | flank_methylation_mnase |
| all3_flank_cg_900 | flank_methylation_mnase |
| all3_flank_cg_1000 | flank_methylation_mnase |
| all3_flank_cg_1100 | flank_methylation_mnase |
| all3_flank_cg_1200 | flank_methylation_mnase |
| all3_flank_cg_1300 | flank_methylation_mnase |
| all3_flank_cg_1400 | flank_methylation_mnase |
| all3_flank_cg_1500 | flank_methylation_mnase |
| all3_flank_cg_1600 | flank_methylation_mnase |
| all3_flank_cg_1700 | flank_methylation_mnase |
| all3_flank_cg_1800 | flank_methylation_mnase |
| all3_flank_cg_1900 | flank_methylation_mnase |
| all3_flank_cg_2000 | flank_methylation_mnase |
| all3_flank_chg_100 | flank_methylation_mnase |
| all3_flank_chg_200 | flank_methylation_mnase |
| all3_flank_chg_300 | flank_methylation_mnase |
| all3_flank_chg_400 | flank_methylation_mnase |
| all3_flank_chg_500 | flank_methylation_mnase |
| all3_flank_chg_600 | flank_methylation_mnase |
| all3_flank_chg_700 | flank_methylation_mnase |
| all3_flank_chg_800 | flank_methylation_mnase |
| all3_flank_chg_900 | flank_methylation_mnase |
| all3_flank_chg_1000 | flank_methylation_mnase |
| all3_flank_chg_1100 | flank_methylation_mnase |
| all3_flank_chg_1200 | flank_methylation_mnase |
| all3_flank_chg_1300 | flank_methylation_mnase |
| all3_flank_chg_1400 | flank_methylation_mnase |
| all3_flank_chg_1500 | flank_methylation_mnase |

| | |
|---|---|
| all3_flank_chg_1600 | flank_methylation_mnase |
| all3_flank_chg_1700 | flank_methylation_mnase |
| all3_flank_chg_1800 | flank_methylation_mnase |
| all3_flank_chg_1900 | flank_methylation_mnase |
| all3_flank_chg_2000 | flank_methylation_mnase |
| all3_flank_chh_100 | flank_methylation_mnase |
| all3_flank_chh_200 | flank_methylation_mnase |
| all3_flank_chh_300 | flank_methylation_mnase |
| all3_flank_chh_400 | flank_methylation_mnase |
| all3_flank_chh_500 | flank_methylation_mnase |
| all3_flank_chh_600 | flank_methylation_mnase |
| all3_flank_chh_700 | flank_methylation_mnase |
| all3_flank_chh_800 | flank_methylation_mnase |
| all3_flank_chh_900 | flank_methylation_mnase |
| all3_flank_chh_1000 | flank_methylation_mnase |
| all3_flank_chh_1100 | flank_methylation_mnase |
| all3_flank_chh_1200 | flank_methylation_mnase |
| all3_flank_chh_1300 | flank_methylation_mnase |
| all3_flank_chh_1400 | flank_methylation_mnase |
| all3_flank_chh_1500 | flank_methylation_mnase |
| all3_flank_chh_1600 | flank_methylation_mnase |
| all3_flank_chh_1700 | flank_methylation_mnase |
| all3_flank_chh_1800 | flank_methylation_mnase |
| all3_flank_chh_1900 | flank_methylation_mnase |
| all3_flank_chh_2000 | flank_methylation_mnase |
| SAM_avg_cg | TE_methylation_mnase |
| SAM_avg_chg | TE_methylation_mnase |
| SAM_avg_chh | TE_methylation_mnase |
| SAM_flank_cg_100 | flank_methylation_mnase |
| SAM_flank_cg_200 | flank_methylation_mnase |
| SAM_flank_cg_300 | flank_methylation_mnase |
| SAM_flank_cg_400 | flank_methylation_mnase |
| SAM_flank_cg_500 | flank_methylation_mnase |
| SAM_flank_cg_600 | flank_methylation_mnase |
| SAM_flank_cg_700 | flank_methylation_mnase |
| SAM_flank_cg_800 | flank_methylation_mnase |
| SAM_flank_cg_900 | flank_methylation_mnase |
| SAM_flank_cg_1000 | flank_methylation_mnase |
| SAM_flank_cg_1100 | flank_methylation_mnase |
| SAM_flank_cg_1200 | flank_methylation_mnase |
| SAM_flank_cg_1300 | flank_methylation_mnase |
| SAM_flank_cg_1400 | flank_methylation_mnase |
| SAM_flank_cg_1500 | flank_methylation_mnase |
| SAM_flank_cg_1600 | flank_methylation_mnase |
| SAM_flank_cg_1700 | flank_methylation_mnase |
| SAM_flank_cg_1800 | flank_methylation_mnase |
| SAM_flank_cg_1900 | flank_methylation_mnase |
| SAM_flank_cg_2000 | flank_methylation_mnase |
| SAM_flank_chg_100 | flank_methylation_mnase |
| SAM_flank_chg_200 | flank_methylation_mnase |
| SAM_flank_chg_300 | flank_methylation_mnase |
| SAM_flank_chg_400 | flank_methylation_mnase |
| SAM_flank_chg_500 | flank_methylation_mnase |
| SAM_flank_chg_600 | flank_methylation_mnase |
| SAM_flank_chg_700 | flank_methylation_mnase |
| SAM_flank_chg_800 | flank_methylation_mnase |
| SAM_flank_chg_900 | flank_methylation_mnase |

| | |
|---|---|
| SAM_flank_chg_1000 | flank_methylation_mnase |
| SAM_flank_chg_1100 | flank_methylation_mnase |
| SAM_flank_chg_1200 | flank_methylation_mnase |
| SAM_flank_chg_1300 | flank_methylation_mnase |
| SAM_flank_chg_1400 | flank_methylation_mnase |
| SAM_flank_chg_1500 | flank_methylation_mnase |
| SAM_flank_chg_1600 | flank_methylation_mnase |
| SAM_flank_chg_1700 | flank_methylation_mnase |
| SAM_flank_chg_1800 | flank_methylation_mnase |
| SAM_flank_chg_1900 | flank_methylation_mnase |
| SAM_flank_chg_2000 | flank_methylation_mnase |
| SAM_flank_chh_100 | flank_methylation_mnase |
| SAM_flank_chh_200 | flank_methylation_mnase |
| SAM_flank_chh_300 | flank_methylation_mnase |
| SAM_flank_chh_400 | flank_methylation_mnase |
| SAM_flank_chh_500 | flank_methylation_mnase |
| SAM_flank_chh_600 | flank_methylation_mnase |
| SAM_flank_chh_700 | flank_methylation_mnase |
| SAM_flank_chh_800 | flank_methylation_mnase |
| SAM_flank_chh_900 | flank_methylation_mnase |
| SAM_flank_chh_1000 | flank_methylation_mnase |
| SAM_flank_chh_1100 | flank_methylation_mnase |
| SAM_flank_chh_1200 | flank_methylation_mnase |
| SAM_flank_chh_1300 | flank_methylation_mnase |
| SAM_flank_chh_1400 | flank_methylation_mnase |
| SAM_flank_chh_1500 | flank_methylation_mnase |
| SAM_flank_chh_1600 | flank_methylation_mnase |
| SAM_flank_chh_1700 | flank_methylation_mnase |
| SAM_flank_chh_1800 | flank_methylation_mnase |
| SAM_flank_chh_1900 | flank_methylation_mnase |
| SAM_flank_chh_2000 | flank_methylation_mnase |
| gene_6_7_internode | flank_closest_gene_expression |
| gene_7_8_internode | flank_closest_gene_expression |
| gene_Vegetative_Meristem_16_19_Day | flank_closest_gene_expression |
| gene_Ear_Primordium_2_4_mm | flank_closest_gene_expression |
| gene_Ear_Primordium_6_8_mm | flank_closest_gene_expression |
| gene_Embryo_20_DAP | flank_closest_gene_expression |
| gene_Embryo_38_DAP | flank_closest_gene_expression |
| gene_Endosperm_12_DAP | flank_closest_gene_expression |
| gene_Endosperm_Crown_27_DAP | flank_closest_gene_expression |
| gene_Germinatin_Kernels_2_DAI | flank_closest_gene_expression |
| gene_Pericarp_Aleurone_27_DAP | flank_closest_gene_expression |
| gene_Leaf_Zone_1__Symmetrical_ | flank_closest_gene_expression |
| gene_Leaf_Zone_2__Stomatal_ | flank_closest_gene_expression |
| gene_Leaf_Zone_3__Growth_ | flank_closest_gene_expression |
| gene_Mature_Leaf_8 | flank_closest_gene_expression |
| gene_Primary_Root_5_Days | flank_closest_gene_expression |
| gene_Root___Cortex_5_Days | flank_closest_gene_expression |
| gene_Root___Elongation_Zone_5_Days | flank_closest_gene_expression |
| gene_Root___Meristem_Zone_5_Days | flank_closest_gene_expression |
| gene_Secondary_Root_7_8_Days | flank_closest_gene_expression |
| gene_B73_Mature_Pollen | flank_closest_gene_expression |
| gene_Female_Spikelet_Collected_on_day_as_silk | flank_closest_gene_expression |
| gene_Silk | flank_closest_gene_expression |
| famsize | TE_features |
| gene_coefvar | flank_closest_gene_expression |
| gene_median | flank_closest_gene_expression |

| | |
|---|---|
| TEfam_ear_primordium.2mm | TE_expression |
| TEfam_ear_primordium.6mm | TE_expression |
| TEfam_embryo_20d | TE_expression |
| TEfam_embryo_38d | TE_expression |
| TEfam_endosperm_12d | TE_expression |
| TEfam_endosperm_crown | TE_expression |
| TEfam_germinating.kernels_2d | TE_expression |
| TEfam_internode_6to7 | TE_expression |
| TEfam_internode_7to8 | TE_expression |
| TEfam_leaf_8 | TE_expression |
| TEfam_leaf_growth.zone | TE_expression |
| TEfam_leaf_stomatal.zone | TE_expression |
| TEfam_leaf_symmetrical.zone | TE_expression |
| TEfam_meristem_vegetative | TE_expression |
| TEfam_pollen_mature | TE_expression |
| TEfam_root_cortex | TE_expression |
| TEfam_root_elongation.zone | TE_expression |
| TEfam_root_meristem.zone | TE_expression |
| TEfam_root_primary | TE_expression |
| TEfam_root_secondary | TE_expression |
| TEfam_seed_pericarp.aleurone | TE_expression |
| TEfam_silk_mature | TE_expression |
| TEfam_spikelet_female | TE_expression |
| TEfamMedian | TE_expression |
| TEfamMedianPerCopy | TE_expression |
| TEfamMedianPerBp | TE_expression |
| TEfam_tau | TE_expression |
| cmmb | flank_selection |
| subgenome | flank_selection |

**Table S2.** 14 families with at least 10 copies in the B73 genome, with at least 75% of copies coding for transposition related proteins.

| sup | fam | propAuton | famsize | medianORF | meanORF |
|---|---|---|---|---|---|
| DTC | DTC00001 | 0.773399014778325 | 203 | 614 | 591.751269035533 |
| DTC | DTC00014 | 0.833333333333333 | 12 | 678 | 635.75 |
| DTC | DTC00017 | 0.9 | 10 | 448 | 453.2 |
| DTC | DTC00027 | 0.9 | 20 | 571.5 | 580.1 |
| DTC | DTC00041 | 0.8 | 10 | 388 | 407.4 |
| DTC | DTC00053 | 0.75 | 28 | 745.5 | 724.821428571429 |
| DTC | DTC00063 | 0.75 | 12 | 495.5 | 480.916666666667 |
| RLC | RLC00023 | 0.76878612716763 | 519 | 992 | 941.737769080235 |
| RLC | RLC00074 | 0.765625 | 64 | 901.5 | 928.3125 |
| RLC | RLC00137 | 0.857142857142857 | 28 | 1287.5 | 1072.78571428571 |
| RLC | RLC00149 | 0.777777777777778 | 27 | 1253 | 1103.11111111111 |
| RLC | RLC00158 | 0.8 | 25 | 769 | 894.84 |
| RLC | RLC00232 | 0.882352941176471 | 17 | 1473 | 1147.17647058824 |
| RLC | RLC00252 | 0.8 | 15 | 1112 | 1163.26666666667 |
| RLC | RLC00305 | 0.909090909090909 | 11 | 1008 | 1009.63636363636 |
| RLC | RLC00367 | 0.9 | 10 | 1616.5 | 1404.8 |
| RLG | RLG00287 | 0.846153846153846 | 13 | 1173 | 1101.38461538462 |
| RLG | RLG00363 | 0.8 | 10 | 1442 | 1235.9 |

**Table S3.** 842 families with at least 10 copies in the B73 genome that lack coding representatives.

| sup | fam | propAuton | famsize | medianORF | meanORF |
|---|---|---|---|---|---|

| | | | | | |
|---|---|---|---|---|---|
| DHH | DHH00001 | 0 | 5946 | 69 | 170.178694158076 |
| DHH | DHH00003 | 0 | 949 | 541 | 521.511956521739 |
| DHH | DHH00004 | 0 | 1611 | 369 | 347.998137802607 |
| DHH | DHH00005 | 0 | 265 | 221 | 295.777777777778 |
| DHH | DHH00006 | 0 | 1013 | 79 | 77.5364705882353 |
| DHH | DHH00007 | 0 | 403 | 202.5 | 238.27380952381 |
| DHH | DHH00008 | 0 | 738 | 739 | 651.770065075922 |
| DHH | DHH00009 | 0 | 283 | 250 | 276.218637992832 |
| DHH | DHH00011 | 0 | 81 | 225 | 266.012345679012 |
| DHH | DHH00012 | 0 | 164 | 226.5 | 248.440789473684 |
| DHH | DHH00013 | 0 | 156 | 67 | 145.893203883495 |
| DHH | DHH00014 | 0 | 90 | 402 | 446.134831460674 |
| DHH | DHH00015 | 0 | 119 | 68 | 129.794392523364 |
| DHH | DHH00016 | 0 | 149 | 61 | 90.4491525423729 |
| DHH | DHH00017 | 0 | 29 | 331 | 325.655172413793 |
| DHH | DHH00018 | 0 | 62 | 988 | 890.947368421053 |
| DHH | DHH00019 | 0 | 110 | 44.5 | 133.735294117647 |
| DHH | DHH00020 | 0 | 25 | 232 | 306 |
| DHH | DHH00021 | 0 | 51 | 388 | 413.392156862745 |
| DHH | DHH00022 | 0 | 35 | 192 | 281.318181818182 |
| DHH | DHH00023 | 0 | 46 | 253 | 227.673913043478 |
| DHH | DHH00024 | 0 | 40 | 49 | 74.4054054054054 |
| DHH | DHH00025 | 0 | 63 | 121.5 | 258.648148148148 |
| DHH | DHH00026 | 0 | 36 | 135 | 149.852941176471 |
| DHH | DHH00027 | 0 | 14 | 64 | 203 |
| DHH | DHH00028 | 0 | 12 | 218.5 | 319.416666666667 |
| DHH | DHH00029 | 0 | 75 | 179 | 242.808219178082 |
| DHH | DHH00030 | 0 | 18 | 322 | 269.470588235294 |
| DHH | DHH00032 | 0 | 68 | 197 | 239.754098360656 |
| DHH | DHH00036 | 0 | 21 | 1303 | 1193.04761904762 |
| DHH | DHH00037 | 0 | 16 | 221 | 281.933333333333 |
| DHH | DHH00038 | 0 | 11 | 142 | 214.555555555556 |
| DHH | DHH00040 | 0 | 16 | 306 | 374.4 |
| DHH | DHH00041 | 0 | 16 | 128.5 | 184 |
| DHH | DHH00043 | 0 | 18 | 146.5 | 173.777777777778 |
| DHH | DHH00044 | 0 | 15 | 109.5 | 130.214285714286 |
| DHH | DHH00045 | 0 | 10 | 81 | 147.714285714286 |
| DHH | DHH00048 | 0 | 24 | 146 | 151.434782608696 |
| DHH | DHH00049 | 0 | 17 | 362 | 318.714285714286 |
| DHH | DHH00050 | 0 | 15 | 54 | 120.142857142857 |
| DHH | DHH00052 | 0 | 13 | 555 | 478.666666666667 |
| DHH | DHH00059 | 0 | 11 | 83.5 | 93.5 |
| DHH | DHH00063 | 0 | 10 | 122 | 94.1111111111111 |
| DTA | DTA00003 | 0 | 12 | 74 | 74 |
| DTA | DTA00006 | 0 | 12 | 306 | 308.75 |
| DTA | DTA00016 | 0 | 10 | 315 | 322 |
| DTA | DTA00025 | 0 | 13 | 445 | 405.692307692308 |
| DTA | DTA00029 | 0 | 12 | 307.5 | 386.166666666667 |
| DTA | DTA00030 | 0 | 16 | 345 | 383.8125 |
| DTA | DTA00040 | 0 | 36 | 36 | 36.9285714285714 |
| DTA | DTA00045 | 0 | 23 | 43 | 46.25 |
| DTA | DTA00046 | 0 | 19 | 27 | 42.8333333333333 |
| DTA | DTA00049 | 0 | 38 | 30 | 58.4285714285714 |
| DTA | DTA00051 | 0 | 42 | 39.5 | 86.6666666666667 |
| DTA | DTA00059 | 0 | 12 | 51 | 48.1111111111111 |
| DTA | DTA00060 | 0 | 120 | 33 | 131.263157894737 |

| | | | | | |
|---|---|---|---|---|---|
| DTA | DTA00061 | 0 | 54 | 20 | 54.8 |
| DTA | DTA00062 | 0 | 10 | 131 | 131 |
| DTA | DTA00064 | 0 | 19 | 104 | 161.333333333333 |
| DTA | DTA00065 | 0 | 19 | 37 | 42.5 |
| DTA | DTA00066 | 0 | 35 | 29.5 | 33.7 |
| DTA | DTA00073 | 0 | 148 | 49.5 | 46.25 |
| DTA | DTA00076 | 0 | 55 | 42 | 42 |
| DTA | DTA00077 | 0 | 15 | 433 | 432.727272727273 |
| DTA | DTA00078 | 0 | 26 | 35 | 30.625 |
| DTA | DTA00080 | 0 | 72 | 106 | 129.1875 |
| DTA | DTA00085 | 0 | 15 | 417 | 455.6 |
| DTA | DTA00089 | 0 | 19 | 87.5 | 101.75 |
| DTA | DTA00090 | 0 | 23 | 84 | 169.285714285714 |
| DTA | DTA00091 | 0 | 30 | 86 | 95.6923076923077 |
| DTA | DTA00092 | 0 | 70 | 26 | 102.454545454545 |
| DTA | DTA00099 | 0 | 19 | 65 | 88.4 |
| DTA | DTA00102 | 0 | 157 | 29.5 | 37.5853658536585 |
| DTA | DTA00104 | 0 | 36 | 90 | 286 |
| DTA | DTA00106 | 0 | 31 | 58.5 | 134.230769230769 |
| DTA | DTA00114 | 0 | 147 | 65 | 79.5365853658537 |
| DTA | DTA00115 | 0 | 12 | 94.5 | 219.25 |
| DTA | DTA00117 | 0 | 234 | 31.5 | 40.3809523809524 |
| DTA | DTA00118 | 0 | 11 | 33 | 355.333333333333 |
| DTA | DTA00119 | 0 | 80 | NA | NaN |
| DTA | DTA00120 | 0 | 25 | 88 | 137.714285714286 |
| DTA | DTA00124 | 0 | 13 | 41 | 41 |
| DTA | DTA00126 | 0 | 20 | 24 | 24 |
| DTA | DTA00131 | 0 | 46 | 88 | 101.555555555556 |
| DTA | DTA00133 | 0 | 14 | 82 | 95 |
| DTA | DTA00135 | 0 | 132 | 56 | 53.8064516129032 |
| DTA | DTA00136 | 0 | 40 | 131 | 156.782608695652 |
| DTA | DTA00138 | 0 | 15 | 78 | 95 |
| DTA | DTA00141 | 0 | 192 | 43 | 50.8 |
| DTA | DTA00142 | 0 | 16 | 47 | 51 |
| DTA | DTA00143 | 0 | 35 | 36 | 42.5 |
| DTA | DTA00144 | 0 | 24 | 48.5 | 54 |
| DTA | DTA00148 | 0 | 10 | 58 | 118.8 |
| DTA | DTA00149 | 0 | 58 | 20 | 47.3529411764706 |
| DTA | DTA00151 | 0 | 20 | 47 | 97.3333333333333 |
| DTA | DTA00152 | 0 | 17 | 60 | 56.75 |
| DTA | DTA00153 | 0 | 11 | 19 | 19 |
| DTA | DTA00154 | 0 | 14 | 609.5 | 512.25 |
| DTA | DTA00155 | 0 | 27 | 87 | 115.117647058824 |
| DTA | DTA00156 | 0 | 22 | 55 | 52.4 |
| DTA | DTA00159 | 0 | 11 | 110 | 101.142857142857 |
| DTA | DTA00164 | 0 | 10 | 69 | 146.285714285714 |
| DTA | DTA00167 | 0 | 11 | 254 | 254 |
| DTA | DTA00169 | 0 | 12 | 41 | 40.3333333333333 |
| DTA | DTA00171 | 0 | 38 | 38 | 90.5 |
| DTA | DTA00172 | 0 | 263 | 25 | 26.6338028169014 |
| DTA | DTA00174 | 0 | 18 | 21 | 31 |
| DTA | DTA00179 | 0 | 19 | 25 | 25 |
| DTA | DTA00180 | 0 | 16 | NA | NaN |
| DTA | DTA00186 | 0 | 46 | 23 | 26.2 |
| DTA | DTA00191 | 0 | 17 | 20.5 | 20.5 |
| DTA | DTA00192 | 0 | 84 | 76 | 81.5806451612903 |
| DTA | DTA00194 | 0 | 56 | 61 | 64.6666666666667 |

| | | | | | |
|---|---|---|---|---|---|
| DTA | DTA00199 | 0 | 13 | 121.5 | 231.5 |
| DTA | DTA00204 | 0 | 68 | 37.5 | 37.5 |
| DTA | DTA00210 | 0 | 88 | NA | NaN |
| DTA | DTA00217 | 0 | 77 | 22 | 35.9 |
| DTA | DTA00219 | 0 | 11 | NA | NaN |
| DTA | DTA00224 | 0 | 13 | 84 | 96 |
| DTA | DTA00227 | 0 | 13 | 29 | 30.6 |
| DTA | DTA00238 | 0 | 19 | NA | NaN |
| DTA | DTA00239 | 0 | 17 | 33.5 | 33.5 |
| DTA | DTA00242 | 0 | 11 | 82.5 | 72 |
| DTA | DTA00243 | 0 | 13 | NA | NaN |
| DTA | DTA00245 | 0 | 28 | 138 | 297.8 |
| DTA | DTA00249 | 0 | 11 | NA | NaN |
| DTA | DTA00252 | 0 | 89 | 65 | 53.551724137931 |
| DTA | DTA00253 | 0 | 34 | 28 | 58.48 |
| DTA | DTA00256 | 0 | 15 | 461.5 | 406.166666666667 |
| DTA | DTA00267 | 0 | 11 | 23 | 37.1428571428571 |
| DTA | DTA00283 | 0 | 17 | 33 | 34.3333333333333 |
| DTA | DTA00286 | 0 | 17 | 27 | 30.9 |
| DTA | DTA00289 | 0 | 11 | 26 | 71.8 |
| DTA | DTA00291 | 0 | 42 | 39 | 50.9230769230769 |
| DTA | DTA00293 | 0 | 29 | 64 | 53.1578947368421 |
| DTA | DTA00295 | 0 | 16 | 28 | 25.3333333333333 |
| DTA | DTA00299 | 0 | 23 | NA | NaN |
| DTA | DTA00301 | 0 | 11 | 33 | 105.4 |
| DTA | DTA00307 | 0 | 62 | 25 | 30.2 |
| DTA | DTA00312 | 0 | 18 | 59 | 72.5 |
| DTA | DTA00313 | 0 | 11 | NA | NaN |
| DTA | DTA00323 | 0 | 16 | 25 | 34.5 |
| DTA | DTA00339 | 0 | 12 | 45.5 | 45.5 |
| DTA | DTA00345 | 0 | 16 | 22 | 22 |
| DTA | DTA00346 | 0 | 11 | 48 | 48 |
| DTA | DTA00348 | 0 | 15 | NA | NaN |
| DTA | DTA00383 | 0 | 79 | 29 | 28.1428571428571 |
| DTA | DTA13185 | 0 | 35 | 43 | 42.9333333333333 |
| DTC | DTC00109 | 0 | 10 | 39 | 44.4 |
| DTC | DTC00111 | 0 | 228 | 32 | 35.2894736842105 |
| DTC | DTC00112 | 0 | 67 | 32 | 40.2857142857143 |
| DTC | DTC00113 | 0 | 61 | 87.5 | 94.8125 |
| DTC | DTC00114 | 0 | 86 | 29 | 31.4444444444444 |
| DTC | DTC00115 | 0 | 20 | 37.5 | 58.75 |
| DTC | DTC00116 | 0 | 80 | 40.5 | 42.6666666666667 |
| DTC | DTC00118 | 0 | 79 | 24 | 35.0645161290323 |
| DTC | DTC00122 | 0 | 143 | 42 | 49.3695652173913 |
| DTC | DTC00126 | 0 | 10 | NA | NaN |
| DTC | DTC10850 | 0 | 15 | 30 | 32 |
| DTC | DTC12155 | 0 | 62 | 29 | 28.4090909090909 |
| DTC | DTC15669 | 0 | 14 | 35 | 43 |
| DTH | DTH00004 | 0 | 11 | 345 | 322.181818181818 |
| DTH | DTH00010 | 0 | 17 | 344 | 302.882352941176 |
| DTH | DTH00011 | 0 | 11 | 251 | 255.363636363636 |
| DTH | DTH00012 | 0 | 11 | 198 | 167.272727272727 |
| DTH | DTH00015 | 0 | 10 | 418 | 397.6 |
| DTH | DTH00024 | 0 | 17 | 186 | 279.058823529412 |
| DTH | DTH00032 | 0 | 11 | 219 | 238.636363636364 |
| DTH | DTH00037 | 0 | 97 | 35 | 40.2631578947368 |
| DTH | DTH00040 | 0 | 48 | 39 | 44.1666666666667 |

| | | | | | |
|---|---|---|---|---|---|
| DTH | DTH00041 | 0 | 21 | 69 | 67.8571428571429 |
| DTH | DTH00042 | 0 | 11 | 29 | 33 |
| DTH | DTH00043 | 0 | 15 | 46.5 | 46.5 |
| DTH | DTH00044 | 0 | 20 | 43 | 44.3333333333333 |
| DTH | DTH00046 | 0 | 13 | 43 | 52.25 |
| DTH | DTH00049 | 0 | 46 | 28 | 44.8333333333333 |
| DTH | DTH00050 | 0 | 16 | 36 | 47 |
| DTH | DTH00051 | 0 | 27 | 118 | 134.714285714286 |
| DTH | DTH00053 | 0 | 10 | NA | NaN |
| DTH | DTH00054 | 0 | 190 | 34 | 40.1351351351351 |
| DTH | DTH00057 | 0 | 181 | 67 | 58.7205882352941 |
| DTH | DTH00058 | 0 | 72 | 63 | 84.2 |
| DTH | DTH00062 | 0 | 29 | 27 | 26 |
| DTH | DTH00067 | 0 | 602 | 36 | 65.5 |
| DTH | DTH00071 | 0 | 11 | NA | NaN |
| DTH | DTH00073 | 0 | 122 | 42 | 128.777777777778 |
| DTH | DTH00076 | 0 | 29 | 45 | 50.0909090909091 |
| DTH | DTH00084 | 0 | 10 | 57 | 54 |
| DTH | DTH00090 | 0 | 214 | 32 | 35.4545454545455 |
| DTH | DTH00093 | 0 | 65 | 21 | 24 |
| DTH | DTH00098 | 0 | 495 | 26 | 29.0246305418719 |
| DTH | DTH00102 | 0 | 1291 | 44 | 40.5723684210526 |
| DTH | DTH00114 | 0 | 25 | 46 | 52.5 |
| DTH | DTH00118 | 0 | 397 | 37 | 41.1511627906977 |
| DTH | DTH00119 | 0 | 124 | 29 | 31.8235294117647 |
| DTH | DTH00127 | 0 | 164 | 25 | 30.6923076923077 |
| DTH | DTH00129 | 0 | 86 | 31 | 35.3636363636364 |
| DTH | DTH00135 | 0 | 38 | 34 | 40.2727272727273 |
| DTH | DTH00139 | 0 | 125 | 30 | 31.1315789473684 |
| DTH | DTH00145 | 0 | 19 | 46 | 46.3333333333333 |
| DTH | DTH00153 | 0 | 93 | 24 | 27.5 |
| DTH | DTH00154 | 0 | 606 | 33 | 34.3125 |
| DTH | DTH00160 | 0 | 196 | 35 | 37.2962962962963 |
| DTH | DTH00162 | 0 | 72 | 79 | 125.8 |
| DTH | DTH00163 | 0 | 968 | 38 | 34.8842105263158 |
| DTH | DTH00178 | 0 | 10 | 38.5 | 38.5 |
| DTH | DTH00180 | 0 | 16 | 34 | 33.1666666666667 |
| DTH | DTH00181 | 0 | 44 | 40.5 | 42.75 |
| DTH | DTH00184 | 0 | 26 | 42 | 39.0588235294118 |
| DTH | DTH00194 | 0 | 118 | 38 | 55.0618556701031 |
| DTH | DTH00198 | 0 | 17 | 54 | 54 |
| DTH | DTH00202 | 0 | 44 | 29 | 42.6 |
| DTH | DTH00220 | 0 | 10 | 42 | 40.7 |
| DTH | DTH00226 | 0 | 40 | 40 | 41 |
| DTH | DTH00233 | 0 | 292 | 33 | 38.804347826087 |
| DTH | DTH00241 | 0 | 35 | 39.5 | 39.5 |
| DTH | DTH00244 | 0 | 11 | 30 | 30 |
| DTH | DTH00273 | 0 | 11 | 34 | 84.6666666666667 |
| DTH | DTH00280 | 0 | 289 | 53 | 50.9074074074074 |
| DTH | DTH00292 | 0 | 10 | 116 | 116 |
| DTH | DTH00307 | 0 | 20 | NA | NaN |
| DTH | DTH00311 | 0 | 79 | 53 | 68.75 |
| DTH | DTH00314 | 0 | 10 | NA | NaN |
| DTH | DTH00320 | 0 | 95 | 23 | 27.4 |
| DTH | DTH00326 | 0 | 56 | 33 | 33 |
| DTH | DTH00327 | 0 | 481 | 40 | 60.0769230769231 |
| DTH | DTH00335 | 0 | 32 | 21 | 21.7272727272727 |

| | | | | | |
|---|---|---|---|---|---|
| DTH | DTH00337 | 0 | 75 | 32 | 64.4 |
| DTH | DTH00340 | 0 | 27 | 49 | 47.9333333333333 |
| DTH | DTH00347 | 0 | 14 | 124 | 110.2 |
| DTH | DTH00351 | 0 | 153 | 26.5 | 32.25 |
| DTH | DTH00355 | 0 | 67 | 44.5 | 47.875 |
| DTH | DTH00361 | 0 | 85 | 27 | 36.125 |
| DTH | DTH00364 | 0 | 76 | 43 | 51.0869565217391 |
| DTH | DTH00367 | 0 | 10 | 60.5 | 60.5 |
| DTH | DTH00382 | 0 | 13 | 24 | 24.5 |
| DTH | DTH00385 | 0 | 82 | 22.5 | 28.85 |
| DTH | DTH00389 | 0 | 212 | 26 | 34.48 |
| DTH | DTH00409 | 0 | 748 | 27 | 55.68 |
| DTH | DTH00410 | 0 | 76 | 20 | 30.6666666666667 |
| DTH | DTH00412 | 0 | 52 | 215.5 | 178.666666666667 |
| DTH | DTH00418 | 0 | 111 | 39 | 44.4705882352941 |
| DTH | DTH00431 | 0 | 74 | 43 | 42.8666666666667 |
| DTH | DTH00434 | 0 | 347 | 27 | 44.4545454545455 |
| DTH | DTH00437 | 0 | 152 | 49 | 60.1071428571429 |
| DTH | DTH00442 | 0 | 74 | 36 | 55 |
| DTH | DTH00453 | 0 | 18 | 31.5 | 31.5 |
| DTH | DTH00460 | 0 | 81 | 37 | 30.4 |
| DTH | DTH00463 | 0 | 18 | 33 | 33 |
| DTH | DTH00471 | 0 | 42 | 32 | 99.3333333333333 |
| DTH | DTH00485 | 0 | 58 | 37.5 | 40 |
| DTH | DTH00486 | 0 | 16 | 21 | 21 |
| DTH | DTH00489 | 0 | 99 | 23 | 27.1428571428571 |
| DTH | DTH10044 | 0 | 86 | 40 | 41.75 |
| DTH | DTH10047 | 0 | 169 | 30.5 | 42.9375 |
| DTH | DTH10048 | 0 | 13 | 52.5 | 50.75 |
| DTH | DTH10060 | 0 | 11 | 34.5 | 37.25 |
| DTH | DTH10064 | 0 | 28 | 20 | 20 |
| DTH | DTH10085 | 0 | 20 | 44 | 53.5714285714286 |
| DTH | DTH10107 | 0 | 290 | 44 | 45.2631578947368 |
| DTH | DTH10113 | 0 | 37 | 45 | 45 |
| DTH | DTH10153 | 0 | 61 | 43 | 43.5454545454545 |
| DTH | DTH10158 | 0 | 98 | 29 | 55.4090909090909 |
| DTH | DTH10176 | 0 | 146 | 32 | 38.9245283018868 |
| DTH | DTH10182 | 0 | 26 | 27.5 | 46 |
| DTH | DTH10187 | 0 | 80 | 29 | 29.1153846153846 |
| DTH | DTH10226 | 0 | 15 | 31 | 33 |
| DTH | DTH10230 | 0 | 107 | 28 | 30.3888888888889 |
| DTH | DTH10239 | 0 | 25 | 41 | 43.25 |
| DTH | DTH10248 | 0 | 21 | 26.5 | 33 |
| DTH | DTH10268 | 0 | 516 | 46 | 42.5354838709677 |
| DTH | DTH10298 | 0 | 21 | 33.5 | 33.8333333333333 |
| DTH | DTH10310 | 0 | 106 | 42 | 40.4390243902439 |
| DTH | DTH10314 | 0 | 13 | 44 | 46.75 |
| DTH | DTH10328 | 0 | 82 | 40.5 | 37.9705882352941 |
| DTH | DTH10339 | 0 | 13 | 38 | 36.5555555555556 |
| DTH | DTH10363 | 0 | 1053 | 41 | 39.4405594405594 |
| DTH | DTH10364 | 0 | 36 | 34 | 41.7857142857143 |
| DTH | DTH10379 | 0 | 42 | 35 | 36.6153846153846 |
| DTH | DTH10380 | 0 | 33 | 31.5 | 36.7777777777778 |
| DTH | DTH10388 | 0 | 43 | 41.5 | 52.75 |
| DTH | DTH10391 | 0 | 13 | 26 | 33.3333333333333 |
| DTH | DTH10406 | 0 | 20 | 28 | 38 |
| DTH | DTH10440 | 0 | 14 | 37 | 37 |

| | | | | | |
|---|---|---|---|---|---|
| DTH | DTH10442 | 0 | 25 | 36 | 36.8571428571429 |
| DTH | DTH10444 | 0 | 959 | 31 | 31.5714285714286 |
| DTH | DTH10445 | 0 | 312 | 30 | 52.3333333333333 |
| DTH | DTH10454 | 0 | 44 | 45 | 52.8421052631579 |
| DTH | DTH10481 | 0 | 16 | 38 | 40.7777777777778 |
| DTH | DTH10483 | 0 | 180 | 27 | 37.6727272727273 |
| DTH | DTH10494 | 0 | 17 | 39 | 43.4285714285714 |
| DTH | DTH10560 | 0 | 30 | 42.5 | 48.5 |
| DTH | DTH10573 | 0 | 104 | 35 | 36.3103448275862 |
| DTH | DTH10659 | 0 | 10 | NA | NaN |
| DTH | DTH10699 | 0 | 10 | 32 | 30.4 |
| DTH | DTH10700 | 0 | 368 | 33 | 33.4689655172414 |
| DTH | DTH10715 | 0 | 86 | 34 | 33.5471698113208 |
| DTH | DTH10741 | 0 | 10 | 31 | 31 |
| DTH | DTH10775 | 0 | 910 | 134 | 128.749152542373 |
| DTH | DTH10781 | 0 | 12 | 44.5 | 42.25 |
| DTH | DTH10818 | 0 | 14 | 31 | 31 |
| DTH | DTH10852 | 0 | 15 | 34 | 34 |
| DTH | DTH10855 | 0 | 659 | 28 | 29.8343558282209 |
| DTH | DTH10856 | 0 | 126 | 28 | 31.3870967741935 |
| DTH | DTH10946 | 0 | 179 | 25 | 30.0425531914894 |
| DTH | DTH11074 | 0 | 33 | 35 | 41.1111111111111 |
| DTH | DTH11101 | 0 | 409 | 21 | 26.9003984063745 |
| DTH | DTH11152 | 0 | 1624 | 34 | 54.9435431537962 |
| DTH | DTH11199 | 0 | 93 | 52 | 58.3793103448276 |
| DTH | DTH11200 | 0 | 52 | 50 | 58.7894736842105 |
| DTH | DTH11203 | 0 | 513 | 74 | 78 |
| DTH | DTH11206 | 0 | 176 | 135.5 | 141.850574712644 |
| DTH | DTH11207 | 0 | 245 | 123 | 118.725738396624 |
| DTH | DTH11208 | 0 | 187 | 38 | 43.7260273972603 |
| DTH | DTH11209 | 0 | 107 | 44 | 53.6944444444444 |
| DTH | DTH11211 | 0 | 13 | 38 | 44 |
| DTH | DTH11238 | 0 | 894 | 38 | 45.7435897435897 |
| DTH | DTH11239 | 0 | 1029 | 37 | 42.6699029126214 |
| DTH | DTH11253 | 0 | 30 | 20 | 20 |
| DTH | DTH11350 | 0 | 75 | 46 | 52.8125 |
| DTH | DTH11351 | 0 | 29 | 19 | 49.25 |
| DTH | DTH11388 | 0 | 16 | 31.5 | 37.875 |
| DTH | DTH11403 | 0 | 79 | 44 | 42.4210526315789 |
| DTH | DTH11404 | 0 | 10 | 37.5 | 37 |
| DTH | DTH11419 | 0 | 172 | 45 | 41.1692307692308 |
| DTH | DTH11440 | 0 | 10 | 49 | 49 |
| DTH | DTH11441 | 0 | 28 | 25 | 27.3333333333333 |
| DTH | DTH11449 | 0 | 935 | 90 | 80.3986784140969 |
| DTH | DTH11450 | 0 | 449 | 90 | 79.4233409610984 |
| DTH | DTH11457 | 0 | 64 | 62 | 69.7924528301887 |
| DTH | DTH11481 | 0 | 35 | 21.5 | 21.5 |
| DTH | DTH11482 | 0 | 24 | 29.5 | 28 |
| DTH | DTH11541 | 0 | 464 | 83 | 77.7443365695793 |
| DTH | DTH11542 | 0 | 567 | 77 | 78.0234741784038 |
| DTH | DTH11594 | 0 | 65 | 19 | 19.2727272727273 |
| DTH | DTH11602 | 0 | 73 | 38 | 39.4358974358974 |
| DTH | DTH11614 | 0 | 14 | 53 | 53.2 |
| DTH | DTH11708 | 0 | 1556 | 111 | 105.549578742709 |
| DTH | DTH11714 | 0 | 90 | 36 | 45.3829787234043 |
| DTH | DTH11715 | 0 | 14 | 46 | 44.4 |
| DTH | DTH11798 | 0 | 14 | 54 | 54 |

| | | | | | |
|---|---|---|---|---|---|
| DTH | DTH11817 | 0 | 346 | 33 | 33.7368421052632 |
| DTH | DTH11830 | 0 | 140 | 25 | 30.4347826086957 |
| DTH | DTH11834 | 0 | 71 | 26 | 26.0909090909091 |
| DTH | DTH11854 | 0 | 104 | 115 | 113.7 |
| DTH | DTH11856 | 0 | 60 | 19 | 30.8571428571429 |
| DTH | DTH11862 | 0 | 53 | 44 | 49.1176470588235 |
| DTH | DTH11932 | 0 | 44 | 42 | 43.1111111111111 |
| DTH | DTH11948 | 0 | 17 | 33 | 30.6666666666667 |
| DTH | DTH11949 | 0 | 14 | 40 | 37 |
| DTH | DTH12058 | 0 | 11 | NA | NaN |
| DTH | DTH12059 | 0 | 13 | 20 | 20 |
| DTH | DTH12181 | 0 | 158 | 29 | 29 |
| DTH | DTH12188 | 0 | 16 | 56 | 54.3333333333333 |
| DTH | DTH12191 | 0 | 43 | NA | NaN |
| DTH | DTH12217 | 0 | 58 | 34 | 109.2 |
| DTH | DTH12228 | 0 | 26 | 62 | 62 |
| DTH | DTH12232 | 0 | 109 | 67.5 | 70.3448275862069 |
| DTH | DTH12250 | 0 | 215 | 63 | 51.8571428571429 |
| DTH | DTH12251 | 0 | 132 | 60 | 66.7368421052632 |
| DTH | DTH12258 | 0 | 2623 | 58 | 69.0553449583017 |
| DTH | DTH12289 | 0 | 13 | 22 | 30.6666666666667 |
| DTH | DTH12290 | 0 | 46 | 106 | 76.7179487179487 |
| DTH | DTH12298 | 0 | 203 | 34 | 37.8190476190476 |
| DTH | DTH12305 | 0 | 50 | 24 | 59.7142857142857 |
| DTH | DTH12326 | 0 | 267 | 24 | 45.75 |
| DTH | DTH12381 | 0 | 108 | 26 | 27.4117647058824 |
| DTH | DTH12383 | 0 | 10 | 22 | 45.3333333333333 |
| DTH | DTH12388 | 0 | 110 | 29 | 34.3 |
| DTH | DTH12389 | 0 | 103 | 29 | 38.304347826087 |
| DTH | DTH12390 | 0 | 296 | 27.5 | 28.2 |
| DTH | DTH12393 | 0 | 153 | 40 | 110.3 |
| DTH | DTH12412 | 0 | 492 | 34 | 39.0961538461538 |
| DTH | DTH12419 | 0 | 24 | 73 | 58.6666666666667 |
| DTH | DTH12507 | 0 | 113 | 41 | 46.0491803278689 |
| DTH | DTH12534 | 0 | 259 | 43 | 47.9195402298851 |
| DTH | DTH12537 | 0 | 22 | 35 | 41.125 |
| DTH | DTH12555 | 0 | 18 | 25 | 28.25 |
| DTH | DTH12617 | 0 | 34 | 23 | 25.875 |
| DTH | DTH12628 | 0 | 24 | 37 | 56.5384615384615 |
| DTH | DTH12645 | 0 | 17 | 77 | 76 |
| DTH | DTH12694 | 0 | 14 | 43 | 42.6 |
| DTH | DTH12697 | 0 | 21 | 38 | 44.1111111111111 |
| DTH | DTH12711 | 0 | 19 | 38 | 33.3333333333333 |
| DTH | DTH12718 | 0 | 186 | 30.5 | 151.375 |
| DTH | DTH12769 | 0 | 211 | 63.5 | 64.551724137931 |
| DTH | DTH12801 | 0 | 232 | 154 | 144.188340807175 |
| DTH | DTH12838 | 0 | 350 | 81.5 | 77.9065040650407 |
| DTH | DTH12839 | 0 | 90 | 72.5 | 75.3103448275862 |
| DTH | DTH12973 | 0 | 325 | 28 | 40.7692307692308 |
| DTH | DTH12996 | 0 | 1844 | 32 | 37.4453870625663 |
| DTH | DTH12997 | 0 | 280 | 31 | 37.7747747747748 |
| DTH | DTH13012 | 0 | 22 | 63.5 | 63.5 |
| DTH | DTH13063 | 0 | 32 | 41 | 48.1111111111111 |
| DTH | DTH13110 | 0 | 577 | 55 | 60.1994884910486 |
| DTH | DTH13117 | 0 | 450 | 121 | 126.732426303855 |
| DTH | DTH13200 | 0 | 147 | 30 | 36.2727272727273 |
| DTH | DTH13235 | 0 | 84 | 190 | 186.024096385542 |

| | | | | | |
|---|---|---|---|---|---|
| DTH | DTH13261 | 0 | 28 | 27 | 33.2 |
| DTH | DTH13439 | 0 | 13 | 24 | 25.6666666666667 |
| DTH | DTH13583 | 0 | 1462 | 41 | 48.6263383297645 |
| DTH | DTH13597 | 0 | 21 | 57 | 57 |
| DTH | DTH13636 | 0 | 22 | 72 | 71.8888888888889 |
| DTH | DTH13647 | 0 | 39 | 37 | 40.125 |
| DTH | DTH13657 | 0 | 74 | 38.5 | 46.1785714285714 |
| DTH | DTH13748 | 0 | 10 | 65 | 55 |
| DTH | DTH13854 | 0 | 120 | 47 | 59.5116279069767 |
| DTH | DTH13942 | 0 | 166 | 62 | 68.2527472527473 |
| DTH | DTH13989 | 0 | 33 | 45 | 47.1176470588235 |
| DTH | DTH13990 | 0 | 30 | 46 | 47.2222222222222 |
| DTH | DTH14072 | 0 | 14 | 21 | 21 |
| DTH | DTH14130 | 0 | 11 | 28 | 28 |
| DTH | DTH14236 | 0 | 234 | 25 | 25.8771929824561 |
| DTH | DTH14279 | 0 | 32 | 58 | 56.3636363636364 |
| DTH | DTH14804 | 0 | 1175 | 118 | 118.327630453379 |
| DTH | DTH14898 | 0 | 121 | 80.5 | 84.7307692307692 |
| DTH | DTH15065 | 0 | 78 | 28 | 37.8571428571429 |
| DTH | DTH15116 | 0 | 12 | 45 | 55.3333333333333 |
| DTH | DTH15132 | 0 | 235 | 42 | 47.1702127659574 |
| DTH | DTH15139 | 0 | 12 | 24 | 26.3333333333333 |
| DTH | DTH15158 | 0 | 79 | 56 | 55.5348837209302 |
| DTH | DTH15299 | 0 | 255 | 67 | 79.0152671755725 |
| DTH | DTH15300 | 0 | 100 | 64 | 75.1636363636364 |
| DTH | DTH15301 | 0 | 45 | 61.5 | 83.7 |
| DTH | DTH16098 | 0 | 90 | 57 | 79.2631578947368 |
| DTH | DTH16099 | 0 | 56 | 61 | 87.6 |
| DTH | DTH16100 | 0 | 40 | 105 | 92.7777777777778 |
| DTH | DTH16163 | 0 | 12 | NA | NaN |
| DTH | DTH16174 | 0 | 211 | 50 | 53.9 |
| DTH | DTH16203 | 0 | 29 | 27.5 | 27.5 |
| DTH | DTH16233 | 0 | 867 | 30 | 38.0476190476191 |
| DTH | DTH16329 | 0 | 598 | 70 | 87.0020325203252 |
| DTH | DTH16351 | 0 | 11 | 106 | 97 |
| DTH | DTH16373 | 0 | 611 | 74 | 94.4440559440559 |
| DTH | DTH16443 | 0 | 98 | 28 | 39.1282051282051 |
| DTH | DTH16563 | 0 | 263 | 121 | 132.171875 |
| DTH | DTH16728 | 0 | 1322 | 30 | 51.3277693474962 |
| DTH | DTH16758 | 0 | 768 | 31 | 37.1825396825397 |
| DTH | DTH16778 | 0 | 561 | 91 | 98.0576923076923 |
| DTH | DTH16779 | 0 | 108 | 90 | 90.890625 |
| DTH | DTH16783 | 0 | 149 | 39 | 44.5581395348837 |
| DTH | DTH16799 | 0 | 317 | 172 | 154.546391752577 |
| DTH | DTH16801 | 0 | 14 | 22 | 22 |
| DTM | DTM00118 | 0 | 19 | 22 | 22.3 |
| DTM | DTM00257 | 0 | 376 | 50 | 48.3404255319149 |
| DTM | DTM00266 | 0 | 47 | 64 | 470.333333333333 |
| DTM | DTM00299 | 0 | 22 | 26 | 26 |
| DTM | DTM00460 | 0 | 41 | 34 | 34 |
| DTM | DTM00473 | 0 | 30 | 51 | 54.8235294117647 |
| DTM | DTM00555 | 0 | 37 | 24 | 24 |
| DTM | DTM00796 | 0 | 16 | 50.5 | 46.4 |
| DTM | DTM00800 | 0 | 17 | 41 | 41 |
| DTM | DTM01654 | 0 | 13 | 61 | 50.2307692307692 |
| DTM | DTM02275 | 0 | 10 | NA | NaN |
| DTM | DTM13640 | 0 | 178 | 59 | 54.7647058823529 |

| | | | | | |
|---|---|---|---|---|---|
| DTT | DTT00008 | 0 | 32 | 23 | 30.25 |
| DTT | DTT00009 | 0 | 452 | 27 | 35.5526315789474 |
| DTT | DTT00010 | 0 | 932 | 26 | 29.015037593985 |
| DTT | DTT00011 | 0 | 519 | 30 | 37.6666666666667 |
| DTT | DTT00012 | 0 | 395 | 34 | 190 |
| DTT | DTT00013 | 0 | 137 | 28.5 | 37.375 |
| DTT | DTT00015 | 0 | 234 | 26 | 37.4242424242424 |
| DTT | DTT00016 | 0 | 632 | 32.5 | 51.5634920634921 |
| DTT | DTT00019 | 0 | 12 | 21 | 21 |
| DTT | DTT00022 | 0 | 232 | 26 | 35.5689655172414 |
| DTT | DTT00024 | 0 | 211 | 35 | 45.5689655172414 |
| DTT | DTT00026 | 0 | 1409 | 32 | 39.6736111111111 |
| DTT | DTT00027 | 0 | 316 | 32 | 38.4411764705882 |
| DTT | DTT00031 | 0 | 28 | 22.5 | 24.8333333333333 |
| DTT | DTT00032 | 0 | 291 | 40 | 44.9076923076923 |
| DTT | DTT00033 | 0 | 53 | 38 | 34.4 |
| DTT | DTT00034 | 0 | 80 | 29.5 | 34.7 |
| DTT | DTT00038 | 0 | 15 | 29 | 29 |
| DTT | DTT00040 | 0 | 13 | 28 | 28 |
| DTT | DTT00041 | 0 | 57 | 37 | 32.6666666666667 |
| DTT | DTT00042 | 0 | 429 | 30 | 42.1818181818182 |
| DTT | DTT00044 | 0 | 127 | 28 | 30.4375 |
| DTT | DTT00046 | 0 | 95 | 38.5 | 46 |
| DTT | DTT00049 | 0 | 23 | 64.5 | 61.25 |
| DTT | DTT00053 | 0 | 140 | 29 | 34.5714285714286 |
| DTT | DTT00054 | 0 | 28 | 22 | 22 |
| DTT | DTT00056 | 0 | 301 | 25 | 28.3863636363636 |
| DTT | DTT10003 | 0 | 47 | 39 | 42.9285714285714 |
| DTT | DTT10009 | 0 | 702 | 34.5 | 63.9333333333333 |
| DTT | DTT10033 | 0 | 52 | 38 | 38.5 |
| DTT | DTT10037 | 0 | 93 | 20 | 20 |
| DTT | DTT10038 | 0 | 74 | 26 | 26 |
| DTT | DTT10054 | 0 | 55 | 23 | 28.3333333333333 |
| DTT | DTT10062 | 0 | 273 | 39.5 | 37.5 |
| DTT | DTT10089 | 0 | 148 | 25 | 32 |
| DTT | DTT10101 | 0 | 128 | 23.5 | 38.25 |
| DTT | DTT10115 | 0 | 160 | 28.5 | 31.28125 |
| DTT | DTT10116 | 0 | 112 | 30 | 31.9459459459459 |
| DTT | DTT10125 | 0 | 32 | 25 | 25.4285714285714 |
| DTT | DTT10171 | 0 | 72 | 30 | 33.3333333333333 |
| DTT | DTT10172 | 0 | 73 | 24 | 28.5714285714286 |
| DTT | DTT10193 | 0 | 59 | 20 | 20 |
| DTT | DTT10253 | 0 | 29 | 49.5 | 46.6666666666667 |
| DTT | DTT10254 | 0 | 384 | 47 | 52.6157894736842 |
| DTT | DTT10260 | 0 | 125 | NA | NaN |
| DTT | DTT10319 | 0 | 26 | 29 | 28.6666666666667 |
| DTT | DTT10370 | 0 | 21 | 24 | 24 |
| DTT | DTT10404 | 0 | 13 | 25 | 29.7142857142857 |
| DTT | DTT10422 | 0 | 11 | 24 | 32.3333333333333 |
| DTT | DTT10423 | 0 | 18 | 38 | 38 |
| DTT | DTT10433 | 0 | 21 | 34.5 | 35 |
| DTT | DTT10449 | 0 | 11 | 39.5 | 39.5 |
| DTT | DTT10459 | 0 | 43 | 36 | 37.375 |
| DTT | DTT10475 | 0 | 148 | 24 | 27.9016393442623 |
| DTT | DTT10539 | 0 | 22 | 51 | 48.2857142857143 |
| DTT | DTT10540 | 0 | 13 | 44 | 37.4285714285714 |
| DTT | DTT10577 | 0 | 806 | 34 | 35.7627118644068 |

| | | | | | |
|---|---|---|---|---|---|
| DTT | DTT10584 | 0 | 16 | NA | NaN |
| DTT | DTT10594 | 0 | 15 | 55 | 55 |
| DTT | DTT10647 | 0 | 12 | NA | NaN |
| DTT | DTT10693 | 0 | 79 | 31 | 35.6666666666667 |
| DTT | DTT10706 | 0 | 204 | 22 | 22 |
| DTT | DTT10724 | 0 | 58 | 99 | 77.2727272727273 |
| DTT | DTT10738 | 0 | 51 | 24 | 29 |
| DTT | DTT10742 | 0 | 15 | 20 | 20 |
| DTT | DTT10797 | 0 | 889 | 29 | 27.8618421052632 |
| DTT | DTT10806 | 0 | 18 | 25 | 25 |
| DTT | DTT10833 | 0 | 9953 | 37 | 44.4548395123165 |
| DTT | DTT10841 | 0 | 28 | NA | NaN |
| DTT | DTT10864 | 0 | 113 | 41 | 41.6481481481481 |
| DTT | DTT10865 | 0 | 56 | 30 | 31 |
| DTT | DTT10873 | 0 | 122 | 26 | 33.3478260869565 |
| DTT | DTT10880 | 0 | 6151 | 33 | 37.9612289685443 |
| DTT | DTT10889 | 0 | 267 | 28 | 36.7959183673469 |
| DTT | DTT10892 | 0 | 69 | 41 | 46.3636363636364 |
| DTT | DTT10893 | 0 | 39 | 28.5 | 32.5 |
| DTT | DTT10906 | 0 | 19 | 24 | 24 |
| DTT | DTT10907 | 0 | 15 | 22 | 22 |
| DTT | DTT10927 | 0 | 277 | 25 | 31.3125 |
| DTT | DTT10937 | 0 | 25 | 38.5 | 38.5 |
| DTT | DTT10986 | 0 | 20 | 37.5 | 43 |
| DTT | DTT11051 | 0 | 51 | 34.5 | 39.0769230769231 |
| DTT | DTT11056 | 0 | 231 | 34 | 34.7121212121212 |
| DTT | DTT11071 | 0 | 16 | 30.5 | 30.5 |
| DTT | DTT11073 | 0 | 199 | 33 | 36 |
| DTT | DTT11075 | 0 | 41 | 29 | 38.4375 |
| DTT | DTT11090 | 0 | 169 | 49 | 48.7777777777778 |
| DTT | DTT11146 | 0 | 48 | 32 | 37.2 |
| DTT | DTT11147 | 0 | 12 | 45 | 41.6666666666667 |
| DTT | DTT11169 | 0 | 39 | 24 | 24 |
| DTT | DTT11170 | 0 | 51 | 47.5 | 47.5 |
| DTT | DTT11191 | 0 | 16 | 116 | 97.3333333333333 |
| DTT | DTT11230 | 0 | 53 | 35 | 37.5555555555556 |
| DTT | DTT11244 | 0 | 56 | 47 | 41.1333333333333 |
| DTT | DTT11258 | 0 | 15 | 33 | 30 |
| DTT | DTT11275 | 0 | 16 | 24 | 23.3333333333333 |
| DTT | DTT11278 | 0 | 50 | 30 | 41.3333333333333 |
| DTT | DTT11287 | 0 | 11 | 34 | 34.7777777777778 |
| DTT | DTT11292 | 0 | 73 | 22.5 | 25.6875 |
| DTT | DTT11306 | 0 | 21 | 33 | 33.2 |
| DTT | DTT11307 | 0 | 11 | 25 | 25 |
| DTT | DTT11338 | 0 | 296 | 59.5 | 61.9918032786885 |
| DTT | DTT11408 | 0 | 48 | 50 | 51 |
| DTT | DTT11421 | 0 | 14 | 100.5 | 92.5 |
| DTT | DTT11451 | 0 | 88 | 47.5 | 51.8 |
| DTT | DTT11452 | 0 | 10 | 48 | 48 |
| DTT | DTT11463 | 0 | 45 | 34 | 35.2 |
| DTT | DTT11547 | 0 | 1825 | 38 | 49.8983050847458 |
| DTT | DTT11587 | 0 | 98 | 28.5 | 31.875 |
| DTT | DTT11612 | 0 | 53 | 35 | 41.8666666666667 |
| DTT | DTT11681 | 0 | 44 | 44 | 39.6 |
| DTT | DTT11719 | 0 | 69 | 46 | 52.8181818181818 |
| DTT | DTT11766 | 0 | 11 | 39 | 39 |
| DTT | DTT11806 | 0 | 37 | 37 | 48.4285714285714 |

| | | | | | |
|---|---|---|---|---|---|
| DTT | DTT11836 | 0 | 16 | 58.5 | 58.5 |
| DTT | DTT12019 | 0 | 100 | 42.5 | 41.75 |
| DTT | DTT12061 | 0 | 170 | 39 | 36.6 |
| DTT | DTT12092 | 0 | 33 | 29 | 28 |
| DTT | DTT12107 | 0 | 55 | 53 | 60.8260869565217 |
| DTT | DTT12131 | 0 | 69 | 31.5 | 38.5 |
| DTT | DTT12147 | 0 | 50 | 35 | 42.9230769230769 |
| DTT | DTT12149 | 0 | 33 | 28 | 28.375 |
| DTT | DTT12151 | 0 | 43 | 23 | 26.08 |
| DTT | DTT12171 | 0 | 307 | 133 | 122.582644628099 |
| DTT | DTT12202 | 0 | 1062 | 35 | 42.6472081218274 |
| DTT | DTT12206 | 0 | 22 | 38 | 38 |
| DTT | DTT12283 | 0 | 470 | 38.5 | 43.3490566037736 |
| DTT | DTT12293 | 0 | 146 | 43 | 47.9354838709677 |
| DTT | DTT12378 | 0 | 21 | 40 | 40.3 |
| DTT | DTT12418 | 0 | 1304 | 138 | 132.157399486741 |
| DTT | DTT12506 | 0 | 1108 | 37 | 43.2792792792793 |
| DTT | DTT12510 | 0 | 51 | 22.5 | 22.5 |
| DTT | DTT12572 | 0 | 550 | 33 | 36.5084745762712 |
| DTT | DTT12632 | 0 | 144 | 53 | 58.6521739130435 |
| DTT | DTT12746 | 0 | 56 | 31 | 39.4285714285714 |
| DTT | DTT12754 | 0 | 51 | 23 | 22 |
| DTT | DTT12757 | 0 | 294 | 26 | 33.1 |
| DTT | DTT12762 | 0 | 108 | 50 | 49.6315789473684 |
| DTT | DTT12764 | 0 | 39 | 27.5 | 27.5 |
| DTT | DTT12820 | 0 | 11 | 123 | 121 |
| DTT | DTT12824 | 0 | 48 | 38 | 38.4 |
| DTT | DTT12825 | 0 | 19 | 33 | 35 |
| DTT | DTT12875 | 0 | 634 | 31 | 32.9934640522876 |
| DTT | DTT12876 | 0 | 247 | 29 | 34.0444444444444 |
| DTT | DTT12955 | 0 | 383 | 27 | 43.7407407407407 |
| DTT | DTT12967 | 0 | 172 | 28 | 36.7058823529412 |
| DTT | DTT12974 | 0 | 684 | 35 | 39.0595238095238 |
| DTT | DTT13041 | 0 | 4111 | 57 | 55.946103423161 |
| DTT | DTT13098 | 0 | 114 | 26 | 36.6153846153846 |
| DTT | DTT13262 | 0 | 64 | 32 | 33.125 |
| DTT | DTT13272 | 0 | 58 | 71 | 82.8571428571429 |
| DTT | DTT13542 | 0 | 21 | 31 | 30.75 |
| DTT | DTT13549 | 0 | 13 | 55 | 49.3333333333333 |
| DTT | DTT13551 | 0 | 18 | 20 | 20 |
| DTT | DTT13762 | 0 | 41 | 24 | 25.5555555555556 |
| DTT | DTT13827 | 0 | 16 | 47 | 47 |
| DTT | DTT13868 | 0 | 12 | 51 | 49.7 |
| DTT | DTT13979 | 0 | 117 | 29 | 35.1333333333333 |
| DTT | DTT14079 | 0 | 17 | NA | NaN |
| DTT | DTT14220 | 0 | 57 | 25 | 31.3333333333333 |
| DTT | DTT14352 | 0 | 20 | 24 | 29.2 |
| DTT | DTT14358 | 0 | 231 | 41 | 47.7560975609756 |
| DTT | DTT14774 | 0 | 17 | 30 | 30 |
| DTT | DTT14784 | 0 | 1391 | 27 | 32.1055555555556 |
| DTT | DTT15038 | 0 | 23 | 24 | 24 |
| DTT | DTT15227 | 0 | 14 | NA | NaN |
| DTT | DTT15264 | 0 | 248 | 19.5 | 26.2051282051282 |
| DTT | DTT15335 | 0 | 28 | 26.5 | 24.625 |
| DTT | DTT15443 | 0 | 143 | 35 | 38.6739130434783 |
| DTT | DTT15444 | 0 | 519 | 33.5 | 35.0967741935484 |
| DTT | DTT15534 | 0 | 48 | 42 | 45.6 |

| | | | | | |
|---|---|---|---|---|---|
| DTT | DTT15615 | 0 | 15 | 28.5 | 31 |
| DTT | DTT15785 | 0 | 30 | 27.5 | 31.25 |
| DTT | DTT15993 | 0 | 671 | 38 | 44.8076923076923 |
| DTT | DTT16190 | 0 | 3555 | 31 | 37.0521978021978 |
| DTT | DTT16489 | 0 | 414 | 25.5 | 25.125 |
| DTX | DTX10013 | 0 | 270 | 34 | 82.462962962963 |
| DTX | DTX10015 | 0 | 46 | 44.5 | 47.7777777777778 |
| DTX | DTX10020 | 0 | 94 | 23.5 | 28.4166666666667 |
| DTX | DTX10023 | 0 | 26 | 26 | 26.6666666666667 |
| DTX | DTX10031 | 0 | 100 | 70 | 124.4375 |
| DTX | DTX10032 | 0 | 83 | 71.5 | 71.7241379310345 |
| DTX | DTX10124 | 0 | 56 | 30 | 35.6428571428571 |
| DTX | DTX10131 | 0 | 120 | 27 | 31 |
| DTX | DTX10132 | 0 | 90 | 29 | 33.75 |
| DTX | DTX10177 | 0 | 2850 | 27 | 33.3927272727273 |
| DTX | DTX10207 | 0 | 108 | 39 | 43 |
| DTX | DTX10212 | 0 | 101 | 28 | 30.9375 |
| DTX | DTX10213 | 0 | 15 | NA | NaN |
| DTX | DTX10267 | 0 | 51 | 36 | 37.84 |
| DTX | DTX10308 | 0 | 105 | 40 | 43.4285714285714 |
| DTX | DTX10309 | 0 | 14 | 41 | 39 |
| DTX | DTX10416 | 0 | 19 | 32 | 36.4444444444444 |
| DTX | DTX10541 | 0 | 13 | 25 | 31.8 |
| DTX | DTX10542 | 0 | 26 | 31 | 40.6 |
| DTX | DTX10813 | 0 | 20 | 32 | 33.5454545454545 |
| DTX | DTX10824 | 0 | 38 | 25 | 25 |
| DTX | DTX10825 | 0 | 41 | 26 | 26 |
| DTX | DTX10840 | 0 | 243 | 31 | 34.0263157894737 |
| DTX | DTX10945 | 0 | 10 | 24 | 39.6666666666667 |
| DTX | DTX11005 | 0 | 33 | 25 | 35 |
| DTX | DTX11011 | 0 | 75 | 26 | 34.625 |
| DTX | DTX11123 | 0 | 49 | 39 | 50.4814814814815 |
| DTX | DTX11178 | 0 | 18 | 23 | 22.3333333333333 |
| DTX | DTX11265 | 0 | 21 | 47 | 49.6363636363636 |
| DTX | DTX11433 | 0 | 776 | 30 | 39.36 |
| DTX | DTX11622 | 0 | 14 | 40 | 42.3333333333333 |
| DTX | DTX11661 | 0 | 206 | 61 | 55.7796610169492 |
| DTX | DTX11816 | 0 | 62 | 26 | 27.25 |
| DTX | DTX11871 | 0 | 273 | 43 | 46.4565217391304 |
| DTX | DTX12196 | 0 | 26 | 33 | 33 |
| DTX | DTX12197 | 0 | 141 | 90 | 76.2238805970149 |
| DTX | DTX12438 | 0 | 23 | 56 | 54.5384615384615 |
| DTX | DTX12495 | 0 | 281 | 45 | 168.875 |
| DTX | DTX12496 | 0 | 195 | 26 | 49.6031746031746 |
| DTX | DTX12540 | 0 | 22 | 40 | 40 |
| DTX | DTX12562 | 0 | 18 | 49 | 62.4 |
| DTX | DTX12870 | 0 | 1297 | 33 | 38.7846364883402 |
| DTX | DTX12956 | 0 | 428 | 32 | 36.665306122449 |
| DTX | DTX12962 | 0 | 778 | 45 | 74.0625 |
| DTX | DTX13146 | 0 | 3356 | 39 | 49.6301476301476 |
| DTX | DTX13147 | 0 | 474 | 44 | 52.0334928229665 |
| DTX | DTX13328 | 0 | 708 | 41 | 51.4421768707483 |
| RIL | RIL00001 | 0 | 49 | 490 | 594.673469387755 |
| RIL | RIL00002 | 0 | 28 | 231 | 291.357142857143 |
| RIL | RIL00003 | 0 | 21 | 326 | 418.952380952381 |
| RIL | RIL00004 | 0 | 22 | 584.5 | 608.318181818182 |
| RIL | RIL00005 | 0 | 33 | 387 | 440.878787878788 |

| | | | | | |
|---|---|---|---|---|---|
| RIL | RIL00006 | 0 | 21 | 906 | 822.761904761905 |
| RIL | RIL00007 | 0 | 12 | 503.5 | 587.916666666667 |
| RIL | RIL00008 | 0 | 44 | 338 | 344.5 |
| RIL | RIL00009 | 0 | 27 | 658 | 628.576923076923 |
| RIL | RIL00013 | 0 | 10 | 244 | 274.5 |
| RIL | RIL00015 | 0 | 15 | 580 | 696 |
| RIL | RIL00016 | 0 | 18 | 431.5 | 522.055555555556 |
| RIL | RIL00017 | 0 | 16 | 586.5 | 618.375 |
| RIL | RIL00019 | 0 | 14 | 572 | 641.5 |
| RIL | RIL00020 | 0 | 10 | 958.5 | 771.9 |
| RIL | RIL00021 | 0 | 21 | 608 | 652.285714285714 |
| RIL | RIL00025 | 0 | 17 | 611 | 644.235294117647 |
| RIL | RIL00026 | 0 | 14 | 933 | 900 |
| RIL | RIL00028 | 0 | 11 | 636 | 750.454545454545 |
| RIT | RIT00001 | 0 | 169 | 280 | 304.109090909091 |
| RIT | RIT00002 | 0 | 129 | 159 | 246.258620689655 |
| RLC | RLC00097 | 0 | 43 | 390 | 401.139534883721 |
| RLC | RLC00116 | 0 | 34 | 339.5 | 394.625 |
| RLC | RLC00119 | 0 | 34 | 355.5 | 344.916666666667 |
| RLC | RLC00146 | 0 | 26 | 242.5 | 305.269230769231 |
| RLC | RLC00148 | 0 | 29 | 155 | 236.75 |
| RLC | RLC00154 | 0 | 26 | 61 | 104.923076923077 |
| RLC | RLC00182 | 0 | 22 | 980.5 | 966.727272727273 |
| RLC | RLC00184 | 0 | 23 | 75 | 97.7826086956522 |
| RLC | RLC00200 | 0 | 12 | 88 | 363 |
| RLC | RLC00218 | 0 | 17 | 327 | 402.764705882353 |
| RLC | RLC00228 | 0 | 16 | 365 | 378.2 |
| RLC | RLC00247 | 0 | 14 | 313.5 | 377.5 |
| RLC | RLC00248 | 0 | 15 | 326 | 369.4 |
| RLC | RLC00250 | 0 | 15 | 213 | 248.866666666667 |
| RLC | RLC00255 | 0 | 13 | 292 | 305.153846153846 |
| RLC | RLC00274 | 0 | 16 | 500 | 536.5 |
| RLC | RLC00279 | 0 | 13 | 579 | 538.769230769231 |
| RLC | RLC00293 | 0 | 11 | 513 | 478.545454545455 |
| RLC | RLC00311 | 0 | 11 | 276 | 356.727272727273 |
| RLC | RLC00340 | 0 | 12 | 660.5 | 745.5 |
| RLC | RLC00358 | 0 | 10 | 920 | 905.6 |
| RLC | RLC00375 | 0 | 10 | 471.5 | 787.9 |
| RLC | RLC00462 | 0 | 11 | 75 | 189.909090909091 |
| RLC | RLC01304 | 0 | 19 | 660 | 747.578947368421 |
| RLC | RLC02403 | 0 | 51 | 1064 | 974.901960784314 |
| RLC | RLC02882 | 0 | 33 | 683 | 554.272727272727 |
| RLC | RLC06245 | 0 | 45 | 546 | 495.644444444444 |
| RLC | RLC19829 | 0 | 129 | 728 | 678.573643410853 |
| RLG | RLG00053 | 0 | 121 | 381.5 | 392.708333333333 |
| RLG | RLG00077 | 0 | 60 | 42 | 124.529411764706 |
| RLG | RLG00084 | 0 | 51 | 59 | 87.8823529411765 |
| RLG | RLG00100 | 0 | 55 | 618 | 479.137254901961 |
| RLG | RLG00106 | 0 | 33 | 270 | 304.333333333333 |
| RLG | RLG00114 | 0 | 35 | 300 | 339.971428571429 |
| RLG | RLG00115 | 0 | 34 | 422 | 475.411764705882 |
| RLG | RLG00120 | 0 | 33 | 259 | 289.515151515152 |
| RLG | RLG00131 | 0 | 29 | 385 | 422.793103448276 |
| RLG | RLG00152 | 0 | 26 | 58.5 | 81.8076923076923 |
| RLG | RLG00174 | 0 | 21 | 91 | 173.421052631579 |
| RLG | RLG00181 | 0 | 22 | 304.5 | 353.272727272727 |
| RLG | RLG00186 | 0 | 21 | 254.5 | 308 |

| | | | | | |
|---|---|---|---|---|---|
| RLG | RLG00191 | 0 | 19 | 225 | 279.157894736842 |
| RLG | RLG00192 | 0 | 20 | 402 | 396.1 |
| RLG | RLG00193 | 0 | 17 | 275 | 318 |
| RLG | RLG00223 | 0 | 15 | 292 | 328.8 |
| RLG | RLG00226 | 0 | 15 | 278 | 279.066666666667 |
| RLG | RLG00227 | 0 | 14 | 264.5 | 300.357142857143 |
| RLG | RLG00237 | 0 | 10 | 231 | 305.1 |
| RLG | RLG00239 | 0 | 16 | 320 | 316.4375 |
| RLG | RLG00241 | 0 | 15 | 247 | 262.733333333333 |
| RLG | RLG00242 | 0 | 15 | 272 | 337.933333333333 |
| RLG | RLG00251 | 0 | 15 | 276 | 347 |
| RLG | RLG00261 | 0 | 14 | 77 | 186.142857142857 |
| RLG | RLG00263 | 0 | 14 | 438 | 446.571428571429 |
| RLG | RLG00270 | 0 | 11 | 293.5 | 284.5 |
| RLG | RLG00275 | 0 | 11 | 67 | 72.2727272727273 |
| RLG | RLG00283 | 0 | 13 | 361 | 597.538461538462 |
| RLG | RLG00289 | 0 | 11 | 809 | 752.727272727273 |
| RLG | RLG00306 | 0 | 11 | 238 | 300.454545454545 |
| RLG | RLG00323 | 0 | 12 | 349.5 | 362 |
| RLG | RLG00326 | 0 | 11 | 638 | 638 |
| RLG | RLG00327 | 0 | 13 | 99 | 147 |
| RLG | RLG00330 | 0 | 10 | 209.5 | 270.1 |
| RLG | RLG00331 | 0 | 11 | 292 | 345.090909090909 |
| RLG | RLG00334 | 0 | 11 | 564 | 557.636363636364 |
| RLG | RLG00342 | 0 | 11 | 302 | 329.363636363636 |
| RLG | RLG00357 | 0 | 10 | 167.5 | 184.2 |
| RLG | RLG00376 | 0 | 12 | 345 | 368.583333333333 |
| RLG | RLG00382 | 0 | 15 | 163 | 369.866666666667 |
| RLG | RLG00404 | 0 | 51 | 518 | 560.764705882353 |
| RLG | RLG00417 | 0 | 10 | 341 | 353.2 |
| RLG | RLG00620 | 0 | 198 | 551 | 590.89898989899 |
| RLG | RLG01210 | 0 | 250 | 437 | 497.224 |
| RLG | RLG01224 | 0 | 18 | 356.5 | 421.944444444444 |
| RLG | RLG01297 | 0 | 14 | 75 | 169.307692307692 |
| RLG | RLG01328 | 0 | 36 | 399 | 475.833333333333 |
| RLG | RLG01860 | 0 | 18 | 227 | 240.833333333333 |
| RLG | RLG02129 | 0 | 60 | 634 | 683.466666666667 |
| RLG | RLG02424 | 0 | 21 | 242 | 259.047619047619 |
| RLG | RLG02722 | 0 | 131 | 381 | 397.885496183206 |
| RLG | RLG04884 | 0 | 82 | 577.5 | 526.963414634146 |
| RLG | RLG07846 | 0 | 156 | 628 | 575.532051282051 |
| RLG | RLG07969 | 0 | 18 | 326 | 348.888888888889 |
| RLG | RLG16490 | 0 | 27 | 478 | 483.814814814815 |
| RLG | RLG20706 | 0 | 34 | 72 | 79.0333333333333 |
| RLG | RLG21631 | 0 | 68 | 291 | 275.602941176471 |
| RLG | RLG22910 | 0 | 65 | 317 | 311.138461538462 |
| RLG | RLG23054 | 0 | 44 | 969.5 | 958.363636363636 |
| RLG | RLG23897 | 0 | 11 | 192 | 197.636363636364 |
| RLX | RLX00065 | 0 | 83 | 264.5 | 250.175 |
| RLX | RLX00066 | 0 | 85 | 76 | 82.4939759036145 |
| RLX | RLX00092 | 0 | 63 | 95 | 114.983870967742 |
| RLX | RLX00094 | 0 | 47 | 306 | 252.652173913043 |
| RLX | RLX00095 | 0 | 44 | 613 | 517.790697674419 |
| RLX | RLX00134 | 0 | 29 | 352 | 493.047619047619 |
| RLX | RLX00144 | 0 | 27 | 238 | 318.740740740741 |
| RLX | RLX00168 | 0 | 24 | 303 | 254.04347826087 |
| RLX | RLX00171 | 0 | 27 | 623 | 534 |

| | | | | | |
|---|---|---|---|---|---|
| RLX | RLX00172 | 0 | 23 | 243 | 232.47619047619 |
| RLX | RLX00196 | 0 | 19 | 56 | 97.2105263157895 |
| RLX | RLX00197 | 0 | 18 | 54.5 | 102 |
| RLX | RLX00210 | 0 | 18 | 138.5 | 154.666666666667 |
| RLX | RLX00219 | 0 | 19 | 93 | 111.368421052632 |
| RLX | RLX00220 | 0 | 17 | 220 | 229.294117647059 |
| RLX | RLX00230 | 0 | 16 | 86 | 104 |
| RLX | RLX00234 | 0 | 16 | 72.5 | 101.3125 |
| RLX | RLX00245 | 0 | 16 | 87 | 113.0625 |
| RLX | RLX00264 | 0 | 14 | 103 | 178.153846153846 |
| RLX | RLX00299 | 0 | 12 | 63 | 86.5 |
| RLX | RLX00301 | 0 | 12 | 27 | 27 |
| RLX | RLX00315 | 0 | 13 | 177 | 175.076923076923 |
| RLX | RLX00317 | 0 | 12 | 198.5 | 250.666666666667 |
| RLX | RLX00349 | 0 | 10 | 59 | 69.3333333333333 |
| RLX | RLX00360 | 0 | 10 | 229.5 | 298.6 |
| RLX | RLX00369 | 0 | 10 | 178 | 154.5 |
| RLX | RLX00370 | 0 | 10 | 295 | 275.5 |
| RLX | RLX00560 | 0 | 14 | 249 | 247.428571428571 |
| RLX | RLX00647 | 0 | 12 | 765.5 | 814.916666666667 |
| RLX | RLX00824 | 0 | 50 | 126 | 165.615384615385 |
| RLX | RLX00910 | 0 | 13 | 536 | 591.692307692308 |
| RLX | RLX01857 | 0 | 36 | 641 | 677.944444444444 |
| RLX | RLX01925 | 0 | 14 | 569 | 563.615384615385 |
| RLX | RLX02412 | 0 | 23 | 897 | 832.521739130435 |
| RLX | RLX02653 | 0 | 15 | 320 | 341.733333333333 |
| RLX | RLX02676 | 0 | 75 | 690 | 653.813333333333 |
| RLX | RLX03238 | 0 | 13 | 252 | 206.769230769231 |
| RLX | RLX03863 | 0 | 183 | 943 | 886.07650273224 |
| RLX | RLX04027 | 0 | 32 | 333 | 300.15625 |
| RLX | RLX06052 | 0 | 25 | 760 | 760.84 |
| RLX | RLX06850 | 0 | 10 | 505 | 532 |
| RLX | RLX06925 | 0 | 10 | 416.5 | 358.8 |
| RLX | RLX07110 | 0 | 22 | 456.5 | 459 |
| RLX | RLX07511 | 0 | 41 | 60 | 78.4878048780488 |
| RLX | RLX08341 | 0 | 13 | 265 | 307.769230769231 |
| RLX | RLX08581 | 0 | 28 | 156 | 160.607142857143 |
| RLX | RLX09595 | 0 | 22 | 468.5 | 447.363636363636 |
| RLX | RLX10780 | 0 | 13 | 188 | 180.307692307692 |
| RLX | RLX10916 | 0 | 64 | 756 | 715.859375 |
| RLX | RLX11328 | 0 | 24 | 524 | 484.333333333333 |
| RLX | RLX11700 | 0 | 11 | 212 | 231.727272727273 |
| RLX | RLX12104 | 0 | 32 | 298.5 | 355.03125 |
| RLX | RLX12410 | 0 | 10 | 246 | 284.9 |
| RLX | RLX12632 | 0 | 10 | 34 | 49.3 |
| RLX | RLX12917 | 0 | 43 | 622 | 542.46511627907 |
| RLX | RLX13049 | 0 | 334 | 688 | 673.446107784431 |
| RLX | RLX13352 | 0 | 13 | 364 | 399.461538461538 |
| RLX | RLX13514 | 0 | 153 | 819 | 840.843137254902 |
| RLX | RLX13529 | 0 | 185 | 252 | 230.918918918919 |
| RLX | RLX13698 | 0 | 105 | 627 | 565.857142857143 |
| RLX | RLX13810 | 0 | 12 | 490.5 | 483.333333333333 |
| RLX | RLX14021 | 0 | 31 | 458 | 624.032258064516 |
| RLX | RLX14124 | 0 | 28 | 406.5 | 438.357142857143 |
| RLX | RLX14762 | 0 | 11 | 442 | 423.909090909091 |
| RLX | RLX16782 | 0 | 168 | 652 | 591.690476190476 |
| RLX | RLX17675 | 0 | 105 | 627 | 665.959595959596 |

| | | | | | |
|---|---|---|---|---|---|
| RLX | RLX19305 | 0 | 13 | 150 | 155.846153846154 |
| RLX | RLX19534 | 0 | 21 | 67.5 | 67.2777777777778 |
| RLX | RLX19697 | 0 | 13 | 372 | 406.615384615385 |
| RLX | RLX19762 | 0 | 28 | 278.5 | 319.107142857143 |
| RLX | RLX22845 | 0 | 15 | 358 | 355.933333333333 |
| RLX | RLX23205 | 0 | 16 | 294.5 | 314.625 |
| RLX | RLX23786 | 0 | 41 | 44 | 51.2564102564103 |
| RST | RST00001 | 0 | 22 | 29 | 29.25 |
| RST | RST00003 | 0 | 17 | 169 | 149.352941176471 |
| RST | RST00004 | 0 | 18 | 126 | 110.0625 |
| RST | RST00005 | 0 | 38 | 50 | 48.8888888888889 |
| RST | RST00008 | 0 | 16 | 97 | 100.375 |
| RST | RST00010 | 0 | 25 | NA | NaN |
| RST | RST00014 | 0 | 41 | 54 | 55.6666666666667 |
| RST | RST00016 | 0 | 32 | 27 | 31.6666666666667 |
| RST | RST00019 | 0 | 15 | 43 | 43 |

**Table S4.** Mean methylation levels across superfamilies, averaged across all tissues, and averaged within a tissue (all3=seedling)

| sup | avg_cg | avg_chg | avg_chh | SAM_avg_cg | SAM_avg_chg | SAM_avg_chh | all3_avg_cg | all3_avg_chg | all3_avg_chh | flagleaf_avg_cg | flagleaf_avg_chg | flagleaf_avg_chh | earshoot_avg_cg | earshoot_avg_chg | earshoot_avg_chh | anther_avg_cg | anther_avg_chg | anther_avg_chh |
|---|---|---|---|---|---|---|---|---|---|---|---|---|---|---|---|---|---|---|
| DHH | 0.768 | 0.659 | 0.034 | 0.741 | 0.606 | 0.025 | 0.743 | 0.743 | 0.050 | 0.806 | 0.686 | 0.036 | 0.778 | 0.655 | 0.028 | 0.773 | 0.604 | 0.028 |
| DTA | 0.804 | 0.607 | 0.082 | 0.774 | 0.519 | 0.060 | 0.782 | 0.782 | 0.101 | 0.840 | 0.637 | 0.114 | 0.816 | 0.575 | 0.067 | 0.807 | 0.524 | 0.068 |
| DTC | 0.833 | 0.693 | 0.046 | 0.799 | 0.625 | 0.034 | 0.811 | 0.811 | 0.061 | 0.875 | 0.726 | 0.052 | 0.843 | 0.686 | 0.041 | 0.837 | 0.617 | 0.040 |
| DTH | 0.811 | 0.681 | 0.062 | 0.781 | 0.616 | 0.046 | 0.794 | 0.794 | 0.085 | 0.847 | 0.712 | 0.079 | 0.822 | 0.670 | 0.047 | 0.812 | 0.611 | 0.052 |
| DTM | 0.738 | 0.595 | 0.144 | 0.713 | 0.525 | 0.095 | 0.708 | 0.708 | 0.199 | 0.770 | 0.623 | 0.195 | 0.758 | 0.592 | 0.104 | 0.741 | 0.529 | 0.128 |
| DTT | 0.801 | 0.674 | 0.044 | 0.773 | 0.617 | 0.032 | 0.775 | 0.775 | 0.068 | 0.839 | 0.702 | 0.050 | 0.813 | 0.671 | 0.034 | 0.803 | 0.602 | 0.034 |
| DTX | 0.848 | 0.732 | 0.041 | 0.816 | 0.670 | 0.033 | 0.838 | 0.838 | 0.060 | 0.883 | 0.766 | 0.042 | 0.854 | 0.725 | 0.034 | 0.851 | 0.663 | 0.034 |
| RIL | 0.837 | 0.687 | 0.025 | 0.809 | 0.617 | 0.019 | 0.801 | 0.801 | 0.037 | 0.879 | 0.700 | 0.026 | 0.850 | 0.677 | 0.021 | 0.844 | 0.639 | 0.019 |
| RIT | 0.850 | 0.725 | 0.032 | 0.822 | 0.653 | 0.023 | 0.824 | 0.824 | 0.043 | 0.891 | 0.767 | 0.043 | 0.861 | 0.714 | 0.027 | 0.854 | 0.669 | 0.024 |
| RLC | 0.824 | 0.677 | 0.027 | 0.794 | 0.617 | 0.021 | 0.797 | 0.797 | 0.047 | 0.868 | 0.705 | 0.027 | 0.834 | 0.668 | 0.022 | 0.827 | 0.597 | 0.020 |
| RLG | 0.842 | 0.735 | 0.029 | 0.812 | 0.681 | 0.024 | 0.822 | 0.822 | 0.043 | 0.879 | 0.762 | 0.028 | 0.850 | 0.736 | 0.025 | 0.845 | 0.673 | 0.024 |
| RLX | 0.806 | 0.695 | 0.028 | 0.776 | 0.642 | 0.022 | 0.784 | 0.784 | 0.044 | 0.844 | 0.721 | 0.028 | 0.815 | 0.696 | 0.024 | 0.808 | 0.633 | 0.023 |
| RST | 0.740 | 0.570 | 0.035 | 0.716 | 0.510 | 0.027 | 0.716 | 0.716 | 0.053 | 0.773 | 0.574 | 0.040 | 0.755 | 0.553 | 0.027 | 0.742 | 0.497 | 0.027 |

**Table S5.** TE families that lack methylatable cytosines (presented as family median values).

| sup | fam | famsize | tebp | percGC | nCG | nCHG | nCHH |
|---|---|---|---|---|---|---|---|
| DTH | DTH00067 | 602 | 196 | 0.318 | 0 | 0.009 | 0.139 |
| DTH | DTH00073 | 122 | 127 | 0.441 | 0 | 0 | 0.201 |
| DTH | DTH00090 | 214 | 127 | 0.302 | 0 | 0.016 | 0.120 |
| DTH | DTH00119 | 124 | 138 | 0.388 | 0 | 0 | 0.173 |
| DTH | DTH00163 | 968 | 116 | 0.350 | 0 | 0 | 0.151 |
| DTH | DTH00280 | 289 | 200 | 0.440 | 0 | 0.105 | 0.105 |
| DTH | DTH00320 | 95 | 248 | 0.330 | 0 | 0.002 | 0.154 |
| DTH | DTH00409 | 748 | 178 | 0.296 | 0 | 0 | 0.132 |
| DTH | DTH00410 | 76 | 147 | 0.262 | 0 | 0.015 | 0.105 |
| DTH | DTH00460 | 81 | 116 | 0.364 | 0 | 0 | 0.155 |
| DTH | DTH00486 | 16 | 138 | 0.415 | 0 | 0 | 0.187 |
| DTH | DTH10176 | 146 | 160 | 0.371 | 0 | 0 | 0.175 |
| DTH | DTH10226 | 15 | 130 | 0.354 | 0.008 | 0 | 0.165 |
| DTH | DTH10700 | 368 | 103 | 0.386 | 0 | 0 | 0.190 |
| DTH | DTH10730 | 3,113 | 67 | 0.412 | 0 | 0 | 0.201 |
| DTH | DTH10855 | 659 | 88 | 0.307 | 0 | 0.011 | 0.142 |
| DTH | DTH10856 | 126 | 88 | 0.308 | 0 | 0.011 | 0.142 |
| DTH | DTH10946 | 179 | 87 | 0.483 | 0 | 0.141 | 0.103 |
| DTH | DTH11101 | 409 | 67 | 0.409 | 0 | 0 | 0.205 |
| DTH | DTH11481 | 35 | 52 | 0.347 | 0 | 0.008 | 0.159 |
| DTH | DTH11817 | 346 | 208 | 0.382 | 0 | 0.034 | 0.156 |
| DTH | DTH12058 | 11 | 160 | 0.373 | 0 | 0.006 | 0.169 |
| DTH | DTH12181 | 158 | 354 | 0.286 | 0 | 0.011 | 0.129 |
| DTH | DTH14130 | 11 | 97 | 0.289 | 0 | 0 | 0.127 |
| DTH | DTH15065 | 78 | 275.500 | 0.244 | 0 | 0.001 | 0.116 |
| DTH | DTH15139 | 12 | 358 | 0.375 | 0 | 0.028 | 0.157 |
| DTM | DTM00257 | 376 | 155 | 0.374 | 0 | 0 | 0.174 |
| DTT | DTT10009 | 702 | 147 | 0.283 | 0 | 0.010 | 0.125 |
| DTT | DTT10797 | 889 | 89 | 0.299 | 0 | 0 | 0.146 |
| DTT | DTT10880 | 6,151 | 156 | 0.357 | 0 | 0 | 0.174 |
| DTT | DTT10881 | 2,514 | 156 | 0.357 | 0 | 0 | 0.175 |
| DTT | DTT10986 | 20 | 120 | 0.425 | 0 | 0 | 0.194 |
| DTT | DTT11073 | 199 | 143 | 0.245 | 0 | 0 | 0.108 |
| DTT | DTT11169 | 39 | 108 | 0.290 | 0 | 0.008 | 0.132 |
| DTT | DTT11408 | 48 | 109 | 0.385 | 0 | 0 | 0.183 |
| DTT | DTT11547 | 1,825 | 156 | 0.359 | 0 | 0.006 | 0.171 |
| DTT | DTT12206 | 22 | 190.500 | 0.229 | 0 | 0.011 | 0.097 |
| DTT | DTT14220 | 57 | 157 | 0.282 | 0 | 0.010 | 0.116 |
| DTT | DTT15443 | 143 | 156 | 0.355 | 0 | 0.005 | 0.168 |
| DTT | DTT15444 | 519 | 156 | 0.353 | 0 | 0 | 0.174 |
| DTX | DTX10131 | 120 | 141 | 0.255 | 0 | 0.013 | 0.110 |
| DTX | DTX10132 | 90 | 131.500 | 0.246 | 0 | 0.014 | 0.105 |
| DTX | DTX10177 | 2,850 | 156 | 0.357 | 0 | 0 | 0.176 |
| DTX | DTX10212 | 101 | 138 | 0.391 | 0 | 0 | 0.181 |
| DTX | DTX10213 | 15 | 138 | 0.406 | 0 | 0 | 0.188 |
| DTX | DTX10945 | 10 | 87 | 0.451 | 0 | 0.126 | 0.097 |
| DTX | DTX11178 | 18 | 72 | 0.458 | 0 | 0 | 0.212 |
| DTX | DTX12196 | 26 | 165 | 0.273 | 0 | 0.024 | 0.105 |
| DTX | DTX12438 | 23 | 275 | 0.353 | 0 | 0.004 | 0.162 |
| RST | RST00001 | 22 | 140 | 0.349 | 0.014 | 0 | 0.153 |
| RST | RST00010 | 25 | 309 | 0.298 | 0 | 0.016 | 0.132 |
